# Unexpected contributions of striatal projection neurons coexpressing dopamine D1 and D2 receptors in balancing motor control

**DOI:** 10.1101/2022.04.05.487163

**Authors:** Patricia Bonnavion, Christophe Varin, Ghazal Fakhfouri, Pilar Martinez Olondo, Aurélie De Groote, Amandine Cornil, Ramiro Lorenzo Lopez, Elisa Pozuelo Fernandez, Elsa Isingrini, Quentin Rainer, Kathleen Xu, Eleni Tzavara, Erika Vigneault, Sylvie Dumas, Alban de Kerchove d’Exaerde, Bruno Giros

## Abstract

The central function of the striatum and its dopaminergic (DA) afferents in motor control and the integration of cognitive and emotional processes is commonly explained by the two striatal efferent pathways characterized by striatal projection neurons (SPNs) expressing DA D1 receptors and D2 receptors (D1-SPNs and D2-SPNs), without regard to SPNs coexpressing both receptors (D1/D2-SPNs). We developed an approach that enables the targeting of these hybrid SPNs and demonstrated that although these SPNs are less abundant, they play a major role in guiding the motor function of the other two main populations. D1/D2-SPNs project exclusively to the external globus pallidus (GPe) and have specific electrophysiological features with distinctive integration of DA signals. Optogenetic stimulation and loss-of-function experiments indicated that D1/D2-SPNs potentiate the prokinetic and antikinetic functions of D1-SPNs and D2-SPNs, respectively, and restrain the integrated motor response to psychostimulants. Overall, our findings demonstrate the essential role of this third unacknowledged population of D1/D2 coexpressing neurons, which orchestrates the fine-tuning of DA regulation in the thalamo-cortico-striatal loops.

**One-Sentence Summary:** D1/D2 SPNs modulate the motor function of both D1- and D2-SPNs

## Main text

The thalamo-cortico-striatal loops are essential cerebral modules for the initiation, integration, and regulation of motor, cognitive, and emotional processes^1–4^. The functional specificity and precision of their actions are finely tuned through the complex design of striatal organization^5–7^. Striatal control of cortical excitation and inhibition requires two distinct output pathways^8, 9^: a direct pathway (cortex→striatum→GPi/SNr) facilitates motor cortical activity, whereas an indirect pathway (cortex→striatum→GPe→STN→GPi/SNr) inhibits the motor cortex. The direct pathway involves striatal projection neurons (SPNs) expressing the DA D1 receptor (D1R), while the indirect pathway involves SPNs expressing the dopamine D2 receptor (D2R)^10^. This organization substantiates why DA is such an essential neurotransmitter in movement and motor control^11^ but also in the reward system and cognitive processes^12^. Because DA likely stimulates D1-SPNs in the direct pathway and inhibits D2-SPNs in the indirect pathway, it is often postulated that DA has a synergistic effect promoting locomotion^6^. Supporting this general description, the motor response to cocaine administration is suppressed or blunted in D1R-KO^13^ and SPN-D2R KO^14^, respectively. Despite these effects supporting the classical model attributing a unique and homogeneous role to DA in the striatum, growing evidence indicates several levels of heterogeneity within the rodent striatal system that still poorly integrate the striatal actions of DA under physiological conditions. For example, in contrast to the classical model suggesting opposite motor roles of the two SPN populations, it has been shown that optogenetic activation of D1- and D2-SPNs produces different responses in SNr and cortical neurons, with stimulation of each pathway eliciting both excitation and inhibition^15–17^. Moreover, it has also been shown that both populations are jointly active during a motor task^18–21^. Finally, single-cell molecular analysis of striatal neurons reveals more complex heterogeneity and identifies several distinct clusters of SPNs^22–27^. Notably, in addition to the presence of D1-SPNs and D2-SPNs, the existence of hybrid SPNs coexpressing D1Rs and D2Rs has been described; these hybrid SPNs had previously been identified by indirect approaches^28–33^, but their role and functions remain unknown. Using targeted approaches, we demonstrate here that this population of hybrid D1/D2-SPNs gives rise to a third specific efferent pathway with critical functions for motor control and its DA-mediated modulation in an organized combination with the two canonical D1- and D2-SPN pathways.

### SPNs coexpressing D1/D2 constitute a specific population

Previous single-cell and transgenic studies have provided only indirect and sometimes controversial indications of D1/D2-SPN distribution^28–32^. Here, we used direct and highly sensitive double-probe *in situ* hybridization (sdFISH) throughout the striatum to demonstrate native coexpression of D1Rs and D2Rs. Unbiased automated quantitative analysis showed that D1R/D2R coexpressing cells accounted for 6-7% of all cells (expressing D1Rs and D2Rs) in both the dorsal and ventral parts of the striatum, across its rostro-caudal extent (**Fig. 1a-b, Extended Data Fig. 1a,b**).

**Fig. 1.**
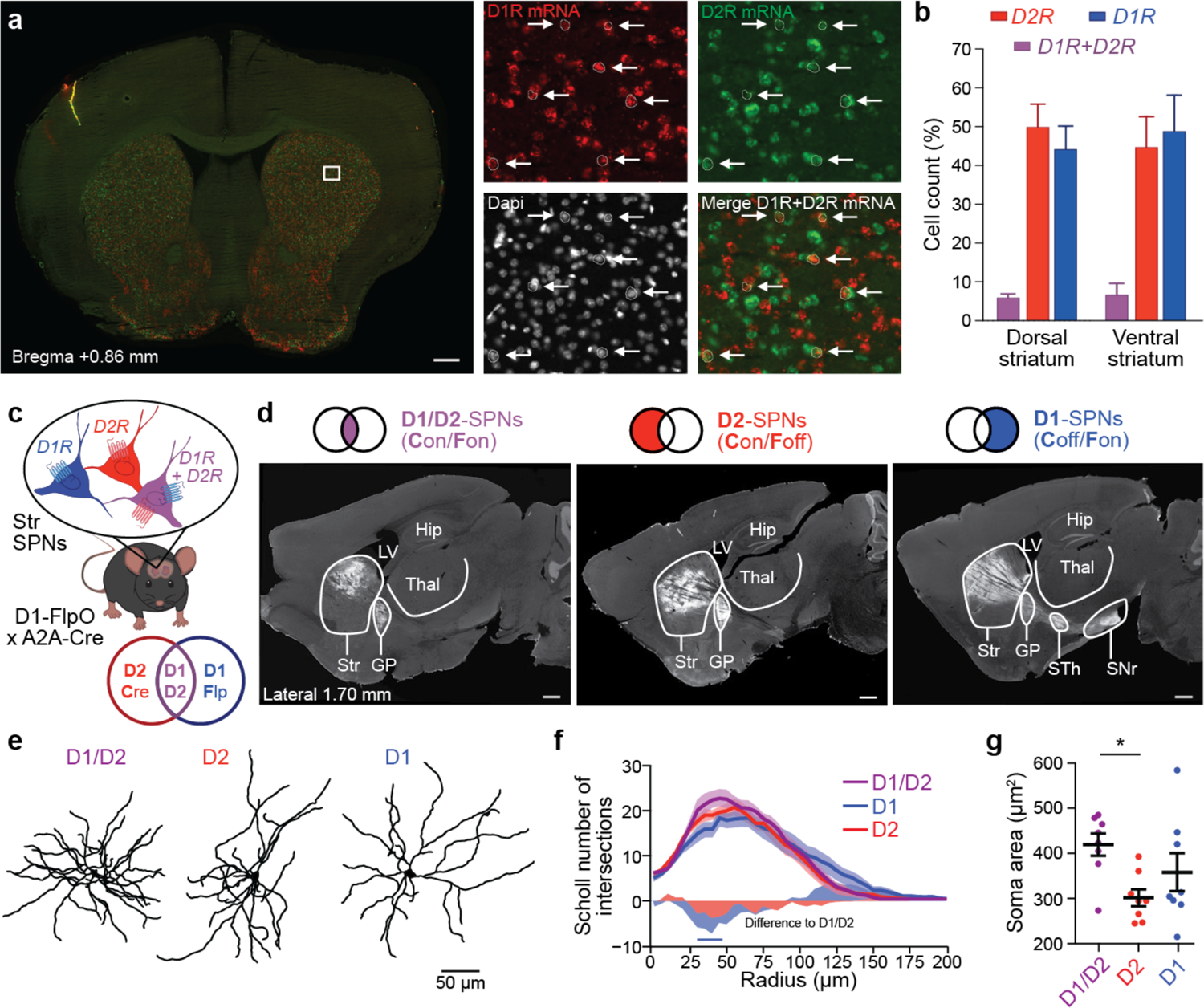
Morphologically distinct D1/D2 co-expressing SPNs establish an additional efferent striatal pathway. **a**, Representative coronal section of fluorescent in situ hybridization illustrating D1/D2 co-expression (arrows) in the striatum (scale: left, 1 mm; zooms, 20 µm). **b**, Percentage of D1-, D2-, and D1/D2-positive neurons over cells expressing either D1 or D2 in the dorsal and ventral striatum (n = 3 mice). **c**, Schematic illustrating INTRSECT targeting strategy applied to D1^FlpO^/A2a^Cre^ mice. **d**, Representative parasagittal sections illustrating projection patterns of D1-, D2-, and D1/D2-SPNs obtained from D1^FlpO^/A2a^Cre^ mice injected with respective INTRSECT AAV constructs (scale: 500 µm). Abbreviations: Str, striatum; GP, globus pallidus; STh, subthalamic nucleus; SNr, substantia nigra pars reticulate; Thal, thalamus, Hip, hippocampus; LV, lateral ventricle. **(e)** Reconstruction of the dendritic arborisation of representative D1-SPN (left), D2-SPN (middle), and D1/D2-SPN (left). **f-g**, Sholl analysis (**f**) and soma area (**g**) of reconstructed SPNs (D1-SPNs, n = 8; D2-SPNs, n = 8; D1/D2-SPNs, n = 8) (in f, horizontal blue line indicates t-test p < 0.05 between D1/D2 and D1; in g, Tukey tests: *p < 0.05).

To selectively target this population of SPNs coexpressing D1R and D2R, we used an intersectional approach^34^ (**Fig. 1c**) by generating a novel D1^FlpO^ driver line expressing Flp recombinase (FlpO) in D1-SPNs (**Extended Data Fig. 1c-d**). D1^FlpO^ were crossed with the A2a^Cre^ driver line expressing Cre recombinase exclusively in D2-SPNs, thereby excluding cholinergic interneurons^35^ (**Extended Data Fig. 1e**). Using this approach, we were able to individually target SPNs expressing both D1Rs and D2Rs (D1/D2-SPNs) for the first time and to access and control SPNs exclusively expressing D1Rs (D1-SPNs) and SPNs exclusively expressing D2Rs (D2-SPNs). This relies upon the presence of Cre (C) and/or FlpO (F): **C**on/**F**on, **C**off/**F**on and **C**on/**F**off (**Fig. 1c, Extended Data Fig. 1f**). We first used eYFP expression with INTRSECT vectors to validate the proper recombination of the constructs in mice expressing both Cre and FlpO and in mice expressing only Cre or FlpO (Extended Data Fig. 1f). No leaking recombination of the Con/Fon, Coff/Fon, and Con/Foff vectors was observed. We subsequently quantified the cell densities observed for each transfected SPN subpopulation (Extended Data Fig 1f, g). With this approach, we found proportions of 16.6 ± 2.4% for D1/D2-SPNs, 45.2 ± 5.2% for D2-SPNs, and 38.2 ± 0.7% for D1-SPNs relative to total SPNs. These proportions, higher than those obtained using sdFISH, are most likely a result of underdetection using sdFISH. Next, we similarly expressed channel rhodopsin 2 (ChR2) fused to eYFP to trace neuronal pathways (Fig. 1d; Extended Data Fig. 1f). We found that D1/D2-SPNs have a single projection pathway to the external globus pallidus (GPe), dissociating them from a direct pathway to the internal globus pallidus (GPi)/substantia nigra pars reticulata (SNr). This mapping highlights that D1/D2-SPNs give rise to an additional indirect pathway, which may therefore propagate a different DA-mediated signal to downstream basal ganglia nuclei. Furthermore, concerning their cell morphology, this population is distinguished by a more ramified proximal arborization than D1-SPNs and a larger soma area than D2-SPNs (**Fig. 1e-g**), suggesting that these D1/D2 coexpressing neurons may have distinct physiological properties and carry different functions than D1-SPNs and D2-SPNs.

### D1/D2-SPNs display a unique combination of electrophysiological features

Based on their morphological specificity, we hypothesized additional distinctions, and we examined the electrophysiological properties of D1/D2-SPNs in comparison with the two classical populations. Thirty electrophysiological parameters were extracted from whole-cell current clamp recordings describing passive membrane properties, firing properties, action potential properties and threshold properties (**Fig. 2a-c, Extended Data Table 1**). As previously reported^36–38^, D1- and D2-SPNs differed with respect to many features (**Fig. 2c-k, Extended Data Fig. 2a-c**). We found some mixed properties for the D1/D2-SPNs; the maximum intensity before the depolarization block and the current frequency early slope are similar to those of D1-SPNs (**Fig. 2h-i**), while the step intensity to the first action potential is similar to that of D2-SPNs (**Fig. 2g, Extended Data Table 1**). More essentially, D1/D2-SPNs exhibit a unique combination of action potential properties, such as a lower AP threshold and larger AP amplitude (**Fig. 2e, j-k, Extended Data Fig. 2**). Taken together, anatomical, morphological and electrophysiological results substantiate the reality of a third functional SPN population.

**Fig. 2.**
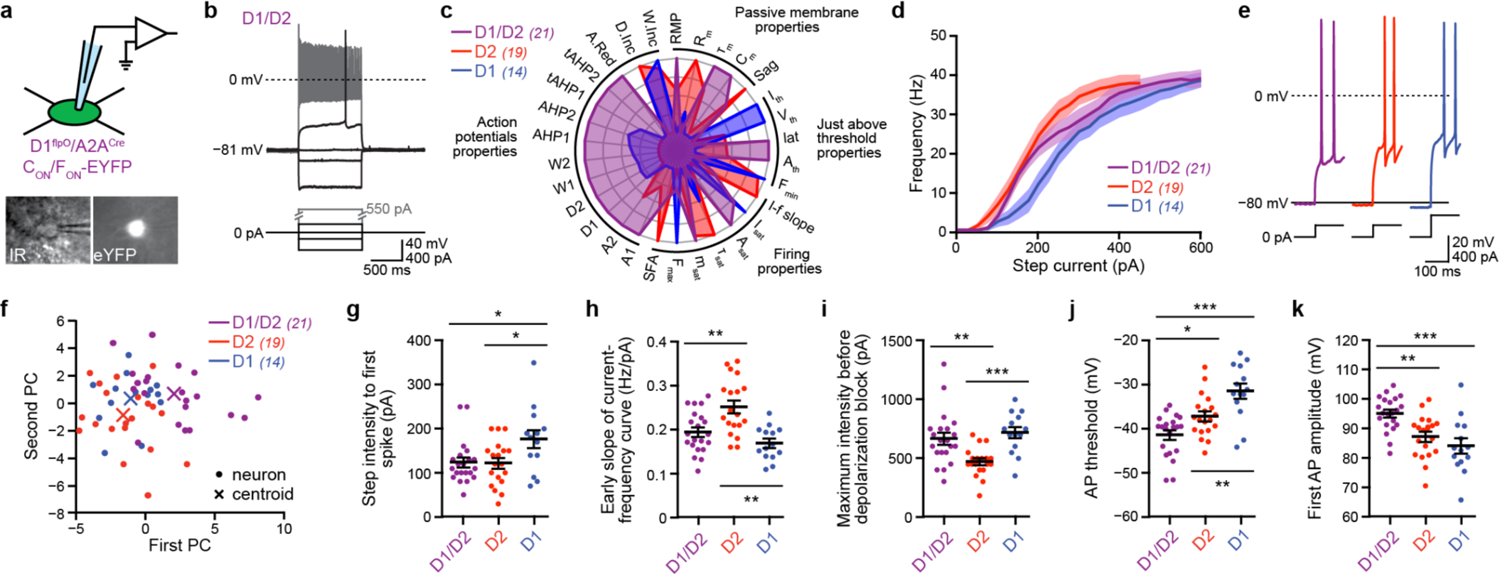
D1/D2-SPNs differ through unique action potential properties. **a**, Diagram of patch-clamp recordings in a YFP labeled D1/D2-SPN (top), and representative photomicrograph under infrared illumination (bottom left) and fluorescent illumination (bottom right). **b**, Typical voltage response of the same D1/D2-SPN as in **a** in response to hyperpolarizing and depolarizing current steps. **c**, Radar plot revealing differences between the three SPN types across all electrophysiological properties extracted. Means for each population are normalized in the range 0-1. **d**, Average current-frequency relationship for the three SPN subpopulations. **e**, Representative examples of action potential (AP) response to depolarizing current in one D1/D2-SPN, one D1-SPN and one D2-SPN. **f**, Two-dimensional projection of the set of neurons recorded (circles) and populations’ centroids (cross) following PCA into the first- and second-order components. **g-k**, Quantification of minimal step current to discharge (**g**), early slope of the current-frequency curve (**h**), current intensity to discharge saturation (**i**), AP threshold (**j**), and amplitude of the first AP (**k**) in D1/D2-SPNs (n = 21 cells from 9 mice), D1-(n = 14 cells from 5 mice) and D2-(n = 19 cells from 5 mice) (Tukey’s tests: *p < 0.05, **p < 0.01, ***p < 0.001). n = 11). **e**, Effect of bilateral optogenetic stimulation (10 x 30 s trials with 90 s between trials) on motor architecture in the open field evaluated during locomotion (top line), small movements (middle line), and immobility (bottom line) by the: time spent in state (left column), the number of episodes per minute in each state (middle column) and the mean duration of these episodes (right column) (Tukey’s test pre vs. light ON: *p < 0.05; **p < 0.01; ***p < 0.001). **f-h**, Effect of laser stimulation on each group on the averaged temporal evolutions of mice velocity (**f**) and acceleration (**g**) around locomotion onset and averaged velocity (black) and top velocity (9^th^ decile, green) during locomotion episodes (**h**).

### D1/D2-SPNs promote immobility and drive the prokinetic and antikinetic functions of D1- and D2-SPNs, respectively

To identify the motor function of this third SPN population, we performed *in vivo* optogenetic stimulations of D1/D2-SPNs in the dorsomedial part of the striatum (**Fig. 3**; **Extended Data Fig. 3**). To obtain additional information on their influence on coactivation with D2- or D1-SPNs, we studied six experimental groups, including exclusive targeting of ChR2 in D1/D2-SPNs (purple; **Fig. 3a-b**), D2-SPNs (red), and D1-SPNs (dark blue), as well as mixed targeting of ChR2 in which D2-SPNs are coactivated with D1/D2-SPNs (pink, D2+D1/D2-SPNs), or D1-SPNs are coactivated with D1/D2-SPNs (light blue, D1+D1/D2-SPNs), and finally D1/D2^eYFP^ control mice (gray). The conditions D2+D1/D2-SPNs and D1+D1/D2-SPNs correspond to what was defined as indirect pathway SPNs or direct pathway SPNs, respectively, in previous studies. Thus, this comprehensive approach also allows us to re-examine the respective roles of exclusive D2- and D1-SPNs.

**Fig. 3.**
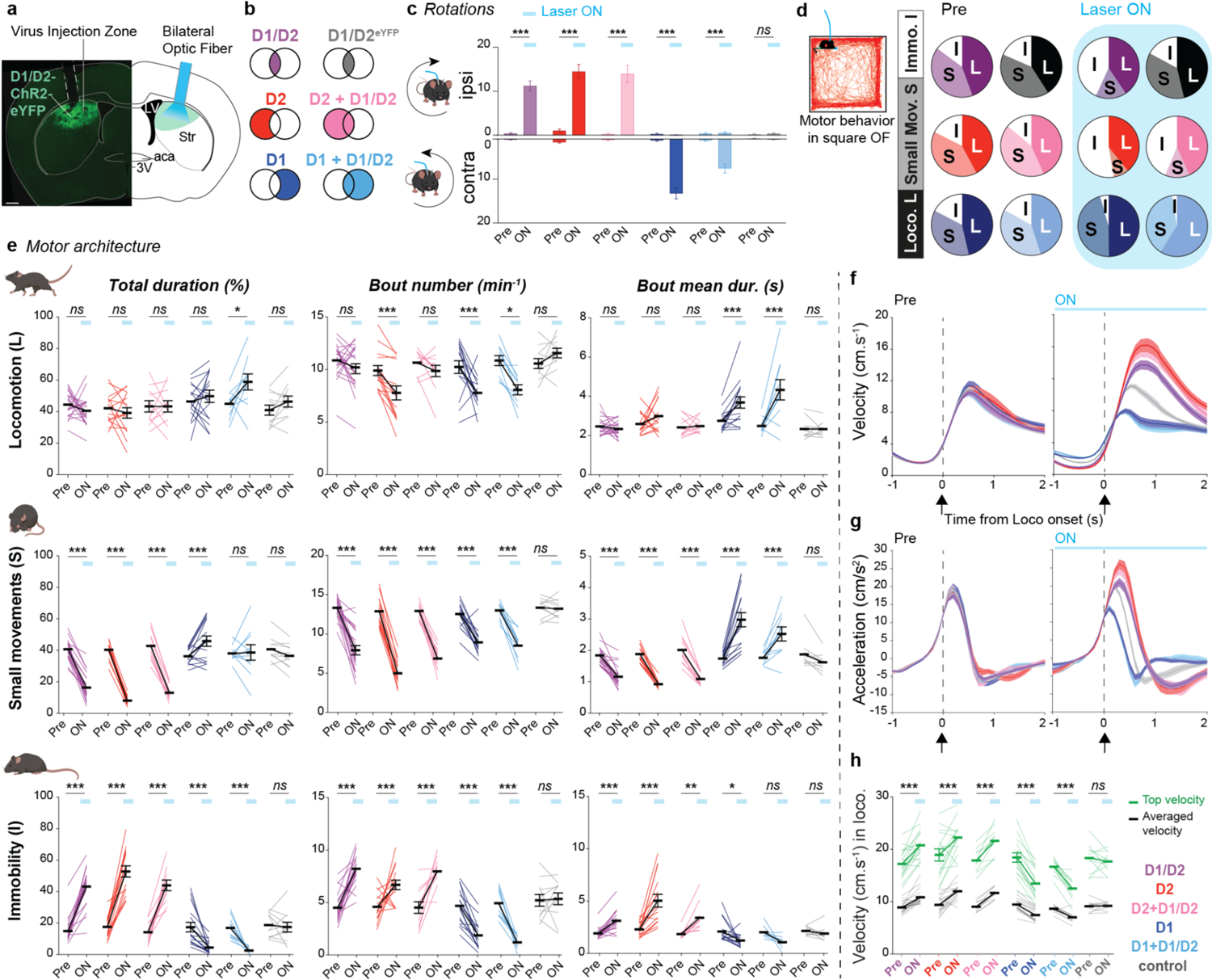
D1/D2-SPNs promote immobility and control the effects of D1- and D2-SPNs on locomotion. **a**, Coronal section and corresponding atlas plane depicting optic fibers placement targeting the dorsal striatum and ChR2 expression in D1/D2-SPNs in D1^FlpO^/A2a^Cre^ mice (scale: 500 μm). **b**, Schematic representation of the experimental groups. c, Rotational behavior elicited by unilateral continuous illumination (60 s) of ChR2-expressing D1/D2- (n = 21), D2- (n = 16), D2+D1/D2- (n = 13), D1- (n = 18), D1+D1/D2- (n = 16) and control D1/D2^eYFP^ (n = 11)-SPNs stimulated mice (Tukey’s test pre vs. light ON: ***p < 0.001). **d**, Distribution and proportions of the three motor states defined in free-running mice exploring an open field during baseline (Pre) and stimulation (Laser ON) periods: D1/D2-SPNs Pre: L: 44.4±1.7% | S: 40.7±1.4% | I: 14.9±1.7%; Laser ON: L: 40.5±2.0% | S: 16.3±1.8% | I: 43.2±2.6%; D2-SPNs Pre: L: 42.2±2.5% | S: 40.3±1.3% | I: 17.6±1.8%; Laser ON: L: 39.1±3.4% | S: 8.1±0.8% | I: 52.8±3.6%; D2+D1.D2-SPNs Pre: L: 43.3±3.6% | S: 42.8±2.7% | I: 13.9±2.1%; Laser ON: L: 43.3±3.6% | S: 13.0±1.5% | I: 43.7±3.8%; D1-SPNs Pre: L: 46.4±2.8% | S: 36.2±1.2% | I: 17.4±3.1%; Laser ON: L: 49.8±3.9% | S: 45.9±3.3% | I: 4.4±1.0; D1+D1/D2-SPNs Pre: L: 45.0±2.8% | S: 38.1±1.6% | I: 16.9±2.2%; Laser ON: L: 58.9±5.1% | S: 38.6±4.9% | I: 2.5±0.5; control D1/D2eYFP-SPNs Pre: L: 40.9±3.2% | S: 40.7±2.3% | I: 18.4±2.4%; Laser ON: L: 46.3±3.5% | S: 36.5±2.3% | I: 17.1±3.1%D2-SPNs Pre: L: 44.7±4.0% | S: 39.1±2.0% | I: 16.2±3.2%; Laser ON: L: 46.6±2.4% | S: 8.5±1.3% | I: 44.9±2.9%. (D1/D2: n = 21, D2: n = 15, D2+D1/D2: n = 11, D1: n = 16, D1+D1/D2: n = 10, control D1/D2^eYFP^: 10 mg/kg cocaine injections between D1^cKO^/D2 and D1^wt^/D2 mice on motor activity during 30 min of open field (effect of days for each genotype: ^✝✝✝^p < 0.001, ns not significant; post-hoc vs. day1: *p < 0.05, **p < 0.01; ANOVA interaction days-genotypes: ^###^p < 0.001). **f**, Conditioned place preference for cocaine (10 mg/kg) in D1^wt^/D2 and D1^cKO^/D2 mice (Sidak’s tests between genotypes: ns not significant).

Unilateral activation of the indirect pathway has been shown to mimic ipsiversive rotational behavior, whereas activation of the direct pathway results in contraversive rotations^39^. Consistent with their GPe-targeted projection pattern, unilateral stimulation of D1/D2-SPNs also results in ipsilateral rotations, similar to that of D2-SPNs or D2+D1/D2-SPNs, while stimulation of D1-SPNs or D1+D1/D2-SPNs results in the opposite rotatory behavior (**Fig. 3c**). These effects directly validate that our intersectional optogenetic strategy is effective in all three populations.

We next examined the impact of bilateral stimulation on free-running mice in a square open field (**Fig. 3d**). A comparative analysis of three different stimulation patterns with pulsed light and a constant 30-s illumination (**Extended Data Fig. 3-4**, Supplementary text) revealed that the most reliable optogenetic paradigm is a constant illumination, which allows for sustained neuronal depolarization with SPNs firing at their own pace (**Extended Data Fig. 3f-i**, Supplementary text). Motor behavior was classified into three states: locomotion, small movements and immobility (**Fig. 3d**). On average, WT mice spent 44 ± 0.7% of their time in locomotion, 40 ± 0.8% in small movements and 16 ± 0.6% in immobility (**Fig. 3d**). Optogenetic activation of each SPN population significantly reorganizes this motor behavior distribution (**Fig. 3d-e**). Remarkably, the activation of D1/D2-SPNs, forming a smaller population, leads to a strong increase in immobility coupled with a marked inhibition of small movements. This pattern was the same as that observed in response to stimulation of the indirect pathway by exclusive activation of D2-SPNs or mixed coactivation of D2+D1/D2-SPNs (**Fig. 3d-e**). These stimulations had no significant effect on the total time spent in locomotion (**Fig. 3d-e**, Supplementary text). However, there is a notable difference in the microstructure of the locomotion state between exclusive stimulation of D2-SPNs and coactivation of D2+D1/D2-SPNs. Unexpectedly, stimulation of exclusively D2-SPNs seemed to consolidate locomotion by significantly reducing the number of locomotor episodes and nonsignificantly lengthening their mean duration (**Extended Data Fig. 5a**). In contrast, stimulation of D1/D2-SPNs or mixed D2+D1/D2-SPNs induces no change in these parameters, suggesting that the coactivation of D1/D2-SPNs can block some pro-locomotor D2-SPN-mediated effects that, alongside immobility facilitation and small movement inhibition, produce a clear antikinetic identity to the indirect pathway (Extended Data text). In addition, we observed significant differences between exclusive stimulation of D1-SPNs and coactivation of D1+D1/D2-SPNs. As previously reported^39, 40^, stimulation of mixed D1+D1/D2 SPNs induces a significant increase in time spent in locomotion by a lengthening of locomotor episodes and an almost complete abolition of the immobility state (**Fig. 3e, Extended Data Fig. 5a**). Instead of inducing a net augmentation effect on locomotion, stimulation of only D1-SPNs resulted in a large increase in the time spent in small movements at the expense of immobility (**Fig. 3e, Extended Data Fig. 5a**). This suggests that coactivation of D1/D2-SPNs with D1-SPNs allows for a net pro-locomotor effect of the direct pathway.

**Fig. 4.**
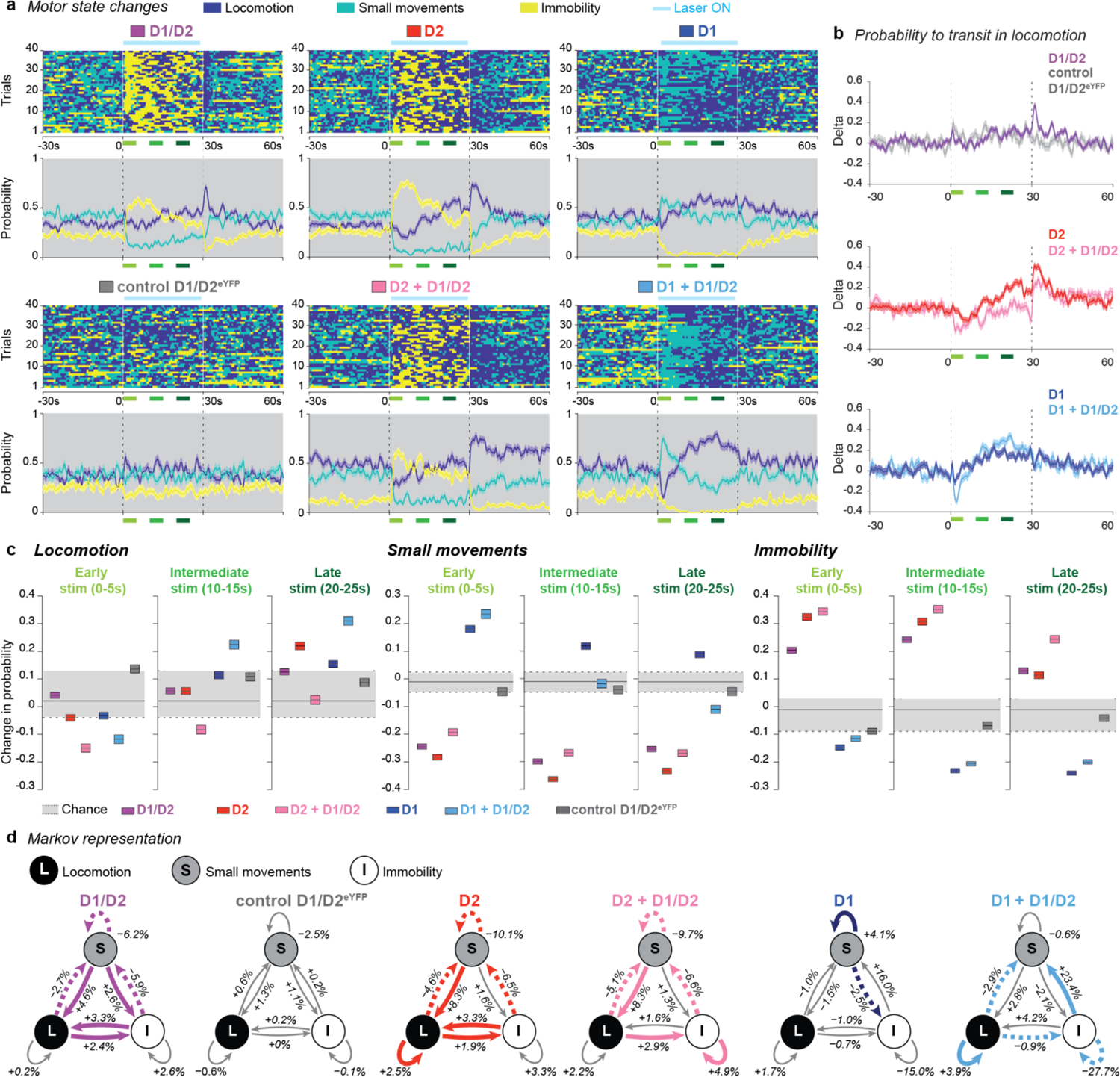
D1/D2-SPN coactivation modulates the temporal dynamics of motor changes elicited by D1- or D2-SPN stimulation. **a**, Motor states for 10 stimulation trials of 4 representative mice of each genotype (top) and corresponding temporal evolution of probability (bottom) to observe locomotion, small movements and immobility, before, during, and after laser stimulation (shading, bootstrap 95% CI). Blue bar, laser stimulation. **b**, Temporal evolution of changes in probabilities to observe locomotion centered on pre-stimulation values. **c**, Changes in occurrence probability of locomotion (left), small movements (middle), and immobility (right) induced by laser stimulation during the early (0-5 s; light green bars in a and b), intermediate (10-15 s; green bars in a and b), and late (20-25 s; dark green bars in a and b) periods of the photostimulation (colored shading, bootstrap 95% CI for each experimental group; light grey, bootstrap 95% CI for chance). **d**, Changes induced by laser stimulation in conditional transition probabilities between the three motor states for each group during the late (20-25 s) photostimulation period, evaluated using a Markov chain. Values indicate the average change in transition probability and solid and dashed colored lines highlight significant increases and decreases, respectively, during stimulation as compared to the baseline period.

**Fig. 5.**
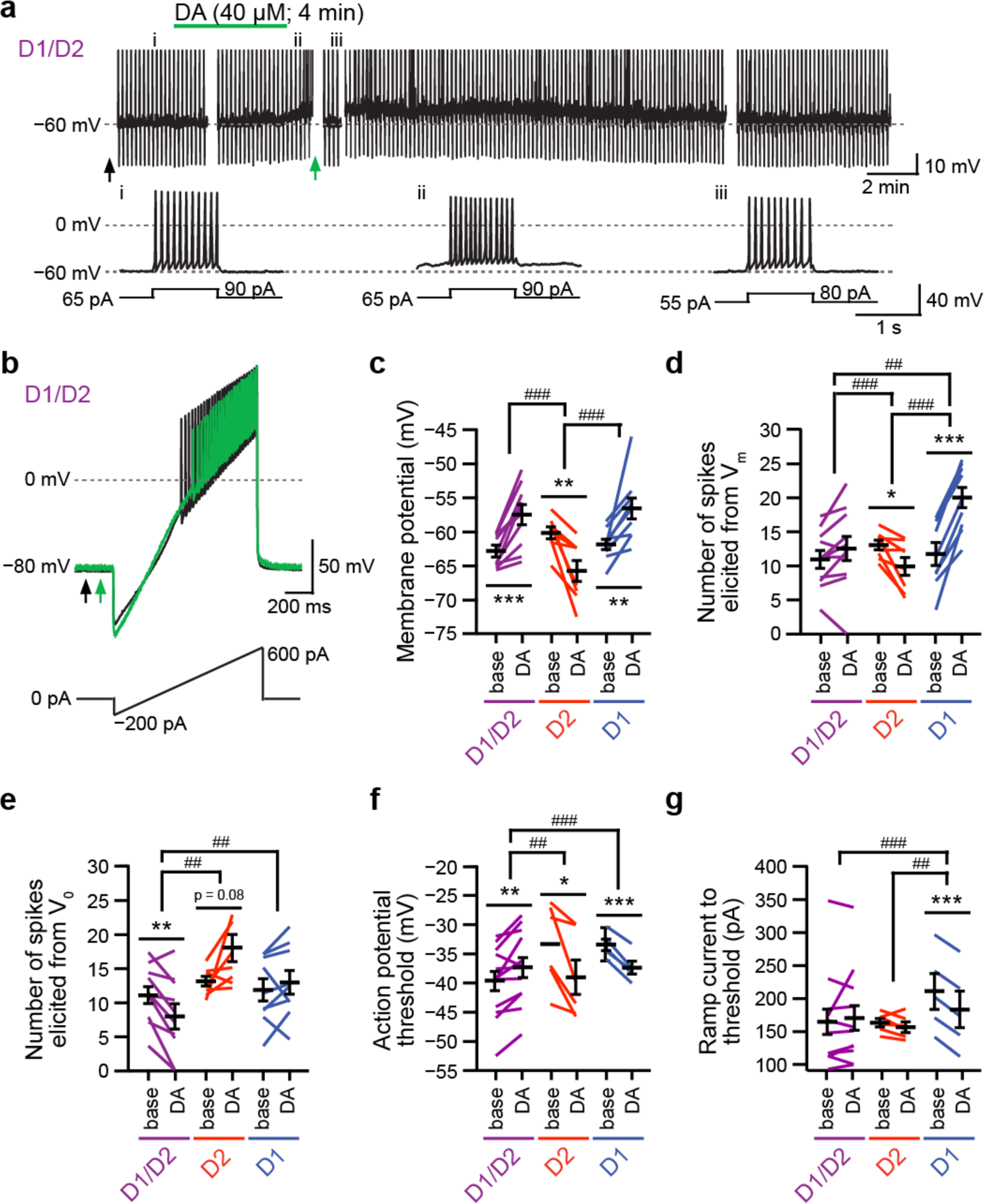
Co-expression of D1 and D2 cancels dopamine-induced changes in excitability. **a**, Effect of bath applied DA (40 µm, 4 min) on membrane potential and evoked firing activity in one D1/D2-SPN (top), and highlights on voltage traces recorded in response to 25 pA depolarizing current steps at times indicated by i-iii (bottom). **b**, Voltage response in one D1/D2-SPN to ramp current injection before (black) and after (green) DA application at times indicated by arrows in **a**. **c-e**, Quantification of DA-induced changes in membrane potential (**c**), number of elicited spiked by a depolarizing pulse from current membrane potential (**d**), and from initial membrane potential (**e**), in D1- (n = 9 cells from 4 mice), D2- (n = 8 [**c-d**], 7 [**e**] cells from 4 mice) and D1/D2-SPNs (n = 11 [c-d], 10 [e] cells from 4 mice). **f-g**, DA-induced effects on AP threshold (**f**) and minimal ramp current to elicit discharge (**g**) in D1- (n = 5 cells from 4 mice), D2- (n = 6 cells from 4 mice) and D1/D2-SPNs (n = 12 cells from 4 mice) (paired comparison between baseline and after DA application: *p < 0.05, **p < 0.01, ***p < 0.001; magnitude of DA effect between SPN subpopulations: ^#^p < 0.05, ^##^p < 0.01, ^###^p < 0.001).

To refine our examination of the locomotion state, we checked the velocity profile and accelerations around the onset of a locomotion state and during locomotion (**Fig. 3f-h, Extended Data Fig. 5b-c**). Stimulation of D1/D2-SPNs as well as D2-SPNs increases both velocity and acceleration when animals engage in locomotion. Conversely, D1-SPN stimulation resulted in a slowing of the locomotor state with lower speeds and acceleration. We hypothesized that these behaviors may respond to the aversive or rewarding effects of indirect or direct pathway activation, respectively^41, 42^. We therefore performed optogenetic “real-time place preference (RTPP)” experiments in which laser stimulation is controlled by the entry and exit of the animals from a specific zone in the open field (**Extended Data Fig. 6**). We found that the stimulated zone was preferred during D1-SPN (exclusive and mixed) activation and avoided during D1/D2-SPN and D2-SPN (exclusive and mixed) stimulation (**Extended Data Fig. 6a**). In addition, we examined the velocity around locomotion episode onsets occurring within and outside the stimulated zone (**Extended Data Fig. 6b**). Consistent with our previous observations, we observed a decrease in velocity with D1-SPN stimulation and an increase with D1/D2- and D2-SPN stimulation within the stimulated zone. These observations reinforce our claim that the effects we observe on speed around locomotion episode onsets result from appetitive and aversive effects of these stimulations (Supplementary text and Supplementary Fig. 2). Overall, these results reveal very effective control of D1/D2-SPNs on motor states with a dual role in locomotion, depending on their coactivation with either D1-SPNs or D2-SPNs, revising at the same time simplistic bimodal outputs of prokinetic and antikinetic roles of SPNs exclusive to D1Rs and D2Rs.

**Fig. 6.**
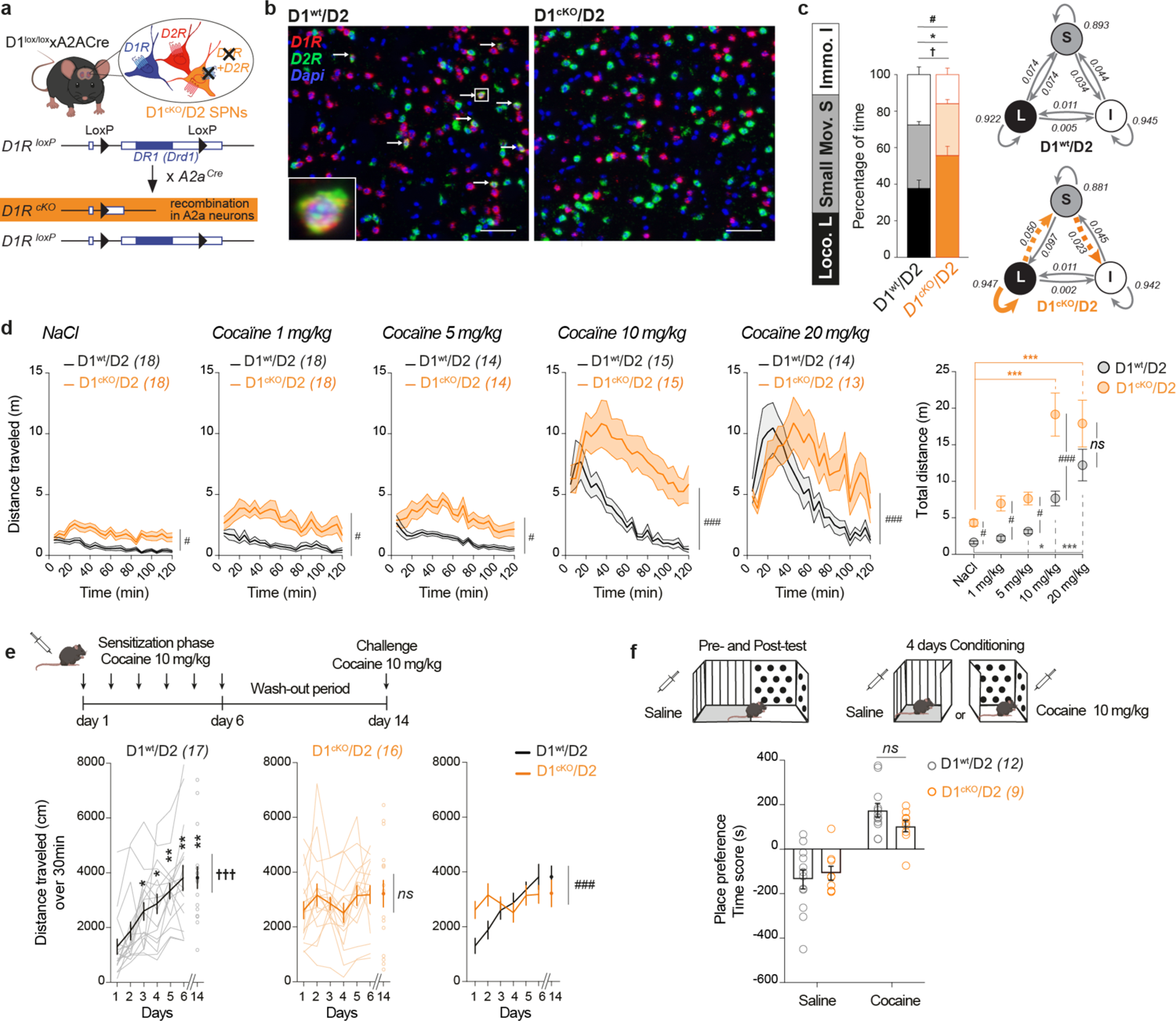
D1/D2-SPNs are necessary for proper *in vivo* integration of dopamine signals. **a**, Diagram illustrating generation of D1^cKO^/D2 mice. **b**, Photomicrographs from RNAscope in D1^wt^/D2 and D1^cKO^/D2 mice illustrating the suppression of DRD1 and DRD2 mRNAs co-expression. Arrows show double positive cells in a representative D1^wt^/D2 mouse with higher magnification on one of these cells in the square. Scale: 50 μm. **c**, Comparison of spontaneous motor behavior in an open field between D1^wt^/D2 mice (n = 12) and D1^cKO^/D2 mice (n = 15) evaluated through the time spent in each motor state (left; t-tests D1^wt^/D2 vs. D1^cKO^/D2: for locomotion, ^✝^p < 0.05; for small movements, **p < 0.01; for immobility, ^#^p < 0.05) and the conditional transition probabilities between each motor state (top right, D1^wt^/D2 mice; bottom right, D1^cKO^/D2; for the plot corresponding to D1^cKO^/D2 mice, orange solid and dashed lines respectively indicate significantly higher or lower probabilities than in D1^wt^/D2 mice). **d**, Temporal distribution of the pro-locomotor effect of acute cocaine injection (left panels; Sidak’s tests D1^wt^/D2 vs. D1^cKO^/D2 for entire duration: ^#^p < 0.05, ^##^p < 0.01, ^###^p < 0.001) and comparison of the dose-response relationships of the locomotor effect of cocaine in D1^wt^/D2 and D1^cKO^/D2 mice (right panel; Sidak’s test D1^wt^/D2 vs D1^cKO^/D2 for each dose: ^#^p < 0.05, ^##^p < 0.01, ^###^p < 0.001, ns not significant; Tukey’s tests vs. NaCl: * p < 0.05, ***p < 0.001). **e**, Effect of repeated

### Dynamic integration of D1/D2-SPNs reshapes the functional identity of D1- or D2-SPNs

We examined the kinetics of motor state change around each optogenetic stimulation event: before, during three periods throughout the light stimulation (early, 0-5 s; intermediate, 10-15 s; and late, 20-25 s), and after stimulation (10 trials per mouse; **Fig. 4a-c**) stimulation.

First, D1/D2-SPN stimulation produced a consistent increase in immobility at the expense of small movements, without any notable alteration in locomotion (Fig 4a-c, Extended Data Fig. 7). These analyses then reveal major differences between exclusive D1-SPNs and mixed D1+D1/D2-SPNs in response to stimulation (**Fig. 4a-c**). Immediately at the onset of stimulation (**Fig. 4a-c**), activation of D1+D1/D2-SPNs produces a decrease in locomotion, which quickly reverses into a large increase in locomotion. This effect on locomotion is mirrored by the temporal evolution of small movements (**Fig. 4a-c, Extended Data Fig. 7**). Conversely, activation of D1-SPNs only produces an increase in locomotion after 20 s, and small movements are increased throughout the whole duration of stimulation (**Fig. 4a-c, Extended Data Fig. 7**). For both D1-SPN and D1+D1/D2-SPN stimulations, the inhibition of immobility follows a similar temporal evolution, immediate and maintained throughout stimulation. These locomotion-promoting effects during simultaneous stimulation of D1- and D1/D2-SPNs are consistent with the stabilization of the locomotor state underpinned by a lengthening of locomotor episodes, not observed in stimulation of exclusively D1-SPNs (**Extended Data Fig. 5a**). In contrast, “exclusive” D1-SPN stimulation results mainly in the induction of small movements, which is prevented by the costimulation of D1/D2-SPNs (**Extended Data Fig. 5c**). To extend this characterization of motor state changes, we applied a Markov chain model to assess how the dynamic organization of motor states is affected by each SPN stimulation (**Fig. 4d, Extended Data Fig. 8a**). Conditional transition probabilities highlight that activation of D1-SPNs alone facilitates small movements by inhibiting exit probabilities from this state. Conversely, this convergence to small movements is less strong when D1-SPNs are coactivated with D1/D2-SPNs because transitions from locomotion to small movements are inhibited and maintenance of locomotion is strengthened (**Fig. 4d, Extended Data Fig. 8a**). Altogether, these findings indicate that activation of D1/D2-SPNs is required to trigger the prokinetic effects associated with activation of the classical direct pathway.

These comparative analyses also identified differences in the exclusive and mixed D2-SPN populations, illustrated by a divergence of effects on state changes to the locomotor state depending on whether D1/D2-SPNs are coactivated or not (**Fig. 4a-c**). Despite a quick and persistent facilitation of immobility (Fig. 4a-c, Extended Data Fig. 8a-b) in both cases, late during the stimulation of exclusive D2-SPNs, the probability of being in the locomotor state is increased (**Fig. 4a-c**), whereas such an effect is prevented by costimulation of D1/D2-SPNs (**Fig. 4a-c**). During most of the stimulation period, D2+D1/D2-SPN stimulation inhibits locomotion. This effect fades away late during the stimulation (Fig. 4a-c). Consistently, computing conditional transition probabilities under D2-SPN and D1/D2+D2-SPN activations indicates an increased probability of remaining in the locomotor state only under stimulation of D2-SPNs (**Fig. 4d, Extended Data Fig. 8a**). These results indicate that coactivation of D1/D2-SPNs alongside D2-SPNs blocks a late prokinetic effect induced by exclusive stimulation of D2-SPNs and thus further facilitates the antikinetic role of indirect pathway activation.

### Dopamine exerts a unique effect on D1/D2-SPN activity

Given the fundamental role of DA in locomotion in the striatum, we explored its actions and functions on these hybrid SPNs. We first evaluated *ex vivo* the effects of DA bath application on SPN activity (**Fig. 5, Extended Data Fig. 9**). We confirmed that at depolarized membrane potentials, DA depolarizes D1-SPNs, whereas it hyperpolarizes D2-SPNs (**Fig. 5c, Extended Data Fig. 9b**). Consistently, it increases D1-SPN excitability and decreases D2-SPN excitability, as indicated by substantial changes in the firing response to depolarizing current pulses, membrane resistance, and the current-frequency relationship (**Fig. 5d-e, Extended Data Fig. 9c-e**). In addition, concerning AP properties, we confirmed a decrease in the AP threshold in both populations in response to DA (**Fig. 5f**), and the ramp current to the AP threshold was significantly decreased in D1-SPNs, whereas it was unchanged in D2-SPNs (**Fig. 5g**).

For D1/D2-SPNs, we found DA-induced depolarization in a D1R activation-dependent manner (**Fig. 5a-c, Extended Data Fig. 9c, h**) but no change in excitability or firing (**Fig. 5d-e, Extended Data Fig. 9a-e**). This lack of effect might be related to the peculiar D1/D2-SPN AP properties previously described (**Fig. 2**). Indeed, in sharp contrast to the two other SPN populations, we observed that DA increases the AP threshold in D1/D2-SPNs (**Fig. 5b, f**), which is abolished in the presence of either D1R or D2R antagonists (Extended Data Fig. 9k-m). In addition, the ramp current to AP threshold is unchanged (**Fig. 5g**). These observations highlight the singularity of D1/D2-SPNs that distinctly integrate DA signals as a result of their peculiar coexpression and potential coactivation of D1Rs and D2Rs.

### Loss of D1/D2 coexpression impairs the *in vivo* DA response

To substantiate the importance of DA signaling integration in D1/D2-SPNs *in vivo*, we generated a model in which the D1R gene was deleted selectively in D2-SPNs (D1^cKO^/D2 mice). As a result, the D1^cKO^/D2 mice have a true engineered two striatal output pathway system, with direct D1-SPN output and one single indirect striatal output with all neurons only expressing D2Rs (**Fig. 6a-b**). In this model, the D1/D2 output is therefore suppressed, with two clear consequences.

First, D1^cKO^/D2 mice displayed significantly higher spontaneous locomotor activity (**Fig. 6c** and Extended Data Fig. 10a) characterized by more time spent in the locomotion state and less immobility than control littermates (D1^wt^/D2). This difference in locomotion between D1^cKO^/D2 and D1^wt^/D2 mice is conserved when animals receive acute doses of psychostimulants: 1, 5 or 10 mg/kg cocaine (**Fig. 6d**) and 1 mg/kg amphetamine (Extended Data Fig. 10b). Conversely, no difference was observed for the highest doses of cocaine or amphetamine tested (**Fig. 6d, Extended Data Fig. 10b**). Moreover, the temporal integration of the cocaine response differs between D1^cKO^/D2 mice and controls.

Second, D1^cKO^/D2 mice are characterized by a significantly different pattern of motor adaptive response to the cocaine sensitization paradigm (**Fig. 6e**): daily cocaine administration does not trigger an increase in locomotion in D1^cKO^/D2 mice, as observed in control animals (**Fig. 6e, Extended Data Fig. 10c**). The locomotor response showed a similar time course as the response to acute administration (Extended Data Fig. 10c). Even though this difference is striking, it could be either that the D1^cKO^/D2 mice are not sensitizing or that alternatively they are already in a “presensitized” state. In addition, the analysis could be confounded by the atypical delayed and long-lasting effects following cocaine administration, which is even more marked during the first days of conditioning (Extended Data Fig. 10c).

These findings demonstrate that D1/D2-SPNs are essential players in an integrated motor response to DA neurotransmission in collaboration with D1-SPNs and D2-SPNs. Notably, the rewarding effects of cocaine were not affected, as cocaine place preference was maintained in D1^cKO^/D2 mice (**Fig. 6f**), indicating that the phenotype we observed seems to engage motor circuits rather than reward-related pathways. Altogether, these observations highlight an essential role for D1/D2-SPNs in the integration of DA signals and in the functioning of the striatum, which cannot operate properly with only two efferent pathways.

## Discussion

Our findings demonstrate that the striatum computes and relays information through three efferent pathways. We highlighted a third population of SPNs in the striatum that is distinguished by coexpression of D1Rs and D2Rs and that projects specifically to the GPe. These neurons may partially account for the described genetic heterogeneity of SPNs^22–27^. Accordingly, further functional subdivisions between SPN subtypes may arise by taking into consideration the existence of genetic distinction between SPNs located in the matrix or in striosomes or located in the medial or lateral subdivisions of the dorsal striatum^22–23^. These additional subtypes would likely display unique properties such as their input‒output connectivity or their integration of various neurotransmitter signals that should be linked with specific functions during defined behaviors. Thanks to the comprehensive characterization of genetic markers expressed by SPNs^22–27^, addressing these issues could be achieved using a combination of multiple genetic markers, as we did in this study, with two-factor intersectional tools. This approach could also be enhanced using three-factor intersectional methods^43^.

Our optogenetic studies revealed that D1/D2-SPNs provide key control over the global functioning of the dorsal striatum to control appropriate transitions between motor states. We observed that D1/D2-SPNs, when coactivated with D1- or D2-SPNs, act as conductors of striatal locomotion and shape the functional identity of D1- and D2-SPNs as described thus far, i.e., pro- and antikinetic, respectively (Supplementary text). Our study also revealed new effects of D1- and D2-SPN excitation on motor states, which differ from bidirectional control of locomotion, and are characterized by a locked state in the promotion of small movements instead of locomotion under D1-SPN activation or a strong permissiveness to locomotor initiations under D2-SPN activation. Optogenetics is a powerful tool to dissect neuronal circuits but also poses great challenges. Stimulation protocols eliciting a neuronal response should ideally be as close as possible to the physiological one, which is remarkably difficult, as these are not always characterized. *In vivo* recordings demonstrate that SPNs are relatively quiet (< 5 Hz) and change their firing rate in response to discrete stimuli or actions in less than a second or during a couple of seconds^44–46^. As a consequence, one may argue that prolonged stimulation can lead to artificial effects not observed in natural conditions (Supplementary text). However, in the context of dopamine loss or cocaine use, long-lasting excitation or inhibition of some SPNs can occur^21, 47^. We therefore believe that additional experiments assessing the activity pattern of each of the three SPN populations during different tasks will be essential to better understand the functions of D1/D2-SPNs and how the three subpopulations co-orchestrate behavioral control.

Moreover, disruption of this pathway, which comprises a small subset of SPNs, has major functional consequences for spontaneous behavior and integration of DA signals. The striking difference in psychostimulant effects in D1^cKO^/D2 mice clearly demonstrates that under normal conditions, DA activation of D1Rs in hybrid D1/D2-SPNs has opposite effects to D2R stimulation of indirect D2-SPNs in inhibiting locomotion. In the D1^cKO^/D2 mice, this motor inhibition could no longer be effective, but a more prominent locomotor behavior is observed, and the effects of cocaine are enhanced (or no longer buffered). This is a direct consequence of the unique electrophysiological properties of D1/D2-SPNs. Indeed, although D1/D2-SPNs displayed DA-induced depolarization following D1R activation, their firing rate remained unchanged. Therefore, upon glutamate activation by cortico-striatal and thalamo-striatal principal neurons, and in contrast to D2-SPNs that are inhibited by DA^48^, D1/D2-SPNs will promote the activation of the indirect pathway resulting in motor inhibition. Consequently, D1/D2-SPNs, by maintaining an adequate inhibitory tone on the GPe, provide a necessary balance between the direct and indirect pathways to prevent DA-mediated convergent inhibition of the basal ganglia output nuclei, leading to overactivation of the thalamo-cortical systems. We therefore propose that D1/D2-SPNs provide a key balance necessary for proper striatal functioning. This balance between D1/D2-, D1-, and D2-SPNs is likely formed during embryonic stages and matures during early postnatal development, following a stage where more SPNs coexpress both D1Rs and D2Rs^49–51^. Only a fraction of SPNs in the indirect pathway will conserve the expression of both D1Rs and D2Rs, whereas most express only D2Rs, and all SPNs in the direct pathway express only D1Rs. It remains to be determined how these less abundant D1/D2-SPNs interact with and integrate into the other two SPN populations in different areas of the striatum during more elaborate striatal-dependent behaviors^52^.

Improper establishment of this balance or its subsequent dysregulation could lie at the core of certain neurophysiological disorders involving a dysregulation of striatal function. For example, the positive symptoms of schizophrenia are related to a hyperdopaminergic state in the mesolimbic system^53^ and in particular in the dorsal striatum^54^. Thus, abnormal coordination of the three striatal output pathways, and in particular the one formed by D1/D2-SPNs, could be an unsuspected susceptibility risk for psychiatric disorders, and could be involved in motor control in neurological disorders such as Parkinson’s disease.

## Acknowledgments

We thank Marc G Caron, Ami Citri, Philippe Faure, Christophe Kellendonk, Christian Lüsher, Marina Piciotto, Gilad Silberberg and Laurent Venance for critical reading of the manuscript. We thank Charu Ramakrishnan and Karl Deisseroth for providing additional Intrsect viruses. We thank Delphine Houtteman, Souad Laghmari, Perrine Hagué and Laetitia Cuvelier for technical assistance and mouse colonies in Brussels, Yeqing Geng for taking care of the mouse colonies in Montréal and Stéphanie De Gois for assistance in imaging techniques. PB is a Research Associate of FRS-FNRS (Belgium). CV is a postdoctoral researcher of FRS-FNRS (Belgium), and GF is a postdoctoral researcher of FRQS (Canada). PMO, ADG, AC and EPF are research fellows of FRS-FNRS-FRIA (Belgium). AKE is a Research Director of the FRS-FNRS and a WELBIO investigator. BG is a Canada Research Chair and a visiting professor at Université de Paris.

## Author contributions

Conceptualization: BG, PB and AKE; Design and investigation for optogenetic experiments: PB, with support from PMO; Methodology and investigation for motor analysis: PB, CV with support from AC; Methodology and investigation for electrophysiological experiments: CV; Investigation for 3D reconstruction: ADG; Investigation for behavioral experiments on D1^cKO^/D2: GF, QR with support from KX and EI, analysis: PB and CV; Investigation for the FISH experiments: SD with support from AC for automated analysis; Investigation for viral-based cell quantification and tracing: PB in collaboration with AC and RLL (automated analysis), with support from EPF (viral injections); Supervision for the animal colonies: EV; Resources for the D1flpO strain: ET; Formal analysis: PB and CV; Validation and supervision: PB, BG and AKE; Writing: PB, CV, AKE and BG.

## Data and materials availability

All data are available in the manuscript or the supplementary materials. The three transgenic mouse strains will be shared after completion of an MTA to prevent conflicts of interest on research subjects.

## Materials and Methods

### Generation of transgenic mice

*D1-FLPo (D1^FlpO^) mice.* D1^FlpO^ mouse strains were engineered at the Mouse Clinical Institute (Institut Clinique de la Souris, MCI/ICS, Illkirch, France). Briefly, the cassette containing the Flipase-O recombinase-T2A-GFP and a floxed NeoR/KanR selection cassette were inserted at the ATG of the D1 dopamine receptor gene by homologous recombination into the RP24-266B BAC (Fig. S2A). The modified BAC after deletion of the selection cassette was microinjected into 300 C57Bl/6N oocytes and several founders have been obtained. F1 offsprings were evaluated for the expression pattern of the Flp-O (Institut Clinique de la Souris, Illkirch, France).

*D1^loxP/loxP^ mice.* The floxed D1DAR mice were produced as follows. The KOMP ES cell line Drd1tm1a(KOMP)Wts, clone EPD0507_1_B12, derived on the JM8A3.N1 cell line, were obtained from the KOMP Repository at the University of California at Davis (UC Davis). The ES Cells were generated under NIH Grant U01HG004080 by the CHORI-Sanger-Davis (CSD) consortium. Mice were generated from the ES Cells for our research by the UC Davis Mouse Biology Program (MBP), www.mousebiology.org, using blastocyst microinjection, resulting chimeras were mated to C57BL/6NTac mice and heterozygous tm1a (Knockout First) animals were generated through germline transmission. These ES cells are now distributed through the Mutant Mouse Resource and Research Center (MMRRC) at UC Davis, an NIH-funded strain repository, RRID: MMRRC_054799-UCD. For more information on obtaining KOMP products please or go to www.mmrrc.org.

### Mice breeding

A2A-Cre (A2a^Cre^) driver line, which expresses the Cre recombinase selectively in D2-SPNs, excluding cholinergic interneurons^35^ were crossed with D1^FlpO^ mice (D1^FlpO^/A2a^Cre^) to perform interstional approaches to target the various SPNs subsets and also used for control conditions. Single transgenic A2a^Cre^ as well as D1^FlpO^ mice were used to assess the functions of classically characterized D2- and D1-“total” SPNs populations, respectively. All mice were of C57BL/6J genetic background.

The brain-specific D1R-depleted mice in A2a expressing neurons (D1^cKO^/D2) were obtained by crossing heterozygous D1^loxP/loxP^ mice with A2a^cre^ mice in order to obtain heterozygous offsprings, which crossed thereby produced wildtype (WT; 7/16), knockout (KO; 3/16) and heterozygote (HET; 6/16) mice at the expected ratio. After weaning and sexing, mice were housed in groups of 4-5 animals per cage.

For the collection of *in vivo* optogenetic data, *ex vivo* electrophysiology and FISH experiments, 6-10 weeks old male and female mice were used. Experiments related to D1^cKO^/D2 knockout also used both males and females. Two to four mice were housed together, in polycarbonate cages at constant temperature (21±1 °C) and humidity (40–50%), under 12-hour light/dark cycle. Mice were provided with food and water *ad libitum*.

For experiments taking place in place in Belgium, animals were treated according to protocols and guidelines approved by the Institutional Animal Care Committee and were approved by the local ethics committees (ULB Pôle Santé – Protocol 695N). For experiments taking place in France, experimental procedures were approved by the Regional Ethics Committee No. 3 of Ile-de-France region on Animal Experiments and followed the guidelines of the European Communities Council Directive (86/809/EEC) and the Ministère de l’Agriculture et de la Forêt, Service Vétérinaire de la Santé et de la Protection Animale (agreement A 94-028-21). For experiments taking place in Canada, animal housing, breeding, and care were operated in accordance with the Canadian Council on Animal Care guidelines (CCAC) and all methods were approved by the Animal Care Committee from the Douglas Institute Research Center (protocol number 5570). All methods were performed in accordance with the relevant guidelines and regulations.

### Double-probe fluorescent *in situ* hybridization (FISH)

#### Sample collection

Mice were euthanized by cervical dislocation. Brains were removed, rapidly frozen in cold isopentane (between –30 °C and –35 °C) and have been serially sectioned in 10 series on cryostat at 16 µm thickness. sections for each brain have been processed by in situ hybridization to allow systematic quantitative analysis throughout the whole dorsal and ventral striatum. Double-probe FISH were performed as previously reported^55^

#### DNA template for riboprobe synthesis

DNA templates with T3 and T7 promoter sequences on both sides of each cDNA target were produced by PCR using promoter-attached primers.

#### Probes

Double-probe FISH was performed using antisense riboprobes for the detection of the following mRNAs: D1, NM_010076.3 sequence 1756-2707 and D2, NM_010077.3 sequence 268-1187. Synthesis of digoxigenin and fluorescein-labeled RNA probes were made by a transcriptional reaction with incorporation of digoxigenin or fluorescein labeled nucleotides (Sigma-Aldrich; References 11277073910 and 11685619910). Information of primer sequences for riboprobe synthesis is listed in the table below. Specificity of probes was verified using NCBI blast.

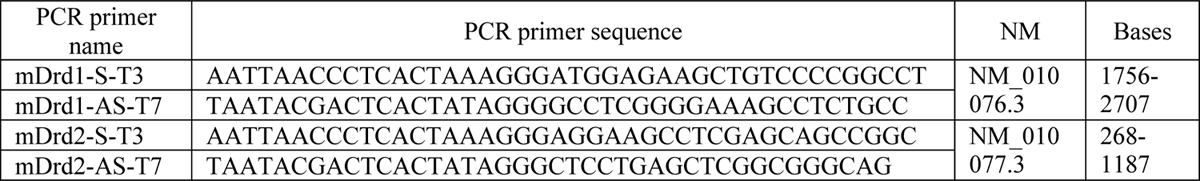

#### Procedure

Cryosections were air-dried, fixed in 4% paraformaldehyde and acetylated in 0.25% acetic anhydride/100 mM triethanolamine (pH 8) followed by washes in PBS. sections were hybridized for 18 h at 65 °C in 100 µl of formamide-buffer containing 1 µg/ml DIG-labeled riboprobe and 1 µg/ml fluorescein-labeled riboprobe. sections were washed at 65 °C with SSC buffers of decreasing strength and blocked with 20% FBS and 1% blocking solution. For revelation steps, DIG epitopes were detected with HRP anti-DIG fab fragments at 1:2500 (Sigma-Aldrich; Reference 11207733910) and revealed using Cy3-tyramide at 1:100. Fluorescein epitopes were detected with HRP anti-fluorescein fab fragments at 1:5000 (Sigma-Aldrich; Reference 11426346910) and revealed using Cy2-tyramide at 1:250. Cy2-tyramide and Cy3-tyramide were synthesized as previously described^56^. Nuclear staining was performed with DAPI. All slides were scanned at 20x resolution using the NanoZoomer 2.0-HT (Hamamatsu, Japan). Laser intensity and time of acquisition was set separately for each riboprobe.

#### Automated quantitative analysis

10 sections per brain in a total of 3 animals were selected for the quantification. The dorsal and ventral parts of the striatum were differentiated for the quantification. A CellProfiler pipeline^57^ was developed to analyze imaging data. Signal segmentation was performed through recognition of objects based on size, shape, intensity, and texture of the signal. All sections were treated with the same segmentation. Cell segmentation was performed on Dapi staining to delineate cell surface area. We used the IdentifyPrimaryObject method to identify Dapi positive cells with the following parameters: typical diameter of object (15–40 pixels), adaptive threshold strategy, two-classes Otsu thresholding method, and the distinction of clumped objects was based on shape and minimal distance between local maxima of 5 pixels. Next, we established a pixel median threshold value in each fluorescent canal: red (D1-mRNA) and green (D2-mRNA). D1- and D2-positive cells were considered with a median pixel intensity > 0.25 and > 0.10, respectively. We then performed an automated quantitative analysis in which neurons were separated between D1-mRNA only-(dapi + red), D2-mRNA only-(dapi + green) and D1/D2-co-expressing (dapi + red + green) cells in the dorsal and ventral striatum.

#### RNAScope fluorescent *in situ* hybridization

Frozen brain samples were serially cut using a cryostat (10 µm in thickness) on superfrost-charged slides and kept at −80 °C until further processing. *In situ* hybridization was performed using RNAScope probes and reagents from Advanced Cell Diagnostics and according to the manufacturer’s instructions. Briefly, sections were first fixed in chilled 10% neutral buffered formalin for 15 min at 4 °C, dehydrated in increasing gradients of ethanol baths (50%, 70%, 95% 100%, 100%) and left to air dry for 5 min. Endogenous peroxidase activity was quenched with hydrogen peroxide reagent for 10 min, followed by protease digestion for 10 min at room temperature. The following sets of probes were then hybridized for 2 h at 40 °C in a humidity-controlled oven (HybEZ II, ACDbio): MsDrD1 (cat. No. 461901), MsDrD2 (cat. No. 406501-C2) to quantify D1/D2 coexpression. Successive addition of amplifiers was performed using the proprietary AMP reagents, and the signal visualized through probe-specific horseradish-peroxidase-based detection by tyramide signal amplification with Opal dyes (Opal 520 and Opal 570; Perkin Elmer) diluted 1:300. Slides were then coverslipped with Fluoromount-G mounting medium with 4,6-diamidino-2-phenylindole (DAPI) for nuclear staining (Invitrogen Fisher, cat. No. 501128966) and kept at 4 °C until imaging.

#### Virus-mediated targeting of channelrhodopsin and eYFP expression

We used an intersectional optogenetic viral strategy to trace, record from and control each SPNs population, involving the stereotaxic injection of D1^FlpO^/A2a^Cre^ mice with INTRsT viruses^33^ transfecting cells with eYFP-tagged ChR2(H134R), or eYFP alone for *ex vivo* recordings and control mice, in a manner conditional on the presence of Cre (**C**) and/or FlpO (**F**) in three configurations: **C**off/**F**on, **C**on/**F**off and **C**on/**F**on, targeting respectively D1-, D2- and D1/D2- SPNs (AAV(DJ)-nEF-**C**on**F**on / **C**off**F**on / **C**on**F**off-hChR2(H134R)-eYFP, AAV(DJ)-hSyn-**C**on**F**on / **C**off**F**on / **C**on**F**off-eYFP; some aliquots were kindly provided by Charu Ramakrishnan and Karl Deisseroth at Stanford University, and most viral vectors were packaged and provided by UNC vector core, North Carolina). To compare ChR2-induced effects in response to photostimulations in these three exclusive SPNs populations with those classically described in mixed direct D1-expressing SPNs (combining D1- and D1/D2-SPNs) and mixed indirect D2-expressing SPNs (combining D2- and D1/D2-SPNs), we also expressed ChR2 in D1^FlpO^ and A2A^Cre^ mice using FRT-ChR2 and lox-ChR2 constructs, respectively (AAV(DJ)-EF1a-fDIO-hChR2(H134R)-eYFP, AAV(5)-EF1a-DIO-hChR2(H134R)-eYFP; UNC vector core).

At 8–10 weeks old, mice used for tracing, quantification of virus-transfected cell densities, assessment of transfection specificity (Extended Data Fig. 1f) and *in vivo* experiments, or at about 6 weeks old, mice used for *ex vivo* experiments were anesthetized with isoflurane (4% induction, 1% maintenance; 0.5 L/min O2). 2×1 μL of recombinant AAV listed above were bilaterally injected under stereotaxic control into the dorsal striatum (anteroposterior (AP), +0.8 mm; mediolateral (ML), 1.5 mm; dorsoventral (DV), –2.7 mm, relative to Bregma) through a 30 G cannula at a rate of 100 nL/min. The injection needle was lowered through a hole drilled lateral to the midline (AP, +0.8 mm; ML, 1.5 mm). Mice were allowed to recover for 10 days before being habituated to patchcord connections and experiments started at least 3 weeks after viral injections. To quantify cell densities transfected with eYFP virus, we automated the quantification of positive cells (cells pointed with a red dot: Extended Data Fig. 1g-h), nuclei, and ROI area by using in-house python scripts. To minimize false positives, we eliminated detected cells with lower central fluorescence compared to their periphery (cells pointed with a blue dot). For this, the cell central region was defined as a circle with a radius equal to one-fourth of the cell’s diameter and the average fluorescence in this region was compared to the average fluorescence in the rest of the cell.

### *Ex vivo* electrophysiology experiment recording and analysis

#### Experiment preparation

In preparation for electrophysiology, brain slices containing the dorsal striatum were obtained from mice injected as described above. Under deep anesthesia the animals were decapitated and brains were quickly removed. Coronal slices (250 μm thick) were cut with a vibrating microtome (VT1000S; Leica) in ice-cold (4 °C) slicing artificial cerebrospinal fluid (ACSF) containing the following (in mM): 140 choline chloride, 2.5 KCl, 1.25 NaH_2_PO_4_, 7 MgCl_2_, 0.5 CaCl_2_, 25 NaHCO_3_, 14 D-glucose, pH 7.3, and constantly oxygenated (95% O2/5% CO2). Slices were transferred to the incubation chamber filled with ACSF containing (in mM): 124 NaCl, 2.5 KCl, 1.2 NaH_2_PO_4_, 2 CaCl_2_, 1 MgCl_2_, 26 NaHCO_3_, 10 D-glucose. Slices were incubated for 1 h at 34 °C and then maintained at room temperature.

#### Whole cell recordings

Slices were placed on the stage of an Axio Examiner.A1 microscope (Carl Zeiss), equipped with an infrared CCD camera (ORCA-05G; Hamamatsu), and visualized by under infrared (IR) illumination. Slices were maintained immersed and continuously surperfused at 2–5 mL/min with oxygenated ACSF. Neurons in the dorsal striatum were selected based on eYFP expression using 470 nm LED light (OptoLED, Carin Research) visualized through FITC/Cy2 filter set (Carin Research). All electrophysiological experiments were performed with an EPC-10 patch-clamp amplifier (HEKA) attached to a computer running PatchMaster software (HEKA). Signals were low-pass filtered at 2.9 kHz and digitized at 20 kHz. Whole-cell patch-clamp recordings were performed with patch micropipettes (6–8 MΩ) pulled from borosilicate glass capillary tubes (1.5 mm outer diameter, 0.225 mm wall thickness; Hilgenberg) on a vertical puller (PIP5; HEKA) filled with 8 µl of internal solution containing the following (in mM): 105 K-gluconate, 30 KCl, 10 4-(2-hydroxyethyl)-1-piperazineethanesulfonic acid, 0.3 ethylene glycol-bis(β-aminoethyl ether)-N,N,N’,N’-tetraacetic acid, 10 Na_2_-phosphocreatine, 4 Mg_2_-ATP, 0.3 Na_2_-GTP, pH 7.4, 290–300 mOsm, and 3 mg/mL biocytin. For *ex vivo* optogenetic stimulation, pulses of blue light (470 nm) were delivered using LEDs (OptoLED, Cairn Research) controlled by PatchMaster (HEKA) through the EPC-10 patch-clamp amplifier (HEKA).

#### Recording analysis

The intrinsic membrane properties of neurons were assessed by applying current steps (1s) from –500 pA to firing saturation in 10 pA increments. To comprehensively describe the electrophysiological properties of recorded SPNs, 30 parameters reflecting the properties of SPNs^36–38, 58^ measured on traces corresponding to the voltage responses induced by hyperpolarizing and depolarizing current pulses. These parameters are following the Petilla nomenclature^59^. (1) Resting membrane potential (RMP) was measured immediately after electrical access to the intracellular compartment of the cell. (2) Input membrane resistance (R_m_), (3) membrane time constant (τ_m_), and (4) membrane capacitance (C_m_) were determined in response to hyperpolarizing current pulses resulting in voltage drops of 10–15 mV relative to rest. R_m_ was measured following Ohm’s law and τ_m_ was computed as the time constant of a single exponential fit of the voltage response from onset to maximum hyperpolarization. C_m_ was obtained following the formula C_m_ = τ_m_/R_m_. Some neurons have been described to undergo a partial repolarization following a hyperpolarization peak in response to negative currents. To quantify this, whole-cell conductance was measured when the sag conductance was inactive (G_hyp_) or active (G_sag_). G_sag_ was measured as the slope of the linear portion of a current–voltage (I–V) plot, in which V was determined at the end of 1 s hyperpolarizing current pulses (–500 to 0 pA) and G_hyp_ as the slope of the linear portion of the I–V plot where V was determined as the maximal negative potential during the 1 s hyperpolarizing pulses. (5) Sag index was calculated as a relative decrease in membrane following (G_sag_ – G_hyp_)/G_sag_. (6) Step intensity to first spike (I_th_) was evaluated as the minimal intensity of the 1 s current pulse that elicited an action potential. As SPNs display a delayed firing supported by I_A_ currents, (7) first spike latency (lat) was computed as the time needed to elicit a spike after the onset of a current pulse corresponding to the step intensity to first spike. (8) Action potential threshold (V_th_) was measured as the potential of first spike onset when the rate of membrane potential change exceeded 30 mV/ms. To describe the firing behavior of recorded SPNs around threshold, the instantaneous firing frequency measured in response to the minimal current injection eliciting more than 3 action potentials was and fitted to a linear curve with (9) adaptation (A_th_) corresponding to the slope of the linear fit and (10) minimal steady state frequency (F_min_) corresponding to its y-intercept (F(t) = A_th_ x t + F_min_). Then to evaluate the change in firing activity in relation with increased depolarizing current steps (11) the slope of the current-frequency relationship (I–f slope) was calculated in its initial linear portion. (12) The maximum step intensity before depolarization block (I_sat_) was defined as the maximal 1 s current pulse eliciting robust firing activity without inducing firing saturation. At high firing frequencies, neurons display spike frequency adaptation which is characterized by an early exponential decrease in spike frequency followed by a linear decrease. To quantify this adaptation, the instantaneous firing frequency during the train of spike induced by the maximal depolarizing current injected not inducing saturation was fitted to decaying single exponential and of a linear function using the following formula F(t) = A_sat_ x exp(–t/τ_sa_t) + m_sat_ x t + F_max_^60^ using the fitting parameters defined as (13) amplitude of early adaptation (A_sat_), (14) time constant of early adaptation (τ_sat_), (15) slope of late adaptation (m_sat_), and (16) maximal steady state frequency (F_max_). To evaluate firing adaptation at an intermediate level of activation (17) spike frequency adaptation (SFA) was measured as the ratio between the instantaneous firing frequency between the first two spikes over the instantaneous firing frequency over the last three spikes on traces corresponding to the depolarizing current injection eliciting a sustained action potential train of about 20 spikes. Spike waveforms have been described to vary between neuronal cell types. To quantify these variations, 6 parameters were measured on the first and sond spikes of traces corresponding to the depolarizing current injection eliciting a sustained action potential train of about 20 spikes. (18,19) The amplitudes of the first (A1) and sond action potential (A2) were defined as the difference of membrane potential between their onsets and peaks. (20,21) Spike durations of the first (D1) and sond spike (D2) were defined the spike with between spike onset and spike offset. (22,23) Spike half-widths of the first (W1) and second spike (W2) were defined as the spike width at half amplitude. (24,25) After hyperpolarization potential (AHP) maximums (AHP1 and AHP2) were calculated using the membrane potential at the onset of the action potential as a reference. Comparatively, (26,27) AHP latencies (tAHP2 and tAHP2) were calculated as the time span between the AHP maximum and the onset of the action potential. During a spike train, spikes can display variations in their amplitude and duration. In order to quantify this phenomenon, (28) amplitude reduction (A.Red), (29) duration increase (D.Inc), and (30) half-width duration increase (W.Inc) were computed as (A1 – A2)/A1, (D2 – D1)/D1, and (W2 – W1)/W1, respectively.

#### Pharmacology

The following drugs were used: dopamine HCl (40 µM, Sigma-Aldrich); sulpiride (10 µM, Sigma-Aldrich); and SCH23390 (10 µM, Sigma-Aldrich). Stock solutions of these compounds were stored as frozen aliquots at –20 °C and dissolved in ACSF to their working concentrations immediately before application. Pharmacological experiments were carried out in current-clamp mode. Neurons were depolarized to about –60 mV by injecting holding current. Changes in membrane voltage were measured at this holding current. To elicit firing activity and quantify membrane resistance, neurons were stimulated using a 1 s depolarizing pulse set to elicit 10–15 action potentials and after 1 s held at holding current neurons were stimulated with a –50 pA hyperpolarizing pulse from holding current lasting 500 ms. This stimulation protocol was repeated every 10 s throughout the recording. Moreover, a current ramp (1 s from –200 pA to 600 pA) was applied during baseline and at the peak of pharmacological effect to extract action potential threshold and current to first action potential as well as interpolate current-frequency curves. Recordings were discarded if access resistance changed by more than 20% throughout the experiment. Results are not corrected for liquid junction potential.

### Three-dimensional reconstruction of biocytin-filled SPNs

Brain slices containing SPNs filled with biocytin, were incubated overnight at 4 °C in paraformaldehyde solution (4% PFA in 0.01 mM PBS). Slices were then washed in PBS and permeabilized in PBS with 0.1% TritonX-100 (Sigma-Aldrich). Slices were incubated with streptavidin-coupled AlexaFluor488 (1:200, Jackson ImmunoResearch) for 2 h at room temperature under constant agitation. After washing in PBS, slices were mounted with Fluorsave medium (Fisher Scientific) between slide and coverslip (VWR).

Serial optical sections in the X, Y and Z planes (Z-stacks) were acquired on a laser-scanning confocal system (LSM 510; Zeiss) mounted on an Axiovert 200M inverted microscope (Zeiss) and equipped with a LD c-Apochromat 40×/1.2W objective (Zeiss). The excitation beam of an argon laser (488 nm) was used and the fluorescence was measured between 500 and 550 nm for the detection of the Alexa 488 fluorophore. Once acquired, ZEN software (Zeiss) was used to stitch the images. Manual reconstruction of the dendritic arborization of the neurons was done with the help of the computer filament tracing function in the 3D image analysis software Imaris (Bitplane, Ireland). Total dendritic arborization reconstruction allowed the study of dendrite branching and length, and sholl analysis. The soma volume was automatically measured by creating a surface that delineated the soma region.

### Optic fiber implantation for *in vivo* experiments

At 8–10 weeks old, within the same surgery following virus injection, mice subjected to *in vivo* experiments were implanted with homemade bilateral optic fiber implants. A cleaved piece of optic fiber ∼12 mm in length (400 µm diameter, NA 0.39; Thorlabs) was inserted and glued into a metal ferrule (1.25 mm diameter; Thorlabs). Optic fiber implants that were inserted were verified before implantation to display at least During the surgery, before viral injection, the skull was completely cleared of all connective tissue and thoroughly dried using alcohol. To insert the optic fiber implant, a hole was drilled in the skull at the following coordinates: AP, +0.9 mm; ML, 2.17 mm, relative to Bregma. After gently cutting through the dura, the implant was lowered through the hole at an angle of 10° with the optic fiber end location targeted 200 µm above the injection site in the dorsal striatum. Following optic fiber placement, dental cement (Metabond) was applied to permanently fix the implant to the skull.

### Rotational behavior

Animals were placed in circular chambers (17 cm diameter) to examine rotational behavior under unilateral stimulations. A camera (Basler ace acA1600-60gc) was placed above the chambers to record videos at 30 frames/s, and rotations were scored manually. Rotations were defined as each 360° rotation that contained no turn of more than 90° in the opposite direction. We analyzed them over three periods ‘Pre’ period, which corresponded to 60 s preceding laser illumination; ‘ON’ period, which corresponded to constant 60 s unilateral laser illumination (473 nm, Laserglow); and ‘OFF’ period of 60 s at the end of laser illumination. Both right and left hemisphere laser illuminations were performed in a randomized manner in two distinct trials separated by a 1-2 h interval period during which mice returned in their home cage.

### Open field measures

Open field is a white square enclosure of 40 x 40 x 40 cm where animals can move freely and are video-monitored from above (Basler ace acA1600-60gc; sampling rate 30 frames/s). Detection and calculation of the animal’s center of mass coordinates were performed using Ethovision xt 14.0 software (Noldus). Animal’s speed was calculated from changes in center of mass coordinates between two successive frames and smoothed (moving average, 0.5 s width) to reduce noise and tracking artifacts. Animal’s behavior in the open field was then classified into three motor states: locomotion when velocity reaches 5 cm/s and remains above 2 cm/s for at least 0.5 s; immobility when velocity is below 0.5 cm/s for at least 0.5 s; and small movement when velocity lays between 0.5 cm/s and 2 cm/s. Any unassigned bout (lasting shorter than 0.5 s) is affected for its first half to the previous state and for its second half to the next state. Averaged temporal evolutions of mice velocity and acceleration were examined around locomotion onset and measures of averaged velocity and top velocity (9^th^ decile) were analyzed only during locomotion episodes.

Animals were recorded for a 10 min reference baseline period before any laser illumination (‘Pre’ period). Mice exhibiting a biased low baseline activity measured as distance traveled below 1700 cm or above 4500 cm over the 10 min baseline were excluded from analysis. Then the laser was illuminated in a series of ten trials. Each trial lasted 30 s with constant illumination (‘ON’ period), followed by a 90 s period during which the laser was turned off (‘OFF’ period). The total time spent in each motor state, the frequency and the mean episode duration of each state were calculated over the 10 min baseline (‘Pre’), and over the average of the 10 trials for the ‘ON’ and ‘OFF’ periods. Other stimulation patterns were also tested: continuous 5Hz for 30 s (mid frequency - tonic activation; 10 trials with 90 s intervals) ii) 20Hz for 500 ms (phasic activation, 20 trials with 30 s intervals) and iii) continuous 20Hz for 30 s (high frequency - tonic activation, 4 trials with 90 s intervals).

To better characterize how each SPN population activation affects motor states dynamics we computed transition probabilities between states using a Markov chain model from time series of motor states resampled at 5 Hz. Briefly, for each time period (‘Pre’, ‘ON’, ‘OFF’) defined above and for each mouse we calculated the transition probability from state X to Y P(Y|X) as the number of time bins in state X immediately followed by state Y over the total number of time bins in state X.

To catch supplementary information on the dynamics of motor states during laser illumination, we also computed probabilities to observe motor states for all stimulation trials. The 95% CI for motor state probability were calculated using bootstrap from time series of motor states resampled at 10 Hz. For each experimental group, we repeatedly resampled data aligned to laser illumination (from 30 s before laser onset to 30 s after laser offset) by randomly generating the same number of total trials (random sampling with replacement). For each of the 10,000 random iterations, we calculated the mean probability to observe each motor state across trials. The lower and upper CIs were obtained for each time point from the distribution of resampled mean values. The 95% CI for changes in probabilities were obtained the same way by calculating the difference in mean probabilities during early (from 0 s to 5 s after laser onset), intermediate (from 10 s to 15 s after laser onset), and late (from 20 s to 25 s after laser onset) periods of laser illumination minus mean probabilities before laser illumination (over 10 s before laser onset). Chance levels for changes in probabilities were calculated the same way by selecting a random time point for trial onset. 95% CIs for actual trials lying outside the 95% CI for chance are considered significantly different. For open field experiments performed on D1^cKO^/D2 mice and their wildtype littermates, mice were placed in the open field and video-tracked for 60 min. All these analyses were performed using Matlab (Mathworks).

### Histological confirmation of optic fiber placement and construct expression

Following completion of all behavioral optogenetic experiments, mice were deeply anesthetized with one injection of avertin (2,2,2-tribromoethanol 1.25%, 2-methyl-2-butanol 0.78%; 20 μl/g, intraperitoneal, Sigma-Aldrich). Mice were then perfused transcardially with 1X PBS, pH 7.4, followed by 4% paraformaldehyde in PBS (PFA). The brains were then extracted and postfixed overnight in PFA at 4 °C and subsequently cryoprotected in 30% sucrose dissolved in PBS for an additional 24 h at 4 °C. Each brain was then frozen and sectioned at 40 μm using a cryostat. To confirm construct expression and optic fiber placement, 1:3 sections over the entire striatum were first washed in PBS 1X-Triton (0.3 %) (PBST), incubated in blocking solution (10% donkey serum (DS) in PBST) for 1 h at room temperature and subsequently incubated for 1 h with GFP boosters (ATTO488, Chromotek) diluted 1:400 in 1% DS. sections were then counterstained for 10 min with the chromosomal dye Hoechst 33258 (prepared in 1X PBS at 1:10,000) and then mounted on glass slides and permanently coverslipped with Fluoromount. Fluorescent images were collected using a fluorescent microscope (V16, Zeiss). Only mice with histologically confirmed bilateral correct placement of optic fiber as well as proper construct expression in the dorsal striatum were used in the present study.

### Psychostimulant dose-response and behavioral sensitization

Locomotor response to acute psychostimulant treatment was assessed by intraperitoneal administration of a single dose of amphetamine HCl (0, 1, 3 mg/kg in 0.9% saline) or cocaine HCl (0, 1, 5, 10, 20 mg/kg in 0.9% saline) to D1^cKO^/D2 mice and their wild-type littermates. The test environment consists in transparent activity boxes (VersaMax automated activity monitor, AccuScan Instruments Inc., Columbus, OH) measuring 42 cm x 42 cm x 30 cm. Mice were first habituated to the environment by being placed in the activity boxes for 20 min daily for 3 consecutive days. Acute injections were made on day 4: mice were individually placed in activity boxes for 15 min, and after injection were immediately placed back in the boxes for an additional 120 min period. Locomotor activity was recorded in 5 min intervals.

In order to induce behavioral sensitization, D1^cKO^/D2 mice and their wild-type littermates received daily administration of cocaine (10 mg/kg) for 6 consecutive days. On day 14, a challenge dose of cocaine (10 mg/kg) was administered to both groups. Injections were made 15 min after placing the mice in the monitor boxes and locomotion recording continued for 120 min after cocaine administration.

### Conditioned Place Preference

The Pavlovian conditioning protocol, using distinctive floor textures as conditioned stimuli, was adapted from Cunningham et al.^61^. The apparatus consisted of a Plexiglas box divided into two compartments (5 cm wide, 15 cm long, 15 cm high), with interchangeable floors made of either stainless steel or plastic. Each apparatus was equipped with photocell beams allowing automatic recording of the animal’s position and horizontal activity. The experiment involved three phases during which each animal was weighed and injected immediately before being placed in the apparatus. (i) During the habituation session, each animal received saline and had free access to both compartments with distinctive floors for 30 min. (ii) The conditioning phase (eight sessions) lasted four consecutive days, with two 30 min trials a day. Mice were restricted to a single compartment, with the less preferred floor paired four times with cocaine (10 mg/kg, 10 mL/kg, intraperitoneal) and the other floor texture paired four times with saline. (iii) Place preference was assessed 24 h after the last conditioning trial. All animals received a saline injection before being exposed to both compartments with distinctive floors for 30 min. The cocaine-conditioned preference was determined by comparing the time spent in the cocaine associated box post versus pre-chronic injection.

### Statistics Analysis

Statistical analysis was performed using GraphPad Prism 7.0e and Matlab. All data are presented as mean ± standard error of the mean (SEM) except explicitly stated otherwise. Statistical significance was set at 0.05 for all procedures. Normality and homoscedasticity assumptions were verified prior to the use of any parametric tests (Shapiro-Wilk normality test and equality of variances *F-test*). In case of violation of any of these assumptions, standard transforms were applied to meet the above criteria. Statistics were performed with repeated measures two-ways or three-ways ANOVAs to test the significance between experimental groups (genotype) and stimulation conditions (in case of optogenetics), or groups (floxed and wild-type littermates), drugs and time. Statistic tests details are available in the statistical table in the supplemental information. Cell count and *in vivo* analyses were performed by researchers blinded to genotype and experimental condition.

---

**Extended Data Figure 1.**
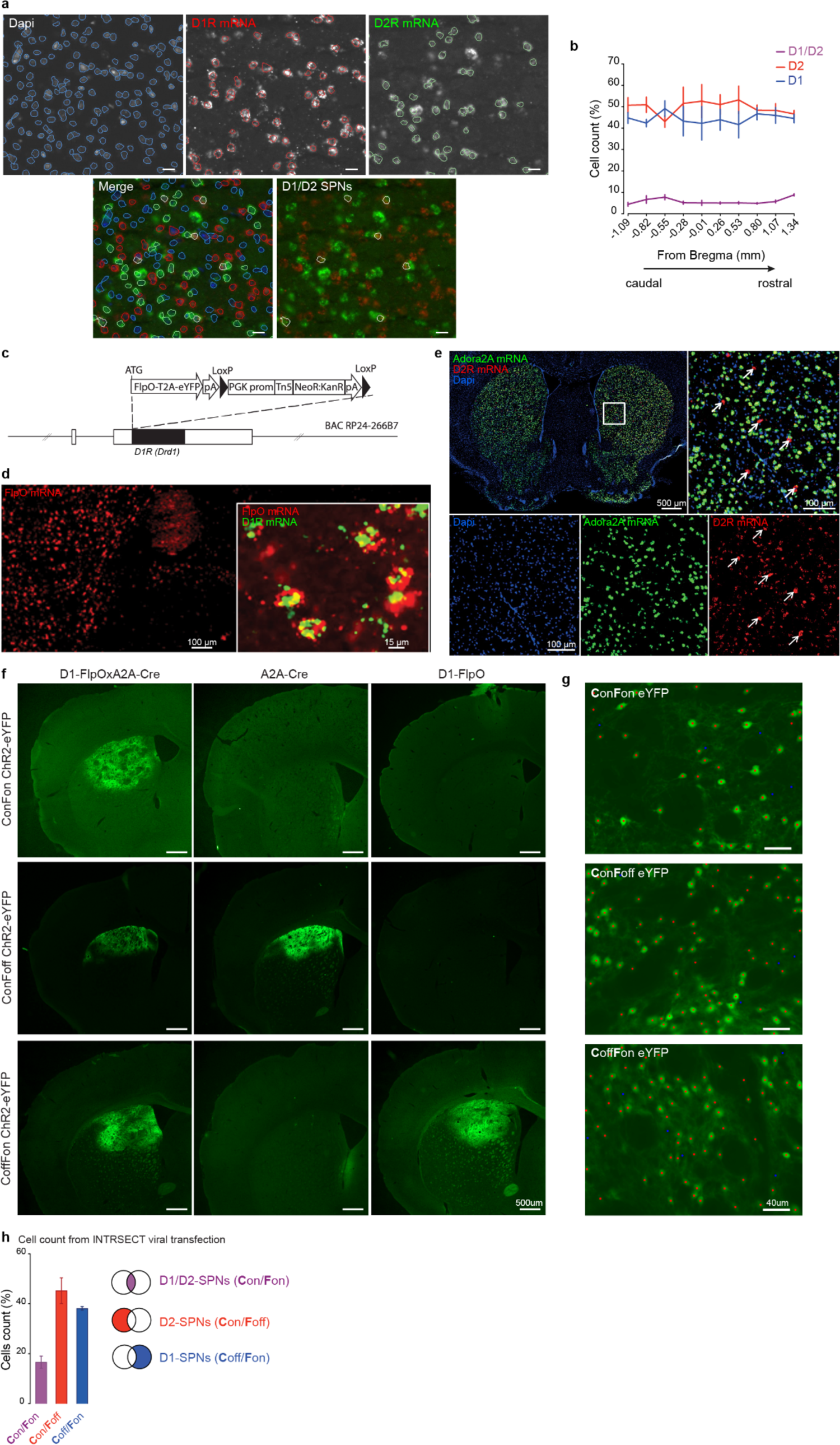
Characterization of D1/D2-SPNs with RNA FISH and their targeting with INTRSECT virus in a novel D1^FlpO^/A2a^Cre^ transgenic line. **a**, Representative coronal section in the dorsal striatum illustrating fluorescent in situ hybridization of D1R and D2R mRNAs and Dapi staining. Using a CellProfiler pipeline, segmentation was first performed on dapi staining to outline all cells (blue contours). Then D1- and D2-posivive cells are identified using an intensity threshold in red and green channels, respectively (red and green contours). Scale: 20 µm. **b**, Quantification of D1-, D2-, and D1/D2-positive cells throughout the rostro-caudal extent of the dorsal striatum (n = 3 mice). **c**, Diagram illustrating the generation of D1^FlpO^ mice. **d**, Photomicrograph of a coronal brain section at the level of the dorsal striatum after in situ hybridization of FlpO (red) and D1R (green) mRNAs illustrating expression of FlpO-recombinase in D1-expressing SPNs. Scales: left: 100 µm; right zoom: 15 µm. **e**, Coronal brain section after in situ hybridization of D2R (green) and Adora2A (red) mRNAs counterstained with Dapi (blue) illustrating specific expression of AdoraA2 in D2-SPNs^34^. Some cells expressing D2R and not AdoraA2 (arrows) correspond to cholinergic interneurons. Scales: top left whole section: 500 µm; zoom below: 100 µm. **f**, INTRSECT ChR2 virus specificity. Representative coronal sections showing **C**on/**F**on-(first line), **C**on/**F**off (second line), **C**off/**F**on-ChR2eYFP (third line) expression in double D1-FlpxA2A-Cre mice (left column), in single A2A-Cre line (middle column) and in single D1-Flp line (right column). We controlled the absence of leaky expression by verifying the absence of recombination of the Con/Fon construct in all single lines, as well as the absence of recombination of the Con/Foff construct in D1Flp or the Coff/Fon construct in A2A-Cre. Scale: 500 µm. **g-h**, Quantification of the density of INTRSECT virus transfected cells using automated detection. **g**, Representative sections in the dorsal striatum illustrating **C**on/**F**on-, **C**on/**F**off, **C**off/**F**on-eYFP virus expression, from top to bottom respectively. Red dots: positive cells. Blue dots: false positive cells with lower fluorescence. Scale: 40 µm. **h**, Density of cells transfected with **C**on/**F**on-eYFP (purple), **C**on/**F**off-eYFP (red) and **C**off/**F**on-eYFP (blue) virus. We found 228.3 ± 26.9 cells/mm^2^ for D1/D2-SPNs, 675.1 ± 58.2 cells/mm^2^ for D2-SPNs, and 561.2 ± 37.1 cells/mm^2^ for D1-SPNs which correspond to proportions of 16.6 ± 2.4% D1/D2-SPNs, 45.2 ± 5.2% D2-SPNs, and 38.2 ± 0.7% D1-SPNs relative to total SPNs (Dapi normalized, n = 3 mice for **C**on/**F**on-eYFP and **C**on/**F**off-eYFP virus, n = 2 mice for **C**off/**F**on-eYFP virus).

**Extended Data Figure 2.**
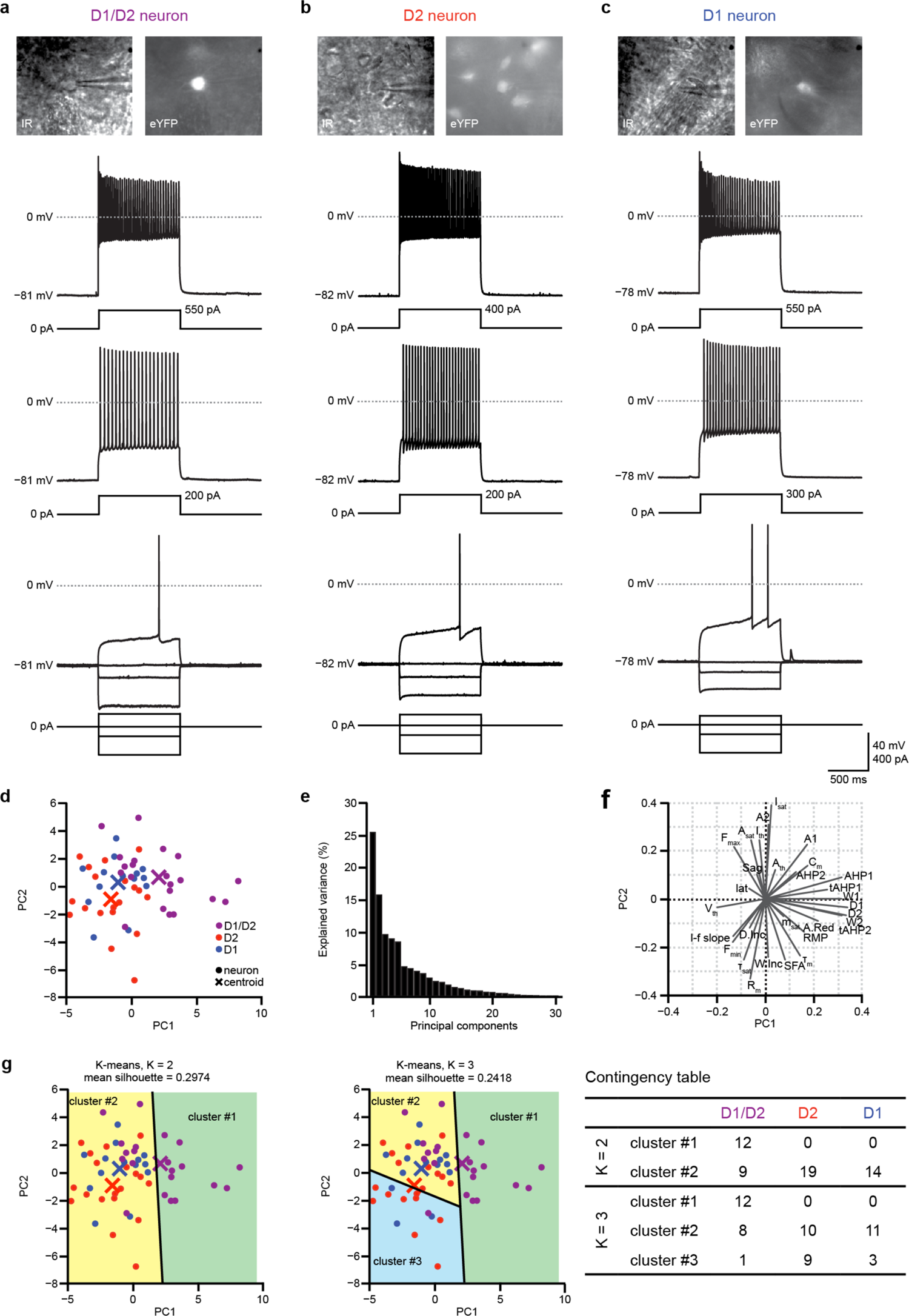
Electrophysiological characterization of the three SPNs subpopulations. **a-c**, Representative example of voltage response to current steps in one D1/D2-SPN (**a**), one D1-SPN (**b**) and one D2-SPN (**c**), with their corresponding photomicrograph under infrared illumination (IR, top left) and under fluorescent illumination (eYFP, top right). **d**, Two-dimensional projection of the set of neurons recorded following PCA into the first- and second-order components. Individual cells are represented in violet (D1/D2), blue (D1) and red (D2) circles; crosses represent centroids of D1/D2-SPNs (violet), D1-SPNs (blue) and D2-SPNs (red). **e**, Percent of variance explained as a function of the number of principal components. **f**, Two-dimensional projection of orthonormal principal component coefficients for each parameter. (*Abbreviations: RMP: resting membrane potential; R_m_: input membrane resistance; τ_m_: membrane time constant; C_m_: membrane capacitance; Sag: Sag index; I_th_: step intensity to first spike; lat: first spike latency; V_th_: action potential; A_th_: adaptation around threshold; F_min_: minimal steady state frequency around threshold; I–f slope: early slope of the current-frequency relationship; I_sat_: maximum intensity before depolarization block; A_sat_: amplitude of early adaptation at high firing; τ_sat_: time constant of early adaptation at high firing; m_sat_: slope of late adaptation at high firing; F_max_: maximal steady state frequency at high firing; SFA: spike frequency adaptation at high firing; A1: amplitude of the first action potential; A2: amplitude of the second action potential; D1: duration of the first action potential; D2: duration of the second action potential; W1: spike half-widths of the first action potential; W2: spike half-widths of the second action potential; AHP1: AHP maximum of the first action potential; AHP2: AHP potential maximum of the second action potential; tAHP1: AHP latency after of the first action potential; tAHP2: AHP latency of the second action potential; A.Red: amplitude reduction; D.Inc: duration increase; W.Inc: half-width duration increase*). **g**, Clusters obtained after k-means partitioning (left, k = 2; middle, k = 3) based on all the electrophysiological parameters recorded, and contingency table (right) comparing clusters assignments (lines) and cell identity defined by INTRSECT eYFP labelling.

**Extended Data Figure 3.**
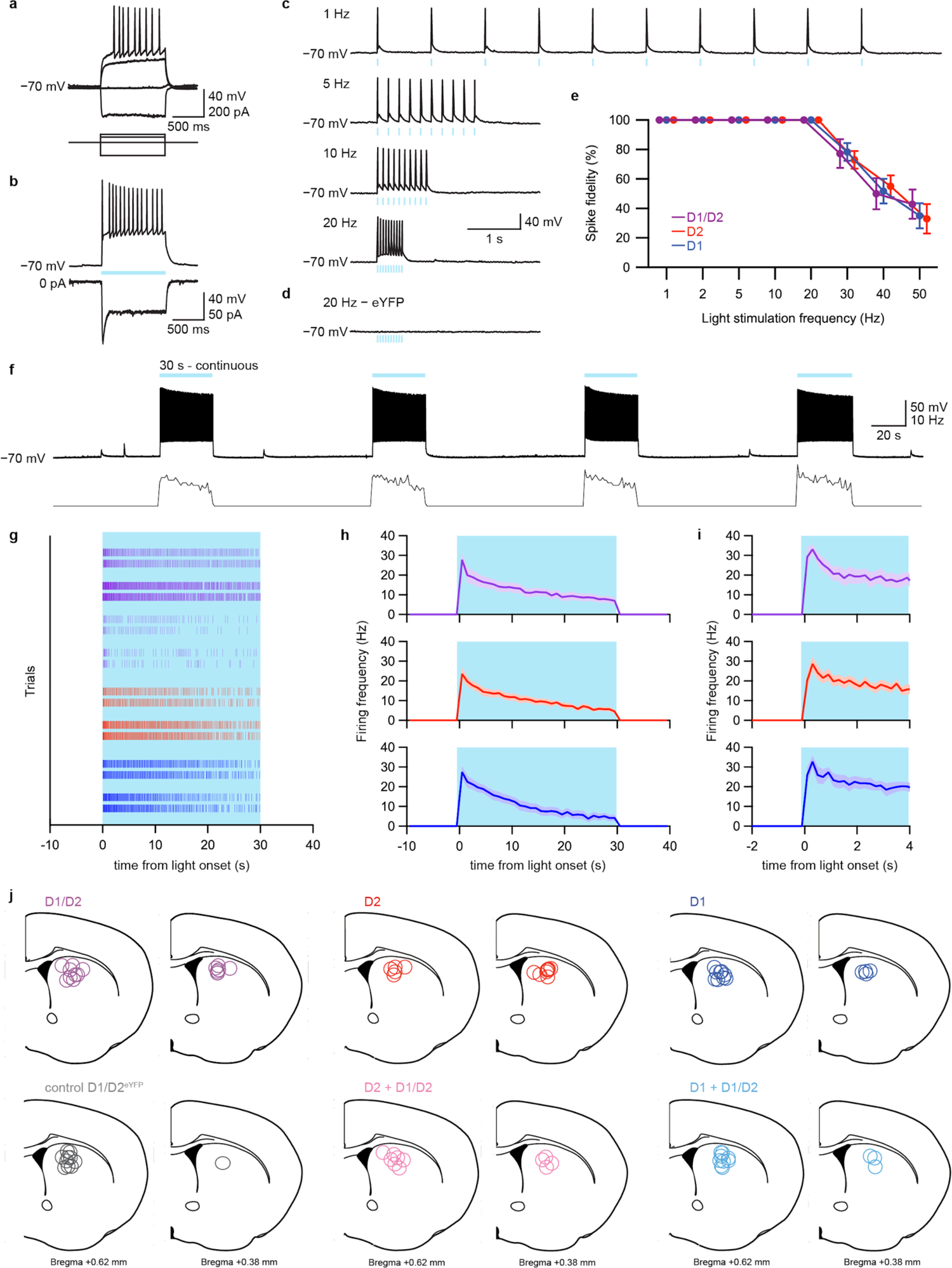
Ex vivo and histological controls for optogenetic experiments. **a**, Voltage response of an eYFP-expressing D1/D2-SPN to hyperpolarizing and depolarizing current injection displaying the typical features of SPNs. **b**, ChR2-expressing SPN (same as in A) displays robust depolarization and firing activity INTin response to a 1 s blue light illumination when recorded in current-clamp mode (top), and an inward current when recorded in voltage-clamp mode (bottom). **c**, Representative example of the same D1/D2-SPN depicted in **a** illustrating that pulses of light (5 ms) faithfully elicited spikes for frequencies up to 20 Hz. **d**, In a D1/D2-SPN only expressing eYFP, light stimulation is inefficient. **e**, Spike fidelity in response to light pulses at different frequencies (1-50 Hz) (D1/D2-SPNs, n = 9 cells from 6 mice; D1-SPNs, n = 6 cells from 4 mice; D2-SPNs, n = 7 cells from 4 mice). **f**, Representative example in one ChR2-expressing D1/D2-SPN of the spiking activity elicited by 30 s of continuous blue light illumnation. **g**, Raster plot of firing activity elicited by continuous light illumination in 2 consecutive trials in 4 representative D1/D2-SPNs (violet), 2 representative D2-SPNs (red), and 2 representative D1-SPNs (blue). **h**, Average firing activity calculated over 1 s bins elicited by 30 s of blue light illumination (D1/D2-SPNs, n = 12 cells from 5 mice; D2-SPNs, n = 9 cells from 3 mice; D1-SPNs, n = 10 cells from 3 mice). **i**, Same as **h** illustrated during the first 4 s of light stimulation calculated over 0.2 s bins. **j**, Schematic representation of the position of the tip of the optic fiber after histological control for all experimental groups (D1/D2, n = 21; control D1/D2eYFP, n = 11; D2, n = 15; D2 + D1/D2, n = 11; D1, n = 16; D1 + D1/D2, n = 10). Control mice correspond to D1^FlpO^/A2a^Cre^ mice injected with INTRSECT viruses transfecting cells with **C**on/**F**on-eYFP construct (D1/D2^eYFP^).

**Extended Data Figure 4.**
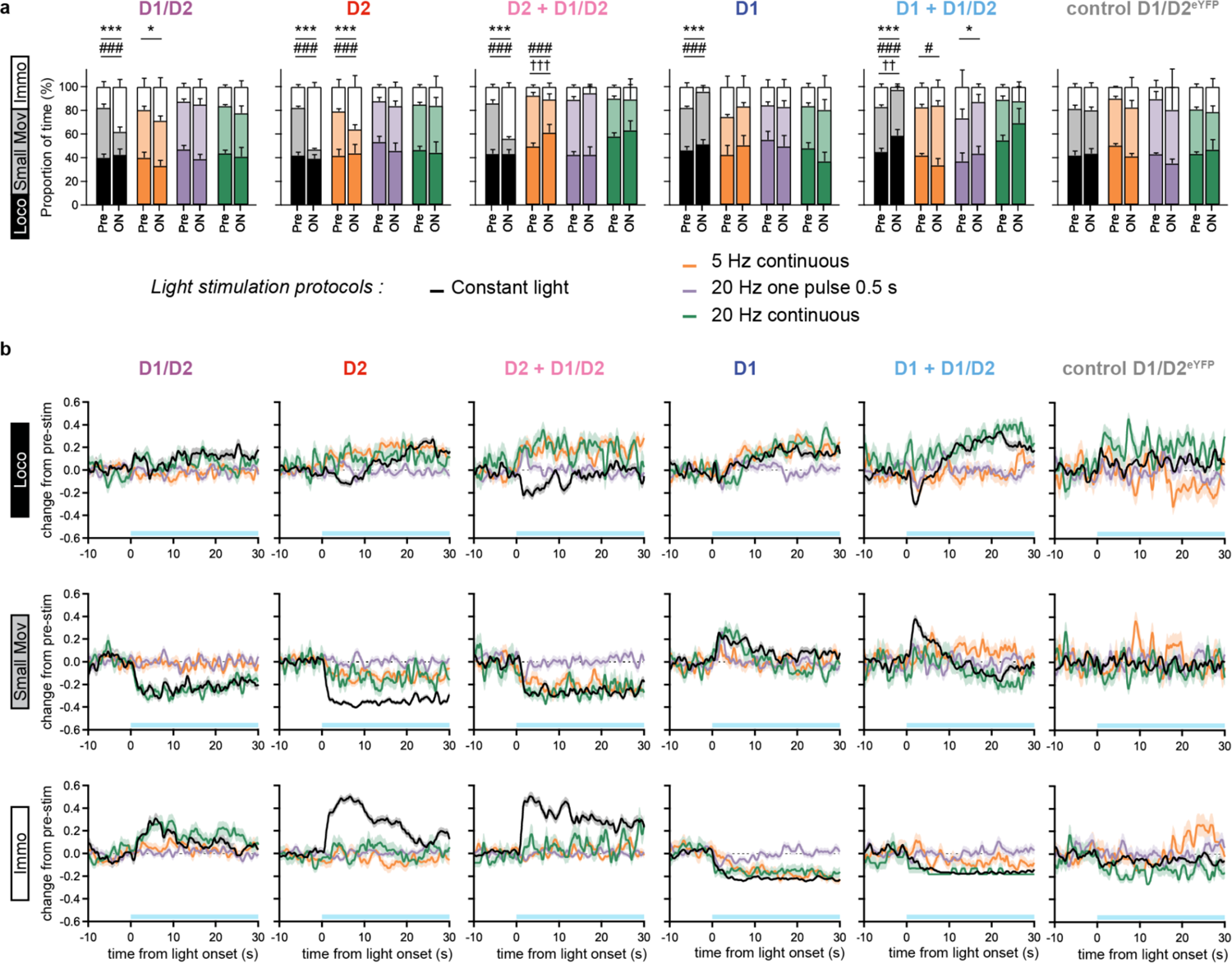
Comparison between optogenetic stimulation protocols to elicit behavioral changes. **a**, Effect of four different optogenetic stimulation protocols (constant light ON during 30 s, black; 5 Hz stimulation during 30 s, orange; one pulse at 20 Hz lasting 0.5 s, violet; 20 Hz stimulation during 30 s, green; D1/D2, n = 21, 8, 8, 13, respectively; D2, n = 15, 8, 8, 8, respectively; D2 + D1/D2, n = 11, 6, 6, 5, respectively; D1, n = 16, 6, 6, 11, respectively; D1 + D1/D2, n = 10, 5, 5, 5, respectively; D1/D2^eYFP^, n = 11, 5, 5, 6, respectively) on the proportion of time spent in locomotion, small movements, and immobility (Tukey’s test pre vs. light ON: for locomotion, ^††^p < 0.01, ^†††^p < 0.001; for small movements, ^#^p < 0.05, ^###^p < 0.001; for immobility, *p < 0.05, ***p < 0.001). **b**, Temporal evolution of changes in locomotion (top), small movements (middle), and immobility (bottom) occurrences induced by the four different optogenetic stimulation protocols constant light ON during 30 s, black; 5 Hz stimulation during 30 s, orange; one pulse at 20 Hz lasting 0.5 s, violet; 20 Hz stimulation during 30 s, green) (shading, bootstrap 95% CI).

**Extended Data Figure 5.**
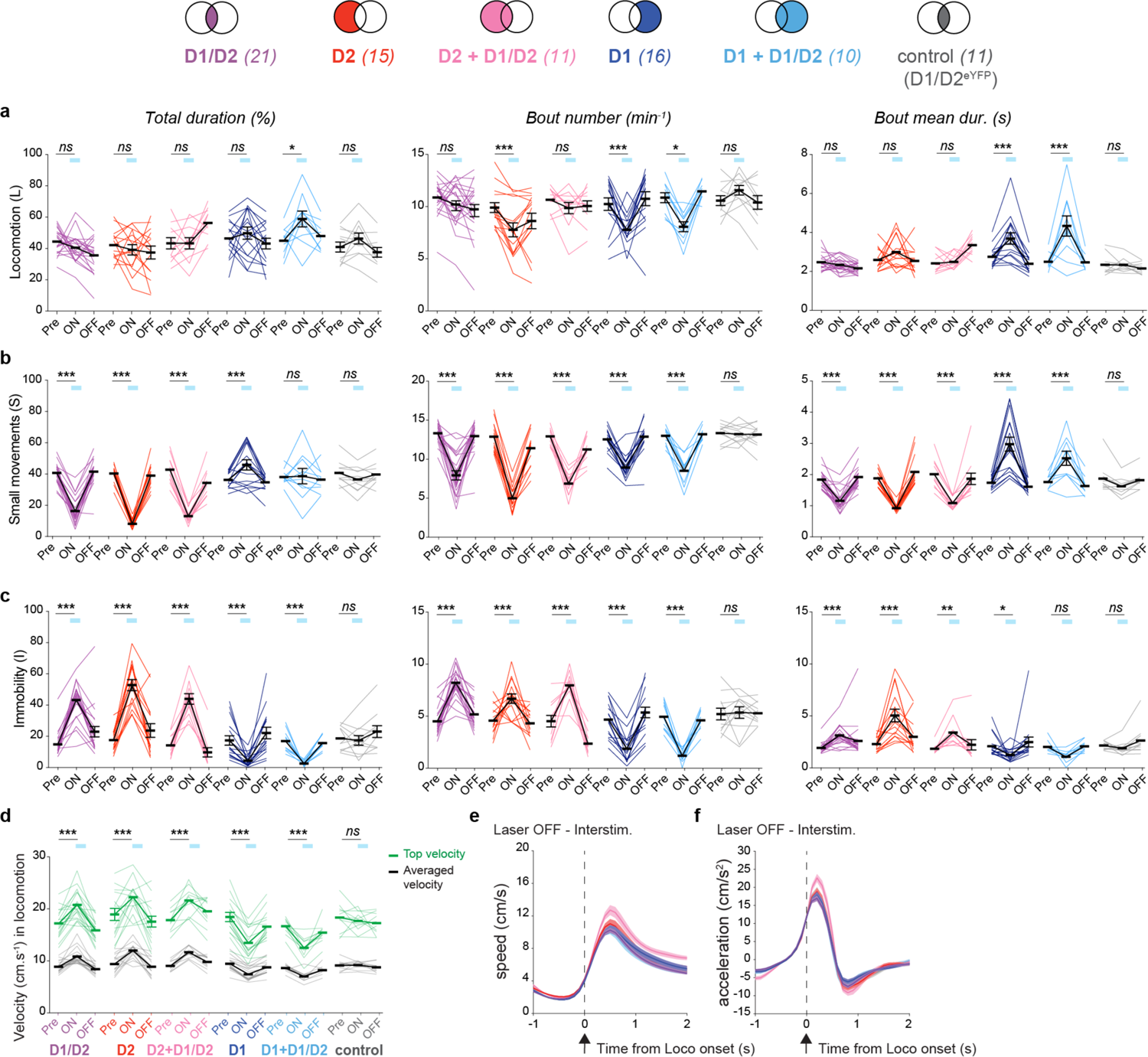
Architecture of motor states and locomotion velocity during optogenetic stimulation. **a-c**, Effect of optogenetic stimulation motor architecture during locomotion (**a**), small movement (**b**), and immobility (**c**) quantified by the proportion of time spent in each state (left panels), bouts frequency (middle panels), and the average episode duration (right panels) in D1/D2, control D1/D2^eYFP^, D1, D1 + D1/D2, D2, and D2 + D1/D2 mice (pre vs. light ON: *p < 0.05, **p < 0.01, ***p < 0.001). **d**, Effect of laser illumination on average velocity (black) and top velocity (9^th^ decile, green) during locomotion (pre vs. light ON: ***p < 0.001). **e-f**, Averaged temporal evolution of mice velocity (**e**) and acceleration (**f**) around locomotion onset during laser “OFF” periods between stimulation trials. For all graphs: D1/D2, n = 21; D2, n = 15; D2 + D1/D2, n = 11; D1, n = 16; D1 + D1/D2, n = 10 and control D1/D2eYFP, n = 11.

**Extended Data Figure 6.**
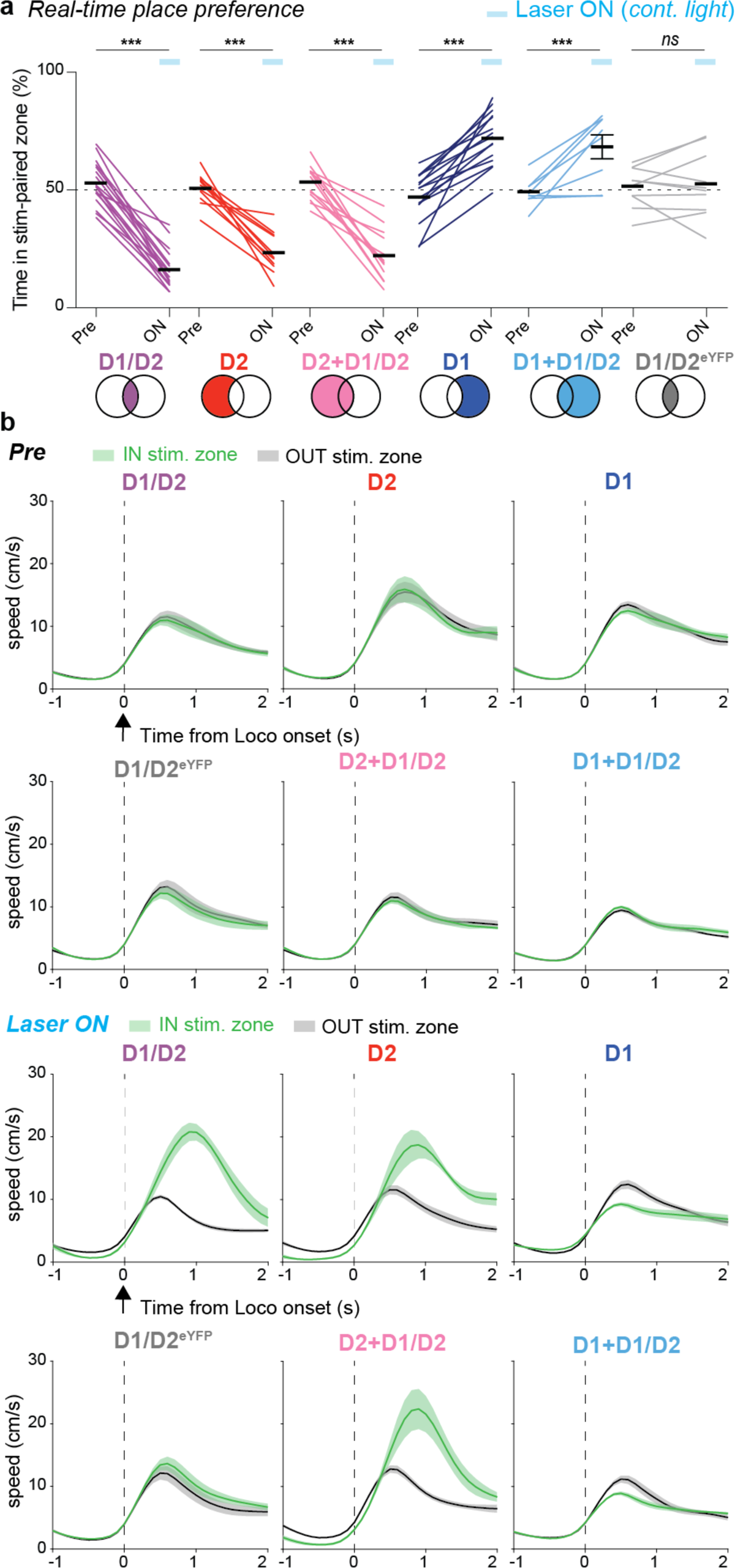
Effect of optogenetic stimulation on real-time place preference. **a**, Proportion of time spent in the laser-paired area before (Pre) and during (ON) the optical activation (continuous blue light) of D1/D2-SPNs (n = 21), D2-SPNs (n = 14), D2+D1/D2-SPNs (n = 14), D1-SPNs (n = 18), D1+D1/D2-SPNs (n = 9) or in control (D1/D2^eYFP^; n = 11) mice in the laser-paired zone (Pre vs. LaserON: ns p > 0.05, ***p < 0.001). **b**, Average temporal evolution of mice velocity aligned to locomotion episode onsets when the animal is exploring the laser-paired zone (green curves) or the unpaired zone (black curves) during the baseline period (Pre, top panels) and during the optogenetic stimulation in the laser paired zone (Laser ON, bottom panels). Note during the Laser ON condition, the large increase in locomotion velocity elicited by the stimulation of D1/D2, D2, and D2+D1/D2-SPNs suggesting the aversive effect of the stimulation, and the decrease in velocity elicited by the stimulation of D1, and D1+D1/D2-SPNs suggesting the appetitive effect of the stimulation.

**Extended Data Figure 7.**
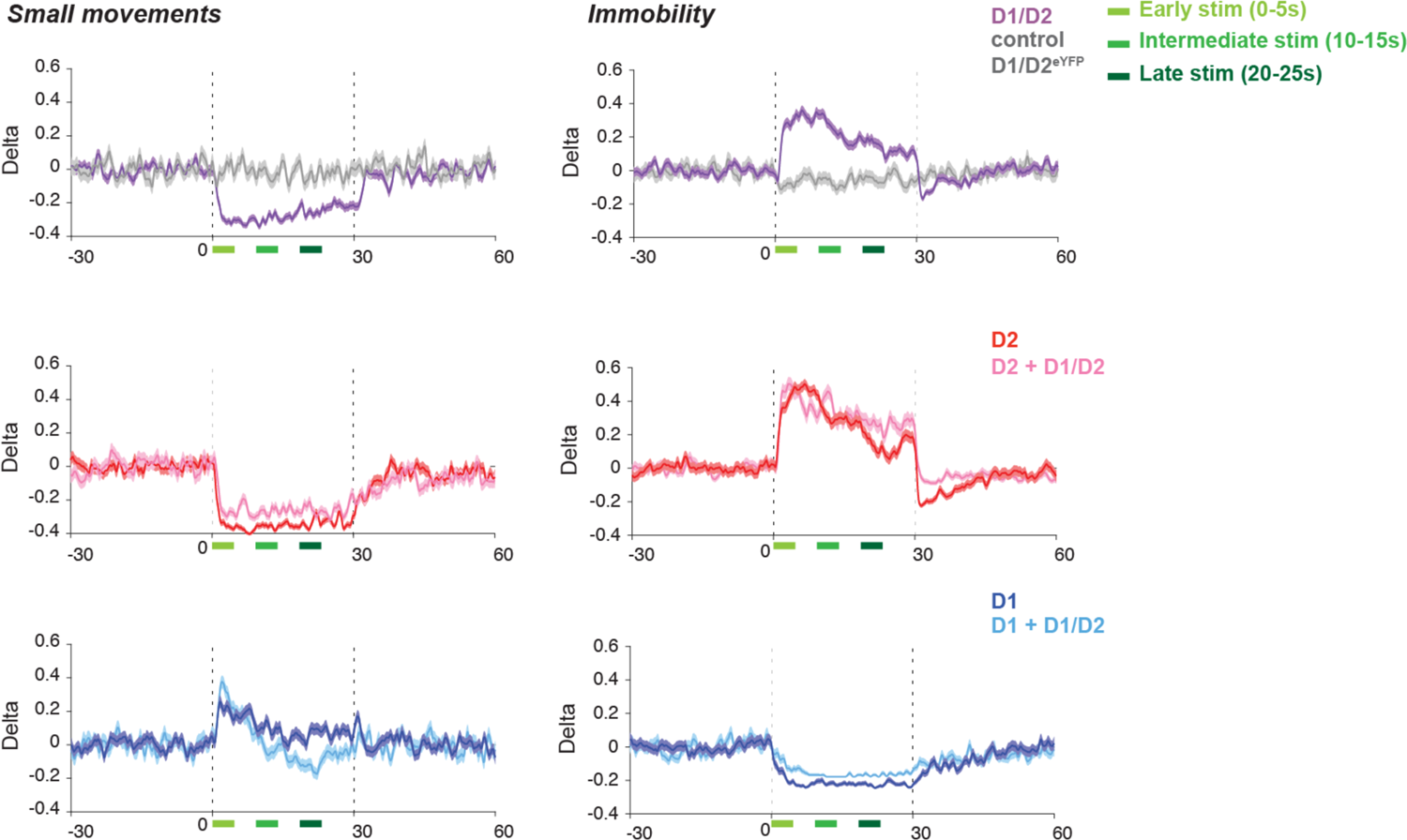
Dynamic changes in small movements and immobility occurrence during optogenetic stimulation. Temporal evolution of changes in small movements (left panels), and immobility (right panels) occurrences induced by optogenetic stimulation (constant light ON during 30 s; D1/D2, n = 21; D1/D2^eYFP^, n = 11; D2, n = 15; D2 + D1/D2, n = 11; D1, n = 16; D1 + D1/D2) (shading, bootstrap 95% CI).

**Extended Data Figure 8.**
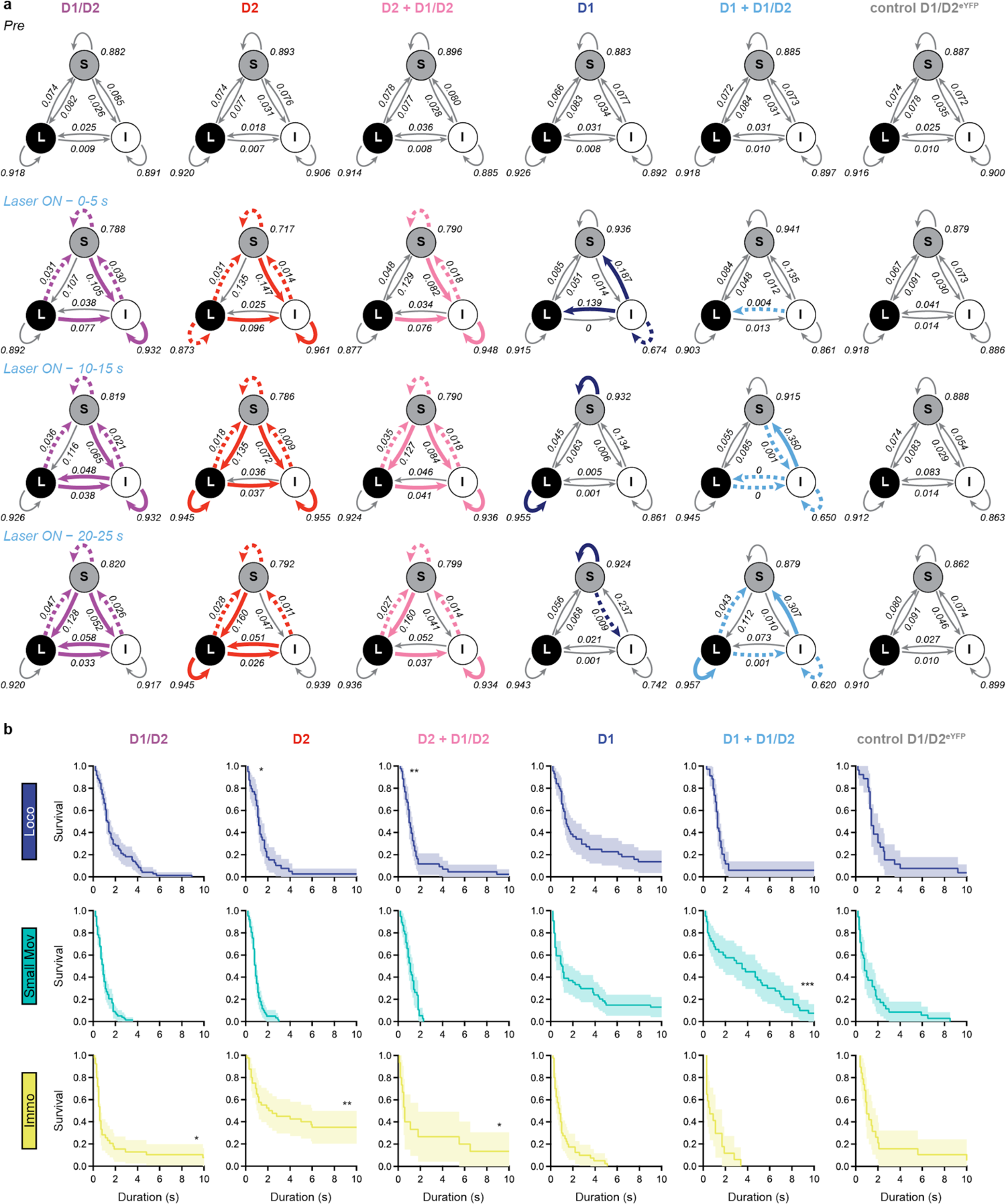
Temporal organization of motor states. **a**, Representation of transition probabilities between the three motor states during baseline (top) and during three periods occurring early (0-5 s), in the middle (10-15 s) or at the end (20-25 s) of the laser illumination (bottom) of D1/D2-SPNs (n = 21 mice), D2-SPNs (n = 15 mice), and D2+D1/D2-SPNs (n = 11 mice), D1-SPNs (n = 16 mice), D1+D1/D2-SPNs (n = 10 mice), and control D1/D2eYFP-SPNs (n = 11 mice). For Laser ON condition, colored arrows indicate significant increases (solid lines) or decreases (dashed lines) in transition probabilities in comparison with baseline condition. **b**, Survival curves of ongoing locomotion (top panels), small movements (middle panels), and immobility (bottom panels) episodes when light stimulation starts (D1/D2, n = 21; D2, n = 15; D2 + D1/D2, n = 11; D1, n = 16; D1 + D1/D2, n = 10; D1/D2eYFP, n = 11) (Kolmogorov-Smirnov test vs D1/D2eYFP: *p < 0.05, **p < 0.01, ***p < 0.001).

**Extended Data Figure 9.**
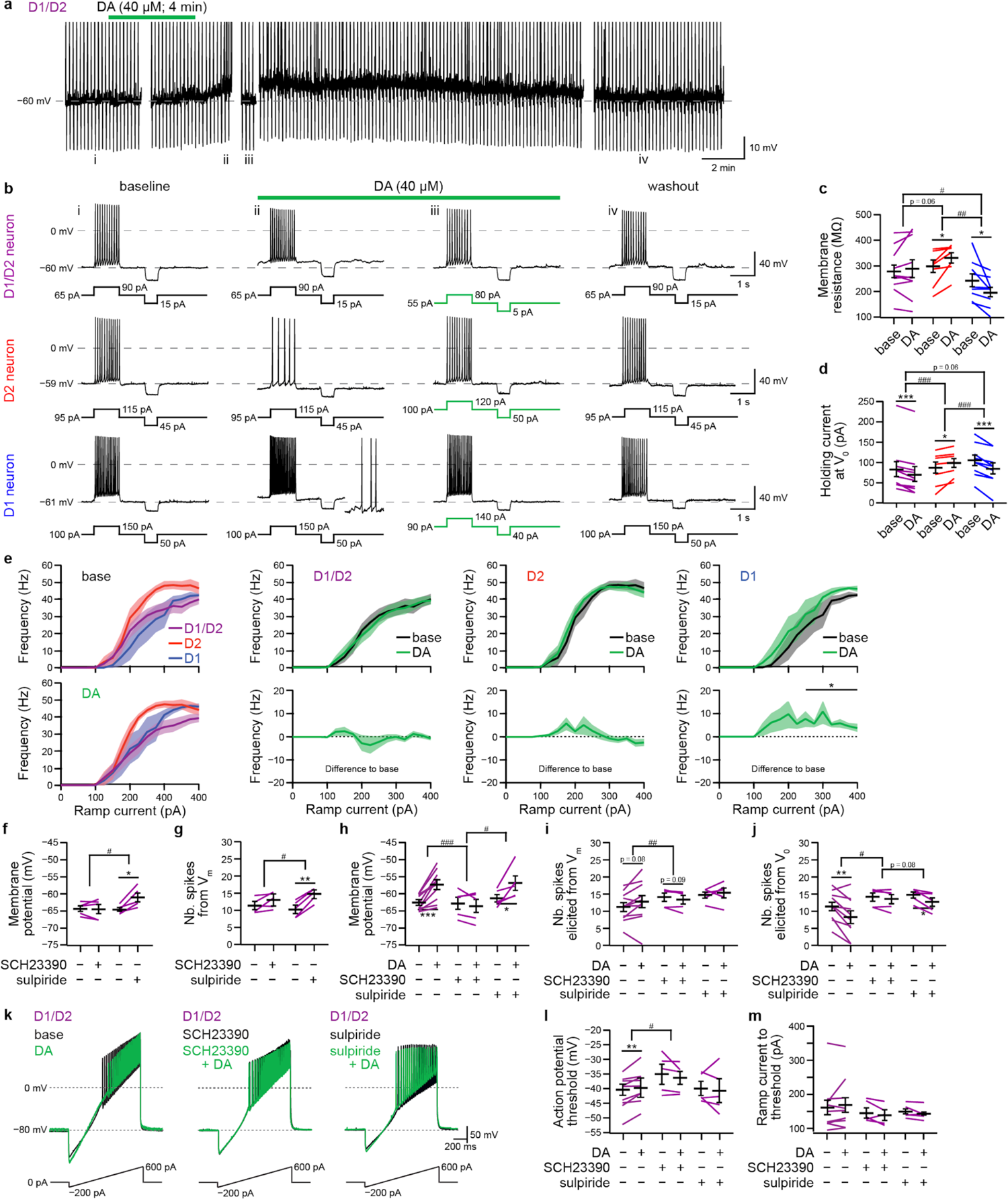
Effects of DA application on SPNs subpopulations. **a**, Representative example of the effect of bath applied DA (40 µM, 4 min) on membrane potential and evoked firing activity in one D1/D2-SPN. **b**, Voltage traces recorded in one representative D1/D2-SPN (top), one D1-SPN (middle), and one D2-SPN (bottom) in response to depolarizing and hyperpolarizing current steps during baseline (first column), at peak of DA effect from the same holding current than in the beginning of the recording (second column), at peak of DA from holding current to restore voltage at the start of the recording (third column), and after DA washout (last column). For the D1/D2-SPN example i-iv represent times indicated in **a**. **c-d**, Quantification of changes in membrane resistance (around –60 mV) induced by DA application (**c**), and change in holding necessary to maintain cells at the same potential than at the begging of the recording before and after DA application (**d**) in D1/D2 (n = 10 cells from 6 mice, violet), D1-(n = 9 cells from 4 mice, blue), and D2-SPNs (n = 7 cells from 4 mice, red) and (paired t-tests between baseline and after DA application: *p < 0.05, **p < 0.01, ***p < 0.001; t-tests comparing the magnitude of DA effect between SPN subpopulations: ^#^p < 0.05, ^##^p < 0.01, ^###^p < 0.001). **e**, Changes in average current-frequency relationship evaluated from ramp current injection during baseline and at the peak of DA effect in D1/D2- (n = 11 cells from 6 mice), D1- (n = 6 cells from 4 mice), and D2-SPNs (n = 8 cells from 4 mice) and (paired t-tests between baseline and after DA application: *p < 0.05). **f-g**, Effect of D1R antagonist (SCH23390, 10 µM) and D2R antagonist (sulpiride, 10 µM) application on membrane potential (**f**) and number of spikes elicited by a depolarizing current step (**g**) in D1/D2-SPNs (SCH23390, n = 5 cells from 3 mice; sulpiride, n = 5 cells from 3 mice) (paired t-tests between baseline and after antagonist application: *p < 0.05, **p < 0.01; t-tests comparing the magnitude of effect between treatments: # p < 0.05). **h-j**, Effect of DA application on D1/D2-SPNs without any pre-treatment, in the presence of SCH23390, or in the presence of sulpiride on the membrane potential (**h**), the number of elicited spiked by a depolarizing pulse from current membrane potential (**i**), and the number of spikes elicited from initial membrane potential (**j**) (no treatment, n = 11 cells from 6 mice; SCH23390, n = 5 cells from 3 mice; sulpiride, n = 5 cells from 3 mice) (paired t-tests between baseline and after DA application: *p < 0.05, **p < 0.01, ***p < 0.001; t-tests comparing the magnitude of DA effect between treatments: ^#^p < 0.05, ^##^p < 0.01, ^###^p < 0.001). **k**, Voltage response and APs of representative D1/D2-SPNs to ramp current injection before (black) and after (green) DA application without pre-treatment (left), in the presence of SCH23390 (middle), or in the presence of sulpiride (right). **l-m**, Effect of DA application on AP threshold (**l**) and minimal ramp current to elicit discharge (**m**) in D1/D2-SPNs without antagonists (n = 10 cells from 6 mice), or in the presence of SCH23390 (n = 5 cells from 3 mice), or sulpiride (n = 5 cells from 3 mice) (paired t-tests between baseline and after DA application: *p < 0.05, **p < 0.01; t-tests comparing the magnitude of DA effect between conditions: ^#^p < 0.05).

**Extended Data Figure 10.**
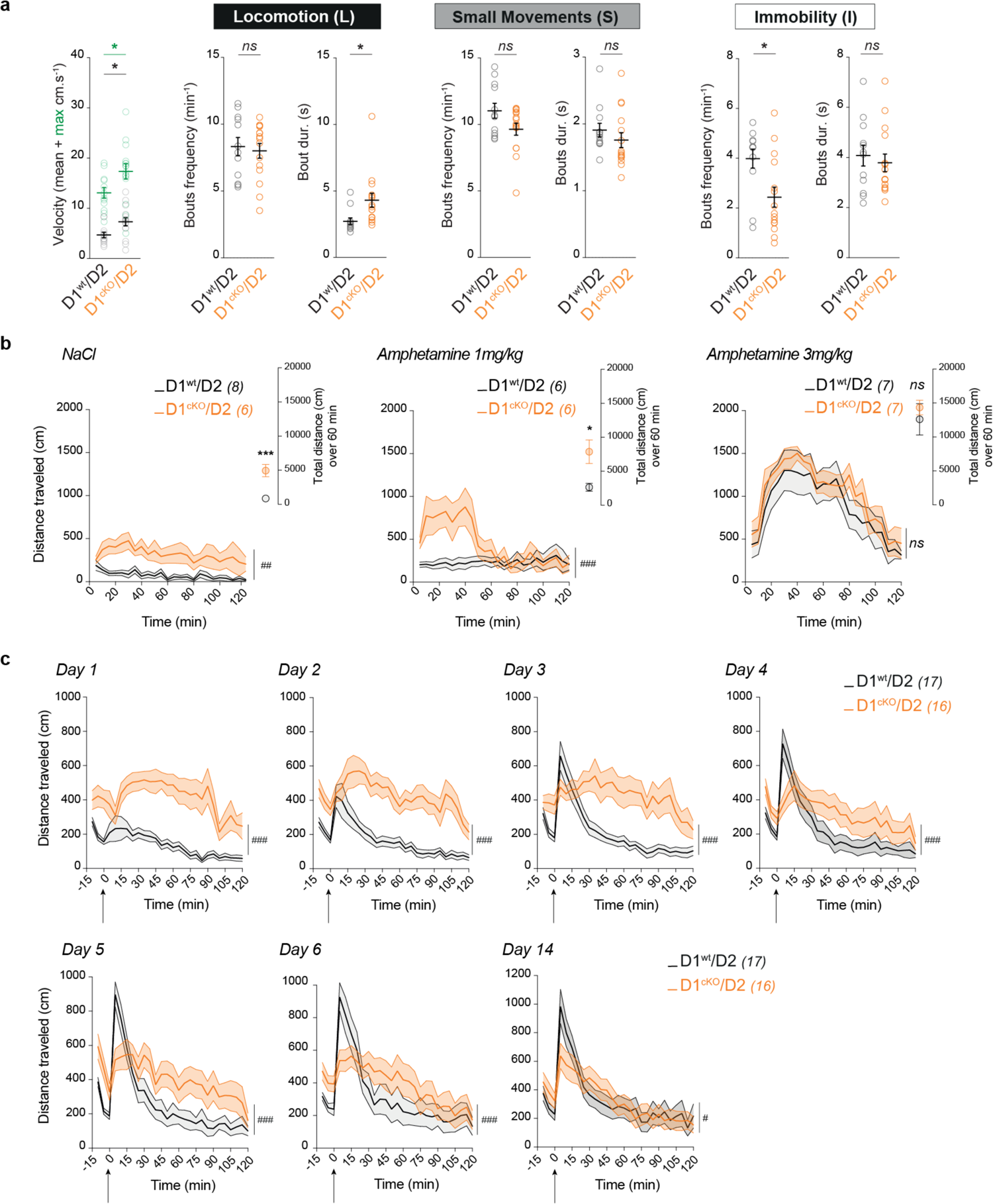
Supplements on the phenotypic characterization of D1cKO/D2 mice. **a**, Comparison of spontaneous locomotor behavior between D1^wt^/D2 (n = 13) and D1^cKO^/D2 (n = 15) mice in term of average and top velocity during locomotion episodes (left panel) and motor states architecture (D1^wt^/D2 vs. D1^cKO^/D2 mice: ns p > 0.05, *p < 0.05). **b**, Temporal distribution of the locomotor effects of acute amphetamine or saline injections in D1^wt^/D2 and D1^cKO^/D2 mice and (inserts) total distance traveled over the first 60 min after injection (D1^wt^/D2 vs. D1^cKO^/D2 mice: *p < 0.05, **p < 0.01, ***p < 0.001). **c**, Temporal evolution of traveled distance across repeated 10 mg/kg cocaine injection in D1^wt^/D2 (black) and D1^cKO^/D2 mice (orange) (D1^wt^/D2 vs. D1^cKO^/D2 mice: *p < 0.05, **p < 0.01, ***p < 0.001).

**Extended Data Table 1.** Complete list of electrophysiological properties recorded in D1/D2-, D1- and D2- SPNs. n: number of cells; NS: nonsignificant. Tukey post hoc tests: <, <<, <<<: inferior with p < 0.05, p < 0.01, p < 0.001, respectively; >, >>, >>>: superior with p < 0.05, p < 0.01, p < 0.001, respectively.

|  | D1/D2 neurons<br>n = 21 | D2 neurons<br>n = 19 | D1 neurons<br>n = 14 |
| --- | --- | --- | --- |
| Passive membrane properties |  |  |  |
| (1) Resting potential (mV)<br>ANOVA: NS ($F_{2,51} = 2.02$; $p = 0.14$) | -69.67 ± 0.77 | -71.05 ± 1.13 | -72.86 ± 1.49 |
| (2) Input Resistance (MΩ)<br>ANOVA: NS ($F_{2,51} = 0.83$; $p = 0.44$) | 110.13 ± 6.93 | 129.94 ± 12.43 | 115.81 ± 16.75 |
| (3) Membrane time constant (ms)<br>ANOVA: NS ($F_{2,51} = 2.55$; $p = 0.09$) | 6.48 ± 0.35 | 6.02 ± 0.39 | 5.16 ± 0.48 |
| (4) Membrane capacitance (pF)<br>ANOVA: NS ($F_{2,51} = 2.10$; $p = 0.13$) | 61.14 ± 3.40 | 50.20 ± 3.58 | 49.70 ± 4.98 |
| (5) Sag index (%)<br>ANOVA: NS ($F_{2,51} = 0.24$; $p = 0.79$) | 13.39 ± 1.02 | 13.83 ± 1.12 | 12.04 ± 1.63 |
| Just above threshold properties |  |  |  |
| (6) Step intensity to first spike (pA)<br>ANOVA: $p = 0.014$ ($F_{2,51} = 4.65$) | 123.33 ± 10.74<br>< D1 | 121.58 ± 11.68<br>< D1 | 176.43 ± 19.66<br>> D1/D2 . D2 |
| (7) Latency to first spike (ms)<br>ANOVA: NS ($F_{2,51} = 0.01$; $p = 0.99$) | 528.12 ± 53.15 | 521.17 ± 49.89 | 516.11 ± 66.61 |
| (8) Action potential threshold (mV)<br>ANOVA: $p < 0.001$ ($F_{2,51} = 15.03$) | -41.39 ± 1.05<br><<< D1 < D2 | -37.15 ± 1.11<br>> D1/D2 << D1 | -31.46 ± 1.70<br>>>> D1/D2 >> D2 |
| (9) Adaptation (Hz/s)<br>ANOVA: NS ($F_{2,51} = 0.58$; $p = 0.57$) | 4.46 ± 1.08 | 3.67 ± 0.61 | 3.06 ± 0.87 |
| (10) Minimal steady state frequency (Hz)<br>ANOVA: NS ($F_{2,51} = 2.00$; $p = 0.15$) | 6.96 ± 0.46 | 8.12 ± 0.49 | 8.25 ± 0.64 |
| Firing properties |  |  |  |
| (11) Early slope of current-frequency curve (Hz/pA)<br>ANOVA: $p < 0.001$ ($F_{2,51} = 1.55$) | 0.20 ± 0.01<br><< D2 | 0.25 ± 0.02<br>>>> D1 >> D1/D2 | 0.17 ± 0.01<br><<< D2 |
| (12) Maximum intensity before depolarization block (pA)<br>ANOVA: $p < 0.001$ ($F_{2,51} = 9.00$) | 666.67 ± 49.80<br>>> D2 | 470.00 ± 27.98<br><<< D1 << D1/D2 | 717.86 ± 46.77<br>>>> D2 |
| (13) Amplitude of early frequency adaptation (Hz)<br>ANOVA: NS ($F_{2,51} = 0.83$; $p = 0.44$) | 34.56 ± 3.15 | 28.64 ± 4.83 | 27.85 ± 4.33 |
| (14) Time constant of early frequency adaptation (ms)<br>ANOVA: $p = 0.006$ ($F_{2,51} = 5.60$) | 42.23 ± 4.34<br><< D2 | 67.53 ± 5.20<br>>> D1/D2 | 54.97 ± 8.10 |
| (15) Slope of late adaptation (Hz/s)<br>ANOVA: NS ($F_{2,51} = 1.66$; $p = 0.20$) | -14.09 ± 1.14 | -14.37 ± 1.60 | -18.20 ± 2.47 |
| (16) Maximal steady state frequency (Hz)<br>ANOVA: NS ($F_{2,51} = 0.75$; $p = 0.48$) | 40.95 ± 2.07 | 39.86 ± 2.53 | 43.81 ± 1.57 |
| (17) Spike frequency adaptation (%)<br>ANOVA: NS ($F_{2,51} = 1.34$; $p = 0.27$) | 97.70 ± 5.06 | 104.64 ± 5.50 | 92.45 ± 3.68 |
| Action potentials properties |  |  |  |
| (18) First spike amplitude (mV)<br>ANOVA: $p < 0.001$ ($F_{2,51} = 10.74$) | 95.12 ± 1.29<br>>>> D1 >> D2 | 87.14 ± 1.68<br><< D1/D2 | 84.03 ± 2.52<br><<< D1/D2 |
| (19) Second spike amplitude (mV)<br>ANOVA: $p = 0.049$ ($F_{2,51} = 3.08$) | 85.87 ± 1.68<br>> D1 | 82.87 ± 1.82 | 79.46 ± 1.76<br>< D1/D2 |
| (20) First spike duration (ms)<br>ANOVA: $p < 0.001$ ($F_{2,51} = 12.75$) | 3.01 ± 0.15<br>>>> D2 >> D1 | 2.22 ± 0.08<br><<< D1/D2 | 2.46 ± 0.09<br><< D1/D2 |
| (21) Second spike duration (ms)<br>ANOVA: $p < 0.001$ ($F_{2,51} = 10.14$) | 3.36 ± 0.17<br>>>> D2 > D1 | 2.57 ± 0.10<br><<< D1/D2 | 2.81 ± 0.09<br>< D1/D2 |
| (22) First spike half-width (ms)<br>ANOVA: $p < 0.001$ ($F_{2,51} = 13.36$) | 1.71 ± 0.09<br>>>> D2 >> D1 | 1.40 ± 0.06<br><<< D1/D2 | 1.38 ± 0.05<br><< D1/D2 |
| (23) Second spike half-width (ms)<br>ANOVA: $p < 0.001$ ($F_{2,51} = 9.94$) | 1.87 ± 0.10<br>>>> D2 > D1 | 1.40 ± 0.06<br><<< D1/D2 | 1.55 ± 0.05<br>< D1/D2 |
| (24) First spike AHP (mV)<br>ANOVA: $p < 0.001$ ($F_{2,51} = 10.37$) | -11.03 ± 0.91<br>>>> D2 > D1 | -16.77 ± 0.89<br><<< D1/D2 | -14.51 ± 1.04<br>< D1/D2 |
| (25) Second spike AHP (mV)<br>ANOVA: NS ($F_{2,51} = 0.931$; $p = 0.40$) | -14.35 ± 0.89 | -16.01 ± 0.77 | -15.28 ± 1.10 |
| (26) First spike AHP latency (ms)<br>ANOVA: $p < 0.001$ ($F_{2,51} = 8.38$) | 5.48 ± 0.26<br>>>> D2 > D1 | 4.29 ± 0.15<br><<< D1/D2 | 4.67 ± 0.20<br>< D1/D2 |
| (27) Second spike AHP latency (ms)<br>ANOVA: $p = 0.002$ ($F_{2,51} = 6.93$) | 8.44 ± 0.69<br>>> D2 > D1 | 5.97 ± 0.43<br><< D1/D2 | 5.97 ± 0.35<br>< D1/D2 |
| (28) Spike amplitude reduction (%)<br>ANOVA: $p = 0.034$ ($F_{2,51} = 3.61$) | 9.66 ± 1.47<br>> D2 . D1 | 4.86 ± 1.13<br>< D1/D2 | 4.92 ± 1.99<br>< D1/D2 |
| (29) Spike duration increase (%)<br>ANOVA: NS ($F_{2,51} = 1.69$; $p = 0.20$) | 12.00 ± 1.27 | 15.47 ± 1.59 | 14.71 ± 1.58 |
| (30) Spike half-width increase (%)<br>ANOVA: NS ($F_{2,51} = 1.51$; $p = 0.23$) | 8.92 ± 1.62 | 12.59 ± 2.00 | 13.06 ± 2.09 |

