## Supplemental for "Unexpected contributions of striatal projection neurons coexpressing dopamine D1 and D2 receptors in balancing motor control"

This PDF file includes:

Supplementary Text

Supplementary Figures 1 to 2

Supplementary Table 1

**Supplementary text:**

**Optogenetic stimulation paradigms in the dorsal striatum**

Optogenetic stimulation has rapidly established itself as a common method of brain stimulation allowing precise and cell-type specific control of neuronal activity *in vivo* in virtually all brain structures. Stimulation protocols should ideally be as close as possible to physiological neuronal activity. In this study, we applied constant light illumination, which has often been used for SPN opto-stimulations[1-5] and was originally applied in the princeps optogenetic study led by Kravitz and coauthors[6]. This protocol enables sustained neuronal depolarization, resulting in neurons firing at their own pace. Alternatively, applying patterned optogenetic manipulations would lead to highly synchronized neuronal activity, likely driving circuits outside their spontaneous physiological patterns. It can confound any causal inference about the circuit's functions[7]. Furthermore, in the dorsal striatum, SPNs organize into functional clusters, the firing of which is sequential and associated with specific behaviors or movements[8-12]. To assess whether different stimulation parameters lead to different behavioral outcomes, we conducted a comparative analysis of the effects on motor states of four different stimulation patterns. In addition to the 30 s constant light protocol, we investigated the effects over 30 s of three other protocols designed to reflect more physiological activation patterns observed in SPNs (Extended Data Fig. **4**):

- 5 Hz pulsed light for 30 s mimicking tonic activation at medium frequency;
- one pulse at 20 Hz for 500 ms mimicking a phasic transient activation;
- 20 Hz for 30 s mimicking long-lasting tonic activation at high frequency.

These stimulation protocols were applied to specifically activate D1/D2-SPNs, D1-SPNs, and D2-SPNs and to coactivate D1+D1/D2-SPNs and D2+D1/D2-SPNs in control mice.

We observed the following:

- one pulse at 20 Hz for 500 ms does not produce any notable change in motor states;
- the stimulation of D2+D1/D2-SPNs at 5 Hz or at 20 Hz for 30 s does not increase immobility;
- the stimulation of D2+D1/D2-SPNs at 5 Hz or at 20 Hz increases locomotion;
- the stimulation of D1+D1/D2-SPNs at 5 Hz does not increase locomotion.

The above effects do not fit with the canonical description of the prokinetic and antikinetic roles of the direct and indirect pathways, respectively[1,3,6]. In contrast, SPN stimulation with constant illumination fully recapitulated the respective pro-locomotion and pro-immobility effects of activating the direct pathway and the indirect pathway. Therefore, keeping in mind that patterned light stimulation would result in nonphysiological oversynchronization of SPNs, constant illumination appears to be the most reliable approach to probe the role of SPNs in motor behaviors.

However, the use of constant illumination is not without limitations. In particular, we do not control the actual firing activity of activated neurons. We performed *ex vivo* characterization of the firing activity elicited by continuous optogenetic activation during 30 s of D1/D2-SPNs, D1-SPNs and D2-SPNs (Extended Data Fig. **3f-i**). Despite ChR2 activation inducing a constant inward current in the recorded cells, the effect of this stimulation on firing activity is not constant during the 30 s of light. On average, after an initial peak at approximately 20-30 Hz lasting less than 1 s, the firing activity gradually decreases from ~ 20 Hz to ~ 10 Hz throughout the duration of the stimulation. This effect is most likely linked with the spike frequency adaptation reported among SPNs[13-14] that may rely on the activation/inactivation of some inwardly rectifying potassium currents and slow AHP currents[15-17]. Moreover, light-evoked neuronal activation varied greatly between two different neurons (Extended Data Fig. **3g**), but we did not observe differences between SPN subpopulations on average (Extended Data Fig. **3h-i**). All firing ceases immediately after light offset.

Furthermore, it may be tempting to establish correlations between the temporal dynamics of motor state changes and the temporal evolution of firing activity in response to light evaluated *ex vivo*. However, it would be highly speculative, as neuronal activity in brain slices loosely corresponds to what happens in an intact brain, as SPNs in slices are strongly hyperpolarized, most cortical and thalamic excitatory inputs are severed, and short-range and long-range neuronal networks are likely alleviated. Nevertheless, the progressive decrease in the amplitude of some motor effects could correspond to the *ex vivo* decrease in firing activity under continuous light: the facilitation of immobility under D1/D2, D2, and D2+D1/D2 stimulation; the inhibition of locomotion under D2+D1/D2 stimulation; or the facilitation of small movements under D1 stimulation (Fig. **4a-c**, Extended Data Fig. **7**). However, some motor effects persisted with the same magnitude throughout the whole stimulation period, such as the suppression of immobility under D1 and D1+D1/D2 stimulation or the suppression of small movements under D1/D2, D2, and D2+D1/D2 stimulation (Fig. **4a-c**, Extended Data Fig. **7**). In particular, for D2 or D2+D1/D2 stimulation, it could mean that, throughout the course of the stimulation, as ChR2-mediated activation of the indirect pathway weakens, the striatum becomes increasingly permissive to activity from the direct pathway, resulting in an increase in locomotion. According to this view, we should normally also observe a concomitant increase in small movements, which is not the case. In addition, there are more complex temporal dynamics, such as in the case of D1+D1/D2 stimulation characterized by a biphasic effect with an initial facilitation of small movements and a concomitant inhibition of locomotion, followed by a facilitation of locomotion concomitant with an inhibition of small movements (Fig. **4a-c**, Extended Data Fig. **7**). This transient change during the first 5 s might reflect the initial peak activation observed *ex vivo*. This would mean that strong activity in the direct pathway results in small movement promotion (including e.g., rearing, grooming, sniffing, and turning on itself) and that locomotion would require moderate activation of the direct pathway. However, we tested *in vivo* the effects of a pulsed stimulation at 20 Hz lasting 0.5 s, and as previously stated, this stimulation did not produce any notable change in motor states (Extended Data Fig. **4**). Eventually, the above correlative interpretations are highly speculative and would require additional experimental evidence to be adequately investigated.

**Comparison with previously reported effects of direct or indirect pathway activation**

A large part of the literature investigating the respective role of direct and indirect pathway SPNs in motor control converges to attribute a prokinetic function and an antikinetic function to the direct and indirect pathways, respectively[1,6,13-16]. However, these prokinetic or antikinetic functions can be translated into the motor phenotype through different means. For example, an antikinetic effect can be observed through a decrease in the time spent in ambulation or an increase in the time spent immobile, a decrease in ambulation or motor speed, or a decreased traveled distance. In our study, we observed that the stimulation of D1-D1/D2-SPNs (direct pathway) induces an increase in the time spent in locomotion at the expense of immobility and that the stimulation of D2+D1/D2-SPNs (indirect pathway) increases immobility at the expense of small movements without changes in the time spent in locomotion (Fig. **3**).

Going further, we also observed that D1+D1/D2-SPN activation transiently increases small movements during the first few seconds after stimulation onset. This observation cannot be directly compared with the seminal study using optogenetics[6,] as the first five seconds after stimulation onset were removed from their quantifications. In the same study[6], it was reported that direct pathway activation induces an ~ 4-fold increase in the time spent in ambulation, whereas we observed a milder ~ 30% increase in the time spent in locomotion with a similar D1+D1/D2 stimulation. This difference is most likely related to the circadian time at which the experiments were conducted. Indeed, mice are nocturnal animals, and we chose to work with a reversed light cycle, when mice are spontaneously more active. In our conditions, mice spend ~ 44% of the time engaged in locomotion during baseline (Fig. **3**), whereas Kravitz et al. reported an average of ~ 15% of time in ambulation[6,] as their mice were most likely manipulated during the light photoperiod. In addition to these effects on basal activity, the choice between dark or light photoperiod affects how optogenetic stimulations of the direct pathway are integrated (Supplementary Fig. **1**).

Moreover, we observed that the stimulation of D2+D1/D2-SPNs (indirect pathway) increases immobility at the expense of small movements without changes in the time spent in locomotion, whereas Kravitz et al. observed that the activation of D2R-expressing neurons (in a D2-cre mouse line) resulted in an increase in immobility at the expense of both small movements and ambulation[6]. This decrease in ambulation we did not replicate may arise from the coactivation, alongside D2-SPNs, of D2-expressing cholinergic interneurons. Indeed, the direct optogenetic activation of cholinergic interneurons inhibits locomotion and decreases velocity[17-18]. Alternatively, this difference may be linked with the difference in photoperiod during which experiments were conducted. In our conditions, during the dark photoperiod, mice are more prone to engage in locomotion, and the activation of the indirect pathway may not be sufficient to circumvent the strong spontaneous drive to engage in locomotion. Notwithstanding these considerations, our experiments demonstrate an antikinetic function of D2+D1/D2-SPNs through increased immobility and a decrease in small movements.

Furthermore, we observed that the stimulation of D1+D1/D2-SPNs decreased running velocity during locomotion episodes and that the stimulation of D2+D1/D2-SPNs increased running velocity during locomotion episodes (Fig. **3f-h**; Extended Data Fig. **5d-f**). We think these effects are well explained by the aversive or appetitive effects of direct pathway or indirect pathway activation[19-20], as demonstrated by the real-time place preference experiments (Extended Data Fig. **6**). However, the above effects may appear contradictory to some previous reports[1,3,21]. For example, Roseberry et al.[3] reported that in head-fixed animals on a trackball, stimulation of the direct pathway in inactive animals (speed at 0 cm/s) produces an increase in speed, thus inducing locomotion. Conversely, they reported that when the animal is running at approximately 10 cm/s, the stimulation of indirect pathway neurons decreases the animal's speed to values of approximately 5 cm/s, indicating that the animal is still moving and in a motor state we would classify as locomotion or small movements. However, there is no report on the effects of stimulating the direct pathway when the animal is moving or the effect of stimulating the indirect pathway when the animal is inactive. In a similar fashion, we examined the effects of optogenetic stimulations on mouse velocity by separating trials that occurred when mice were engaged in locomotion or engaged in immobility (Supplementary Fig. **2**). We observed similar results as those reported by Roseberry et al., namely, that the stimulation of D1-SPNs or D1+D1/D2-SPNs when the animal is in immobility resulted in an increase in velocity and that the stimulation of D2-SPNs, D2+D1/D2-SPNs, or D1/D2-SPNs when the animal is in locomotion resulted in a decrease in velocity. Reciprocally, we also found that the stimulation of D1-SPNs or D1+D1/D2-SPNs when the animal was in locomotion resulted in a decrease in velocity and that the stimulation of D2-SPNs, D2+D1/D2-SPNs, or D1/D2-SPNs when the animal was in immobility resulted in an increase in velocity. Importantly, these curves are not different from those obtained from control animals. This means that in our experimental setup (mice exploring a square open field at their own pace), these changes in velocity are mainly driven by the spontaneous alternations between motor states. These findings fit well with a previous observation[1] that, for mice exploring an open field, not all direct pathway stimulations resulted in locomotion initiation and that not all indirect pathway stimulations resulted in immobility initiation. Altogether, these results clearly indicate that the ongoing state of the animal is a critical determinant of the efficiency of activation of either pathway.

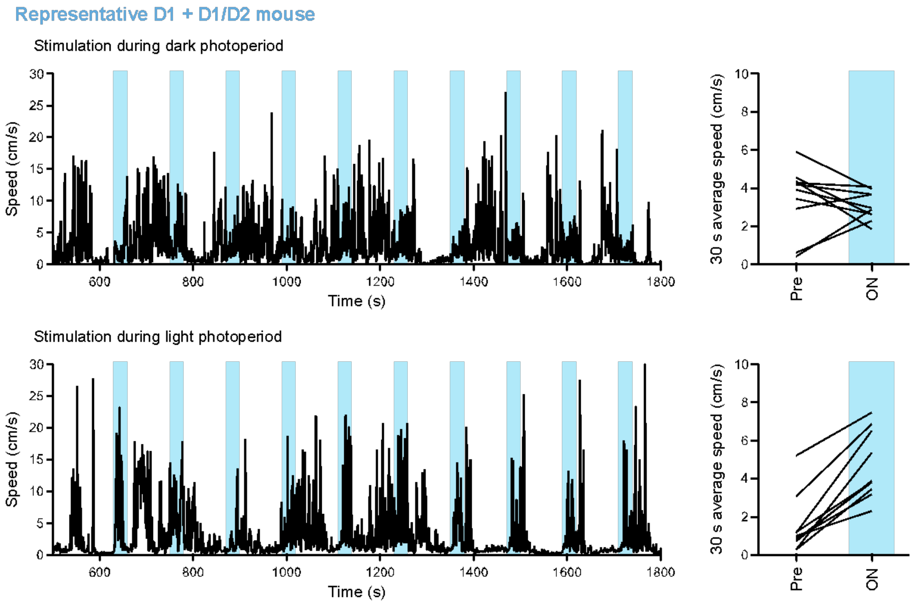


**Supplementary Fig. 1: Impact of the nocturnal and diurnal phases on open field speed of mice in response to direct pathway stimulation. Left**, Representative effects of mixed D1+D1/D2-SPNs stimulations on velocity during dark or light photoperiod recorded in the same mouse. **Right**, Graphs representing the resulting averaged speed calculated during 30 s before and during the 10 consecutive stimulation trials.

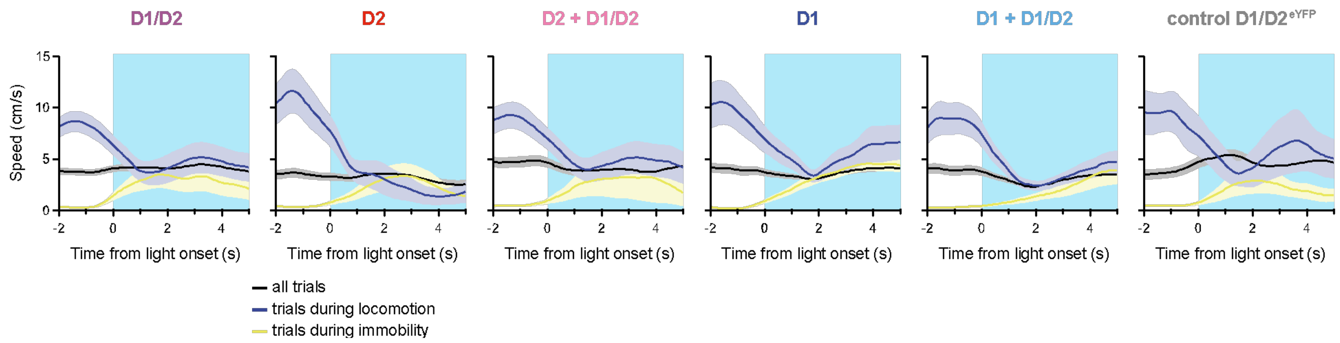


**Supplementary Fig. 2: Temporal evolution of speed around stimulation onset.** Optgenetic activation trials were separated according to the motor state of the animal when the stimulation started: in blue, trials occurring during locomotion; in yellow, trials occurring during immobility; in black, average for all trials. The same temporal evolution of speed is observed for the optogenetic activation of SPNs subpopulations and for control animals.

### Supplementary Table 1: Detailed statistical analysis for Figures, Extended Data Figures, and Extended Table

| Figure number | n | Statistical method | F/t value | p value | Post hoc multiple comparisons |
|---|---|---|---|---|---|
| Figure 1f | D1/D2: 8 cells<br>D1: 8 cells<br>D2: 8 cells | 2-way ANOVA with one factor replication | Geno F(2,21)=0.14<br>Radius F(203,4263)=143.6<br>Geno x radius F(206,4263)=1.79 | Geno P=0.8691<br>Radius P<0.0001<br>Geno x radius P<0.0001 | Comparisons between each pairs of genotypes for all radius bins<br>D1 vs D2: P>0.05 all bins<br>D1 vs D1/D2: P<0.05 between 31μm and 51μm<br>D2 vs D1/D2: P>0.05 |
| Figure 1g | D1/D2: 8 cells<br>D1: 8 cells<br>D2: 8 cells | One way ANOVA | F(2,21)=3.86 | P=0.0374 | D1 vs D2: P=0.3858<br>D1 vs D1/D2: P=0.3443<br>D2 vs D1/D2: P=0.0292 |
| Extended Data Figure 1b | 10 stions per mouse<br>3 mice | Friedman test | $X^2(10)$= 11.29 | P=0.256 | |
| Figure 2g-k Extended Data Table 1 | D1/D2: 21 cells<br>D1: 14 cells<br>D2: 19 cells | One-way ANOVA | F(2,51)<br>See Extended Data Table 1 | See Extended Data Table 1 | See Extended Data Table 1 |
| Figure 3c | D1/D2: 21 mice<br>D2: 16 mice<br>D2 + D1/D2: 13 mice<br>D1: 18 mice<br>D1 + D1/D2: 16 mice<br>D1/D2$^{ctl}$: 11 mice | 2-way ANOVA with one factor replication | Ipsilateral rotations<br>Stim x Genotype<br>F(10,178)=40.85<br>Stim<br>F(2,178)=186.2<br>Genotype<br>F(5,89)=32.6<br>Contralateral rotations<br>Stim x Genotype<br>F(10,178)=39.8<br>Stim<br>F(2,178)=43.49<br>Genotype<br>F(5,89)=20.26 | Stim x Genotype<br>P<0.0001<br>Stim<br>P<0.0001<br>Genotype<br>P<0.0001 | Ipsilateral rotations:<br>Pre vs Stim \| Pre vs Post \| Stim vs Post:<br>D1/D2: P<0.0001 \| P=0.931 \| P<0.0001<br>D2: P<0.0001 \| P=0.730\| P<0.0001<br>D2 + D1/D2: P<0.0001 \| P=0.971 \| P<0.0001<br>D1/D2$^{ctl}$: P=0.985 \| P=0.996 \| P=0.996<br>Contralateral rotations:<br>Pre vs Stim \| Pre vs Post \| Stim vs Post:<br>D1: P<0.0001 \| P=0.933 \| P<0.0001<br>D1 + D1/D2: P<0.0001 \| P>0.999 \| P<0.0001<br>LaserON vs control D1/D2$^{eYFP}$:<br>Ipsilateral D1/D2: P<0.0001<br>Ipsilateral D2: P<0.0001<br>Ipsilateral D2+D1/D2: P<0.0001<br>Contralateral D1: P<0.0001<br>Contralateral D1+D1/D2: P<0.0001 |
| Figure 3e Extended Data Figure 4a | protocol constant light:<br>D1/D2: 21 mice<br>D2: 15 mice<br>D2+D1/D2: 11 mice<br>D1: 16 mice<br>D1+D1/D2: 10 mice<br>D1/D2$^{ctl}$: 11 mice<br><br>protocol 5 Hz continuous:<br>D1/D2: 8 mice<br>D2: 8 mice<br>D2+D1/D2: 6 mice<br>D1: 6 mice<br>D1+D1/D2: 5 mice<br>D1/D2$^{ctl}$: 5 mice<br><br>protocol 20 Hz one pulse 0.5 s:<br>D1/D2: 8 mice<br>D2: 8 mice<br>D2+D1/D2: 6 mice<br>D1: 6 mice<br>D1+D1/D2: 5 mice<br>D1/D2$^{ctl}$: 5 mice<br><br>protocol 20 Hz continuous:<br>D1/D2: 13 mice<br>D2: 8 mice<br>D2+D1/D2: 5 mice<br>D1: 11 mice<br>D1+D1/D2: 5 mice<br>D1/D2$^{ctl}$: 6 mice | 3-way ANOVA with one factor replication | Locomotion<br>Genotype F(5,173)=3.16<br>Stim F(1,173)=0.20<br>Protocol F(3,173)=1.98<br>Geno x Stim<br>F(5,173)=2.95<br>Geno x Proto<br>F(15,173)=1.87<br>Stim x Proto<br>F(3,173)=1.52<br>Geno x Stim x Proto<br>F(15,173)=1.23<br><br>Small movements<br>Genotype F(5,173)=2.20<br>Stim F(1,173)=12.16<br>Protocol F(3,173)=5.95<br>Geno x Stim<br>F(5,173)=6.80<br>Geno x Proto<br>F(15,173)=4.03<br>Stim x Proto<br>F(3,173)=12.04<br>Geno x Stim x Proto<br>F(15,173)=4.25<br><br>Immobility<br>Genotype F(5,173)=3.89<br>Stim F(1,173)=16.17<br>Protocol F(3,173)=3.94<br>Geno x Stim<br>F(5,173)=12.30<br>Geno x Proto<br>F(15,173)=2.80<br>Stim x Proto<br>F(3,173)=5.36<br>Geno x Stim x Proto<br>F(15,173)=5.94 | Locomotion<br>Genotype P=0.0094<br>Stim P=0.6591<br>Protocol P=0.1182<br>Geno x Stim<br>P=0.0139<br>Geno x Proto<br>P=0.0297<br>Stim x Proto<br>P=0.2123<br>Geno x Stim x Proto<br>P=0.2530<br><br>Small movements<br>Genotype P=0.0568<br>Stim P=0.0006<br>Protocol P=0.0007<br>Geno x Stim<br>P<0.0001<br>Geno x Proto<br>P<0.0001<br>Stim x Proto<br>P<0.0001<br>Geno x Stim x Proto<br>P<0.0001<br><br>Immobility<br>Genotype P=0.00023<br>Stim P<0.0001<br>Protocol P=0.0095<br>Geno x Stim<br>P<0.0001<br>Geno x Proto<br>P=0.0005<br>Stim x Proto<br>P=0.0015<br>Geno x Stim x Proto<br>P<0.0001 | Locomotion:<br>Pre vs. Laser ON for each protocol<br>constant light \| 5 Hz cont \| 20 Hz one pulse \| 20 Hz cont<br>D1/D2: P=0.228 \| P=0.069 \| P=0.113 \| P=0.652<br>D2: P=0.430 \| P=0.565 \| P=0.112 \| P=0.776<br>D2+D1/D2: P=0.999 \| P=0.0089 \| P=0.971 \| P=0.608<br>D1: P=0.198 \| P=0.066 \| P=0.293 \| P=0.118<br>D1+D1/D2: P=0.0039 \| P=0.113 \| P=0.335 \| P=0.170<br>D1/D2$^{ctl}$: P=0.733 \| P=0.227 \| P=0.406 \| P=0.705<br><br>Small movements:<br>Pre vs. Laser ON for each protocol<br>constant light \| 5 Hz cont \| 20 Hz one pulse \| 20 Hz cont<br>D1/D2: P<0.0001 \| P=0.355 \| P=0.270 \| P=0.637<br>D2: P<0.0001 \| P<0.0001 \| P=0.432 \| P=0.855<br>D2+D1/D2: P<0.0001 \| P<0.0001 \| P=0.301 \| P=0.554<br>D1: P<0.0001 \| P=0.758 \| P=0.423 \| P=0.254<br>D1+D1/D2: P=0.849 \| P=0.012 \| P=0.264 \| P=0.134<br>D1/D2$^{ctl}$: P=0.307 \| P=0.804 \| P=0.906 \| P=0.546<br><br>Immobility:<br>Pre vs. Laser ON for each protocol<br>constant light \| 5 Hz cont \| 20 Hz one pulse \| 20 Hz cont<br>D1/D2: P<0.0001 \| P=0.0304 \| P=0.366 \| P=0.275<br>D2: P<0.0001 \| P<0.0001 \| P=0.374 \| P=0.899<br>D2+D1/D2: P<0.0001 \| P=0.496 \| P=0.285 \| P=0.931<br>D1: P<0.0001 \| P=0.066 \| P=0.786 \| P=0.586<br>D1+D1/D2: <0.0001 \| P=0.800 \| P=0.0305 \| P=0.894<br>D1/D2$^{ctl}$: P=0.740 \| P=0.354 \| P=0.312 \| P=0.793 |
| Figure 3e Extended Data Figure 5a (left) Locomotion Total duration | D1/D2: 21 mice<br>D2: 15 mice<br>D2+D1/D2: 11 mice<br>D1: 16 mice<br>D1+D1/D2: 10 mice<br>D1/D2$^{ctl}$: 11 mice | 2-way ANOVA with one factor replication | Stim x Genotype<br>F(10,156)=4.033<br>Stim<br>F(2,156)=2.752<br>Genotype<br>F(5,78)=3.468 | Stim x Genotype<br>P<0.0001<br>Stim<br>P=0.067<br>Genotype<br>P=0.007 | Pre vs LaserON \| Pre vs LaserOFF \| LaserON vs LaserOFF :<br>D1/D2: P=0.971 \| P=0.0046 \| P=0.812<br>D2: P=0.999 \| P=0.963 \| P>0.999<br>D2+D1/D2: P>0.999 \| P=0.0256 \| P=0.0256<br>D1: P=0.999 \| P=0.999 \| P=0.586<br>D1+D1/D2: P=0.021 \| P>0.999 \| P=0.170 |

| | | | | | |
|---|---|---|---|---|---|
| | | | | | D1/D2ctl: P=0.970 \| P>0.999 \| P=0.417<br>LaserON vs control D1/D2eYFP:<br>D1/D2: P=0.628<br>D2: P=0.474<br>D2+D1/D2: P=0.979<br>D1: P=0.952<br>D1+D1/D2: P=0.071 |
| Figure 3e Extended Data Figure 5a (middle) Locomotion Bout number | D1/D2: 21 mice<br>D2: 15 mice<br>D2+D1/D2: 11 mice<br>D1: 16 mice<br>D1+D1/D2: 10 mice<br>D1/D2ctl: 11 mice | 2-way ANOVA with one factor replication | Stim x Genotype<br>F(10,156)=5.96<br>Stim<br>F(2,156)=16.34<br>Genotype<br>F(5,78)=3.752 | Stim x Genotype<br>P<0.0001<br>Stim<br>P<0.0001<br>Genotype<br>P=0.0353 | Pre vs LaserON \| Pre vs LaserOFF \| LaserON vs LaserOFF :<br>D1/D2: P=0.314 \| P=0.037 \| P=0.563<br>D2: P=0.0004 \| P=0.0500 \| P=0.237<br>D2+D1/D2: P=0.424 \| P=0.612\| P=0.950<br>D1: P<0.0001 \| P=0.589 \| P<0.0001<br>D1+D1/D2: P=0.0001 \| P=0.629 \| P<0.0001<br>D1/D2ctl: P=0.259 \| P=0.963 \| P=0.160<br>LaserON vs control D1/D2eYFP:<br>D1/D2: P=0.248<br>D2: P<0.0001<br>D2+D1/D2: P=0.180<br>D1: P<0.0001<br>D1+D1/D2: P=0.006 |
| Figure 3e Extended Data Figure 5a (right) Locomotion Bout mean duration | D1/D2: 21 mice<br>D2: 15 mice<br>D2+D1/D2: 11 mice<br>D1: 16 mice<br>D1+D1/D2: 10 mice<br>D1/D2ctl: 11 mice | 2-way ANOVA with one factor replication | Stim x Genotype<br>F(10,156)=11.03<br>Stim<br>F(2,156)=23.13<br>Genotype<br>F(5,78)=7.635 | Stim x Genotype<br>P<0.0001<br>Stim<br>P<0.0001<br>Genotype<br>P<0.0001 | Pre vs LaserON \| Pre vs LaserOFF \| LaserON vs LaserOFF :<br>D1/D2: P=0.721 \| P=0.172 \| P=0.559<br>D2: P=0.117 \| P=0.960 \| P=0.064<br>D2+D1/D2: P=0.950 \| P=0.0003 \| P=0.0010<br>D1: P<0.0001 \| P=0.162 \| P<0.0001<br>D1+D1/D2: P<0.0001 \| P=0.994 \| P<0.00001<br>D1/D2ctl: P=0.999 \| P=0.696 \| P=0.725<br>LaserON vs control D1/D2eYFP:<br>D1/D2: P=0.999<br>D2: P=0.028<br>D2+D1/D2: P=0.939<br>D1: P<0.0001<br>D1+D1/D2: P<0.0001 |
| Figure 3e Extended Data Figure 5a (left) Small movements Total duration | D1/D2: 21 mice<br>D2: 15 mice<br>D2+D1/D2: 11 mice<br>D1: 16 mice<br>D1+D1/D2: 10 mice<br>D1/D2ctl: 11 mice | 2-way ANOVA with one factor replication | Stim x Genotype<br>F(10,156)=44.6<br>Stim<br>F(2,156)=122.7<br>Genotype<br>F(5,78)=6.613 | Stim x Genotype<br>P<0.0001<br>Stim<br>P<0.0001<br>Genotype<br>P<0.0001 | Pre vs LaserON \| Pre vs LaserOFF \| LaserON vs LaserOFF :<br>D1/D2: P<0.0001 \| P=0.896 \| P<0.0001<br>D2: P<0.0001 \| P=0.791 \| P<0.0001<br>D2+D1/D2: P<0.0001 \| P=0.002 \| P<0.0001<br>D1: P<0.0001 \| P=0.742 \| P<0.0001<br>D1+D1/D2: P=0.977 \| P=0.780 \| P=0.656<br>D1/D2ctl: P=0.214 \| P=0.917 \| P=0.405<br>LaserON vs control D1/D2eYFP:<br>D1/D2: P<0.0001<br>D2: P<0.0001<br>D2+D1/D2: P<0.0001<br>D1: P=0.012<br>D1+D1/D2: P=0.955 |
| Figure 3e Extended Data Figure 5a (middle) Small movements Bout number | D1/D2: 21 mice<br>D2: 15 mice<br>D2+D1/D2: 11 mice<br>D1: 16 mice<br>D1+D1/D2: 10 mice<br>D1/D2ctl: 11 mice | 2-way ANOVA with one factor replication | Stim x Genotype<br>F(10,156)=12.44<br>Stim<br>F(2,156)=240.9<br>Genotype<br>F(5,78)=10.89 | Stim x Genotype<br>P<0.0001<br>Stim<br>P<0.0001<br>Genotype<br>P<0.0001 | Pre vs LaserON \| Pre vs LaserOFF \| LaserON vs LaserOFF :<br>D1/D2: P<0.0001 \| P=0.697 \| P<0.0001<br>D2: P<0.0001 \| P=0.0158 \| P<0.0001<br>D2+D1/D2: P<0.0001 \| P=0.0189 \| P<0.0001<br>D1: P<0.0001 \| P=0.785 \| P<0.0001<br>D1+D1/D2: P<0.0001 \| P=0.947 \| P<0.0001<br>D1/D2ctl: P=0.974 \| P=0.948 \| P=0.996<br>LaserON vs control D1/D2eYFP:<br>D1/D2: P<0.0001<br>D2: P<0.0001<br>D2+D1/D2: P<0.0001<br>D1: P<0.0001<br>D1+D1/D2: P<0.0001 |
| Figure 3e Extended Data Figure 5a (right) Small movements Bout mean duration | D1/D2: 21 mice<br>D2: 15 mice<br>D2+D1/D2: 11 mice<br>D1: 16 mice<br>D1+D1/D2: 10 mice<br>D1/D2ctl: 11 mice | 2-way ANOVA with one factor replication | Stim x Genotype<br>F(10,156)=45.86<br>Stim<br>F(2,156)=4.015<br>Genotype<br>F(5,78)=6.207 | Stim x Genotype<br>P<0.0001<br>Stim<br>P=0.0199<br>Genotype<br>P<0.0001 | Pre vs LaserON \| Pre vs LaserOFF \| LaserON vs LaserOFF :<br>D1/D2: P<0.0001 \| P=0.679 \| P<0.0001<br>D2: P<0.0001 \| P=0.184 \| P<0.0001<br>D2+D1/D2: P<0.0001 \| P=0.524 \| P<0.0001<br>D1: P<0.0001 \| P=0.501 \| P<0.0001<br>D1+D1/D2: P<0.0001 \| P=0.642 \| P<0.0001<br>D1/D2ctl: P=0.159 \| P=0.930 \| P=0.304<br>LaserON vs control D1/D2eYFP:<br>D1/D2: P=0.018<br>D2: P=0.0002<br>D2+D1/D2: P=0.012<br>D1: P<0.0001<br>D1+D1/D2: P<0.0001 |
| Figure 3e Extended Data Figure 5a (left) Immobility Total duration | D1/D2: 21 mice<br>D2: 15 mice<br>D2+D1/D2: 11 mice<br>D1: 16 mice<br>D1+D1/D2: 10 mice<br>D1/D2ctl: 11 mice | 2-way ANOVA with one factor replication | Stim x Genotype<br>F(10,156)=33.33<br>Stim<br>F(2,156)=33.84<br>Genotype<br>F(5,78)=10.43 | Stim x Genotype<br>P<0.0001<br>Stim<br>P<0.0001<br>Genotype<br>P<0.0001 | Pre vs LaserON \| Pre vs LaserOFF \| LaserON vs LaserOFF :<br>D1/D2: P<0.0001 \| P=0.0067 \| P<0.0001<br>D2: P<0.0001 \| P=0.121\| P<0.0001<br>D2+D1/D2: P<0.0001 \| P=0.254 \| P<0.0001<br>D1: P<0.0001 \| P=0.0133 \| P<0.0001<br>D1+D1/D2: P=0.0006 \| P=0.943\| P=0.002<br>D1/D2ctl: P=0.932 \| P=0.454 \| P=0.266<br>LaserON vs control D1/D2eYFP: |

| | | | | | |
|---|---|---|---|---|---|
| | | | | | D1/D2: P<0.0001<br>D2: P<0.0001<br>D2+D1/D2: P<0.0001<br>D1: P=0.015<br>D1+D1/D2: P=0.012 |
| Figure 3e<br>Extended<br>Data Figure<br>5a (middle)<br>Immobility<br>Bout number | D1/D2: 21 mice<br>D2: 15 mice<br>D2+D1/D2: 11 mice<br>D1: 16 mice<br>D1+D1/D2: 10 mice<br>D1/D2$^{ctl}$: 11 mice | 2-way ANOVA<br>with one factor<br>replication | Stim x Genotype<br>F(10,156)=33.79<br>Stim<br>F(2,156)=6.168<br>Genotype<br>F(5,78)=11.15 | Stim x Genotype<br>P<0.0001<br>Stim<br>P=0.0026<br>Genotype<br>P<0.0001 | Pre vs LaserON \| Pre vs LaserOFF \| LaserON vs LaserOFF :<br>D1/D2: P<0.0001 \| P=0.201 \| P<0.0001<br>D2: P<0.0001 \| P=0.835 \| P<0.0001<br>D2+D1/D2: P<0.0001 \| P=0.0003 \| P<0.0001<br>D1: P<0.0001 \| P=0.283 \| P<0.0001<br>D1+D1/D2: P<0.0001 \| P=0.826 \| P<0.0001<br>D1/D2$^{ctl}$: P=0.962 \| P=0.989 \| P=0.992<br>LaserON vs control D1/D2$^{eYFP}$:<br>D1/D2: P<0.0001<br>D2: P=0.082<br>D2+D1/D2: P=0.0002<br>D1: P<0.0001<br>D1+D1/D2: P<0.0001 |
| Figure 3e<br>Extended<br>Data Figure<br>5a (right)<br>Immobility<br>Bout mean<br>duration | D1/D2: 21 mice<br>D2: 15 mice<br>D2+D1/D2: 11 mice<br>D1: 16 mice<br>D1+D1/D2: 10 mice<br>D1/D2$^{ctl}$: 11 mice | 2-way ANOVA<br>with one factor<br>replication | Stim x Genotype<br>F(10,156)=9.171<br>Stim<br>F(2,156)=7.041<br>Genotype<br>F(5,78)=5.98 | Stim x Genotype<br>P<0.0001<br>Stim<br>P=0.0012<br>Genotype<br>P<0.0001 | Pre vs LaserON \| Pre vs LaserOFF \| LaserON vs LaserOFF :<br>D1/D2: P=0.0004 \| P=0.078 \| P=0.198<br>D2: P<0.0001 \| P=0.142 \| P<0.0001<br>D2+D1/D2: P=0.0011 \| P=0.653 \| P=0.0182<br>D1: P=0.044 \| P=0.512 \| P=0.0016<br>D1+D1/D2: P=0.088 \| P=0.991 \| P=0.066<br>D1/D2$^{ctl}$: P=0.823 \| P=0.504 \| P=0.204<br>LaserON vs control D1/D2$^{eYFP}$:<br>D1/D2: P=0.020<br>D2: P<0.0001<br>D2+D1/D2: P=0.016<br>D1: P=0.607<br>D1+D1/D2: P=0.476 |
| Figure 3h<br>Extended<br>Data Figure<br>5b<br>Averaged<br>velocity<br>(black) | D1/D2: 21 mice<br>D2: 15 mice<br>D2+D1/D2: 11 mice<br>D1: 16 mice<br>D1+D1/D2: 10 mice<br>D1/D2$^{ctl}$: 11 mice | 2-way ANOVA<br>with one factor<br>replication | Stim x Genotype<br>F(10,156)=26.52<br>Stim<br>F(2,156)=21.97<br>Genotype<br>F(5,78)=9.121 | Stim x Genotype<br>P<0.0001<br>Stim<br>P<0.0001<br>Genotype<br>P<0.0001 | Pre vs LaserON \| Pre vs LaserOFF \| LaserON vs LaserOFF :<br>D1/D2: P<0.0001 \| P=0.171 \| P<0.0001<br>D2: P<0.0001 \| P=0.199 \| P<0.0001<br>D2+D1/D2: P<0.0001 \| P=0.079 \| P<0.0001<br>D1: P<0.0001 \| P=0.060 \| P<0.0001<br>D1+D1/D2: P<0.0001\| P=0.540 \| P=0.0050<br>D1/D2$^{ctl}$: P=0.963 \| P=0.613 \| P=0.453<br>LaserON vs control D1/D2$^{eYFP}$:<br>D1/D2: P=0.002<br>D2: P<0.0001<br>D2+D1/D2: P<0.0001<br>D1: P=0.001<br>D1+D1/D2: P=0.0003 |
| Figure 3h<br>Extended<br>Data Figure<br>5b<br>Top velocity<br>(green) | D1/D2: 21 mice<br>D2: 15 mice<br>D2+D1/D2: 11 mice<br>D1: 16 mice<br>D1+D1/D2: 10 mice<br>D1/D2$^{ctl}$: 11 mice | 2-way ANOVA<br>with one factor<br>replication | Stim x Genotype<br>F(10,156)=21.42<br>Stim<br>F(2,156)=6.872<br>Genotype<br>F(5,78)=7.395 | Stim x Genotype<br>P<0.0001<br>Stim<br>P=0.0014<br>Genotype<br>P<0.0001 | Pre vs LaserON \| Pre vs LaserOFF \| LaserON vs LaserOFF :<br>D1/D2: P<0.0001 \| P=0.053 \| P<0.0001<br>D2: P<0.0001 \| P=0.099 \| P<0.0001<br>D2+D1/D2: P<0.0001 \| P=0.084\| P=0.0247<br>D1: P<0.0001 \| P=0.0134 \| P<0.0001<br>D1+D1/D2: P<0.0001\| P=0.297 \| P=0.0015<br>D1/D2$^{ctl}$: P=0.677 \| P=0.358 \| P=0.857<br>LaserON vs control D1/D2$^{eYFP}$:<br>D1/D2: P=0.016<br>D2: P=0.0003<br>D2+D1/D2: P=0.006<br>D1: P=0.0009<br>D1+D1/D2: P=0.0002 |
| Extended<br>Data Figure<br>5c (left) | D1/D2: 21 mice<br>D2: 15 mice<br>D2+D1/D2: 11 mice<br>D1: 16 mice<br>D1+D1/D2: 10 mice<br>D1/D2$^{ctl}$: 11 mice | 3-way ANOVA<br>with two factors<br>(time, stim)<br>replication | Genotype F(5,79)=5.194<br>Time F(60,4740)=1244.0<br>Geno x time<br>F(300,4740)=10.24<br>Stim F(2,158)=7.270<br>Geno x stim<br>F(10,158)=10.23<br>Time x stim<br>F(120,9480)=41.91<br>Geno x time x stim<br>F(600,9480)=16.91 | Genotype P=0.00036<br>Time P=1.5e-144<br>Geno x time P=1.0e-18<br>Stim P=0.0031<br>Geno x stim P=3.4e-10<br>Time x stim P=3.2e-46<br>Geno x time x stim<br>P=1.5e-64 | Difference with D1/D2<br>Time points: 1.1s/1.2s/1.3s<br>D1: p<0.0001/ p<0.0001 / p<0.0001<br>D2: p=0.040 / p=0.036 / p=0.029<br>D1 + D1/D2: p<0.0001/ p<0.0001 / p<0.0001<br>D2 + D1/D2: p=0.306/ p=0.259 / p=0.205<br>D1/D2$^{ctl}$: p<0.0001/ p=0.0002 / p=0.0003 |
| Extended<br>Data Figure<br>5c (right) | D1/D2: 21 mice<br>D2: 15 mice<br>D2+D1/D2: 11 mice<br>D1: 16 mice<br>D1+D1/D2: 10 mice<br>D1/D2$^{ctl}$: 11 mice | 3-way ANOVA<br>with two factors<br>(time, stim)<br>replication | Genotype F(5,79)=2.55<br>Time F(59,4611)=875.9<br>Geno x time<br>F(295,4611)=7.05<br>Stim F(2,158)=43.81<br>Geno x stim<br>F(10,158)=2.89<br>Time x stim<br>F(118,9322)=20.98<br>Geno x time x stim<br>F(590,9322)=10.88 | Genotype P=0.034<br>Time P=2.9e-150<br>Geno x time P=1.4e-14<br>Stim P=1.7e-14<br>Geno x stim P=0.0036<br>Time x stim P=2.9e-34<br>Geno x time x stim<br>P=9.8e-60 | Difference with D1/D2 during the first sond after locomotion onset<br>D1: p<0.0001<br>D2: p=0.014<br>D1 + D1/D2: p<0.0001<br>D2 + D1/D2: p=0.265<br>D1/D2$^{ctl}$: p<0.0001 |
| Extended<br>Data Figure<br>6a | D1/D2: 21 mice<br>D2: 14 mice<br>D2+D1/D2: 14 mice | 2-way ANOVA<br>with one factor<br>replication | Stim x Genotype<br>F(6,85)=32.33<br>Stim | Stim x Genotype<br>P<0.0001<br>Stim | Pre vs LaserON :<br>D1/D2: P<0.0001<br>D2: P=0.0002 |

| | | | | | |
|---|---|---|---|---|---|
| | D1: 18 mice<br>D1+D1/D2: 9 mice<br>D1/D2$^{ctl}$: 11 mice | | F(1,85)=7.558<br>Genotype<br>F(6,85)=21.89 | P=0.0073<br>Genotype<br>P<0.0001 | D2+D1/D2: P<0.0001<br>D1: P<0.0001<br>D1+D1/D2: P=0.0095<br>D1/D2$^{ctl}$: P>0.9999 |
| Figure 4d<br>Extended<br>Data Figure<br>8a<br>L>L | D1/D2: 21 mice<br>D2: 15 mice<br>D2+D1/D2: 11 mice<br>D1: 16 mice<br>D1+D1/D2: 10 mice<br>D1/D2$^{ctl}$: 11 mice | 2-way ANOVA<br>with one factor<br>replication | Stim x Genotype<br>F(15,255)=1.674<br>Stim<br>F(3,255)=18.73<br>Genotype<br>F(5,75)=2.214 | Stim x Genotype<br>P=0.1023<br>Stim<br>P<0.0001<br>Genotype<br>P=0.0616 | Pre vs LaserON 0-5 s \| LaserON 10-15 s \|<br>LaserON 20-25 s :<br>D1/D2: P=0.2943 \| P=0.6600 \| P=0.9744<br>D1: P=0.910 \| P=0.0115 \| P=0.1322<br>D2: P=0.0367 \| P=0.0354 \| P=0.0056<br>D1+D1/D2: P=0.8855 \| P=0.0735 \| P=0.0004<br>D2+D1/D2: P=0.2476 \| P=0.7541 \| P=0.0679<br>D1/D2$^{ctl}$: P=0.9996 \| P=0.9869 \| P=0.9177 |
| Figure 4d<br>Extended<br>Data Figure<br>8a<br>L>S | D1/D2: 21 mice<br>D2: 15 mice<br>D2+D1/D2: 11 mice<br>D1: 16 mice<br>D1+D1/D2: 10 mice<br>D1/D2$^{ctl}$: 11 mice | 2-way ANOVA<br>with one factor<br>replication | Stim x Genotype<br>F(15,255)=4.476<br>Stim<br>F(3,255)=20.34<br>Genotype<br>F(5,75)=6.129 | Stim x Genotype<br>P<0.0001<br>Stim<br>P<0.0001<br>Genotype<br>P<0.0001 | Pre vs LaserON 0-5 s \| LaserON 10-15 s \|<br>LaserON 20-25 s :<br>D1/D2: P<0.0001\| P<0.0001 \| P<0.0001<br>D1: P=0.2285 \| P=0.05020 \| P=0.5209<br>D2: P=0.0004 \| P=0.0001 \| P<0.0001<br>D1+D1/D2: P=0.7556 \| P=0.3196 \| P=0.0040<br>D2+D1/D2: P=0.0561 \| P=0.0001 \| P<0.0001<br>D1/D2$^{ctl}$: P=0.9614 \| P=1 \| P=0.8956 |
| Figure 4d<br>Extended<br>Data Figure<br>8a<br>L>I | D1/D2: 21 mice<br>D2: 15 mice<br>D2+D1/D2: 11 mice<br>D1: 16 mice<br>D1+D1/D2: 10 mice<br>D1/D2$^{ctl}$: 11 mice | 2-way ANOVA<br>with one factor<br>replication | Stim x Genotype<br>F(15,255)=6.984<br>Stim<br>F(3,255)=30.64<br>Genotype<br>F(5,75)=20.33 | Stim x Genotype<br>P<0.0001<br>Stim<br>P<0.0001<br>Genotype<br>P<0.0001 | Pre vs LaserON 0-5 s \| LaserON 10-15 s \|<br>LaserON 20-25 s :<br>D1/D2: P<0.0001 \| P<0.0001 \| P<0.0001<br>D1: P=0.9089 \| P=0.3982 \| P=0.2593<br>D2: P<0.0001 \| P<0.0001 \| P<0.0001<br>D1+D1/D2: P=0.9977 \| P=0.0309 \| P=0.0206<br>D2+D1/D2: P<0.0001 \| P<0.0001 \| P<0.0001<br>D1/D2$^{ctl}$: P=0.9933 \| P=0.9208 \| P=1 |
| Figure 4d<br>Extended<br>Data Figure<br>8a<br>S>L | D1/D2: 21 mice<br>D2: 15 mice<br>D2+D1/D2: 11 mice<br>D1: 16 mice<br>D1+D1/D2: 10 mice<br>D1/D2$^{ctl}$: 11 mice | 2-way ANOVA<br>with one factor<br>replication | Stim x Genotype<br>F(15,255)=1.018<br>Stim<br>F(3,255)=4.002<br>Genotype<br>F(5,75)=4.786 | Stim x Genotype<br>P=0.4253<br>Stim<br>P=0.0084<br>Genotype<br>P=0.0008 | Pre vs LaserON 0-5 s \| LaserON 10-15 s \|<br>LaserON 20-25 s :<br>D1/D2: P=0.9520 \| P=0.0532 \| P=0.0007<br>D1: P=0.7129 \| P=0.5424 \| P=0.6642<br>D2: P=0.2460 \| P=0.0032 \| P<0.0001<br>D1+D1/D2: P=0.7659 \| P=0.9999 \| P=0.3331<br>D2+D1/D2: P=0.4632 \| P=0.0294 \| P<0.0001<br>D1/D2$^{ctl}$: P=0.9871 \| P=0.9955 \| P=0.8663 |
| Figure 4d<br>Extended<br>Data Figure<br>8a<br>S>S | D1/D2: 21 mice<br>D2: 15 mice<br>D2+D1/D2: 11 mice<br>D1: 16 mice<br>D1+D1/D2: 10 mice<br>D1/D2$^{ctl}$: 11 mice | 2-way ANOVA<br>with one factor<br>replication | Stim x Genotype<br>F(15,255)=5.788<br>Stim<br>F(3,255)=8.461<br>Genotype<br>F(5,75)=19.53 | Stim x Genotype<br>P<0.0001<br>Stim<br>P=0.0004<br>Genotype<br>P<0.0001 | Pre vs LaserON 0-5 s \| LaserON 10-15 s \|<br>LaserON 20-25 s :<br>D1/D2: P=0.0005 \| P<0.0001 \| P<0.0001<br>D1: P=0.1987 \| P=0.0164 \| P=0.0115<br>D2: P<0.0001 \| P<0.0001 \| P<0.0001<br>D1+D1/D2: P=0.3129 \| P=0.4336 \| P=0.9771<br>D2+D1/D2: P=0.0049 \| P<0.0001 \| P<0.0001<br>D1/D2$^{ctl}$: P=0.9955 \| P=0.9999 \| P=0.4354 |
| Figure 4d<br>Extended<br>Data Figure<br>8a<br>S>I | D1/D2: 21 mice<br>D2: 15 mice<br>D2+D1/D2: 11 mice<br>D1: 16 mice<br>D1+D1/D2: 10 mice<br>D1/D2$^{ctl}$: 11 mice | 2-way ANOVA<br>with one factor<br>replication | Stim x Genotype<br>F(15,255)=4.957<br>Stim<br>F(3,255)=9.681<br>Genotype<br>F(5,75)=21.79 | Stim x Genotype<br>P<0.0001<br>Stim<br>P<0.0001<br>Genotype<br>P<0.0001 | Pre vs LaserON 0-5 s \| LaserON 10-15 s \|<br>LaserON 20-25 s :<br>D1/D2: P<0.0001 \| P=0.0040 \| P=0.0085<br>D1: P=0.6220 \| P=0.0879 \| P=0.0364<br>D2: P<0.0001 \| P=0.0016 \| P=0.7828<br>D1+D1/D2: P=0.7695 \| P=0.1503 \| P=0.2255<br>D2+D1/D2: P=0.0329 \| P=0.0006 \| P=0.6295<br>D1/D2$^{ctl}$: P=0.9950 \| P=0.9816 \| P=0.7644 |
| Figure 4d<br>Extended<br>Data Figure<br>8a<br>I>L | D1/D2: 21 mice<br>D2: 15 mice<br>D2+D1/D2: 11 mice<br>D1: 16 mice<br>D1+D1/D2: 10 mice<br>D1/D2$^{ctl}$: 11 mice | 2-way ANOVA<br>with one factor<br>replication | Stim x Genotype<br>F(15,255)=2.234<br>Stim<br>F(3,255)=1.530<br>Genotype<br>F(5,75)=2.709 | Stim x Genotype<br>P=0.0116<br>Stim<br>P=0.2148<br>Genotype<br>P=0.0291 | Pre vs LaserON 0-5 s \| LaserON 10-15 s \|<br>LaserON 20-25 s :<br>D1/D2: P=0.1751 \| P=0.0498 \| P=0.0005<br>D1: P=0.0213 \| P=0.1227 \| P=0.6219<br>D2: P=0.7753 \| P=0.3164 \| P=0.0050<br>D1+D1/D2: P=0.0486 \| P=0.0395 \| P=0.6824<br>D2+D1/D2: P=0.9993 \| P=0.7852 \| P=0.4410<br>D1/D2$^{ctl}$: P=0.3570 \| P=0.7449 \| P=0.9988 |
| Figure 4d<br>Extended<br>Data Figure<br>8a<br>I>S | D1/D2: 21 mice<br>D2: 15 mice<br>D2+D1/D2: 11 mice<br>D1: 16 mice<br>D1+D1/D2: 10 mice<br>D1/D2$^{ctl}$: 11 mice | 2-way ANOVA<br>with one factor<br>replication | Stim x Genotype<br>F(15,255)=11.95<br>Stim<br>F(3,255)=12.95<br>Genotype<br>F(5,75)=38.33 | Stim x Genotype<br>P<0.0001<br>Stim<br>P<0.0001<br>Genotype<br>P<0.0001 | Pre vs LaserON 0-5 s \| LaserON 10-15 s \|<br>LaserON 20-25 s :<br>D1/D2: P<0.0001 \| P<0.0001 \| P=0.0123<br>D1: P=0.0009 \| P=0.9115 \| P=0.8802<br>D2: P<0.0001 \| P<0.0001 \| P=0.0212<br>D1+D1/D2: P=0.2277 \| P<0.0001 \| P<0.0001<br>D2+D1/D2: P<0.0001 \| P<0.0001 \| P=0.0407<br>D1/D2$^{ctl}$: P=0.9978 \| P=0.4546 \| P=1 |
| Figure 4d<br>Extended<br>Data Figure<br>8a<br>I>I | D1/D2: 21 mice<br>D2: 15 mice<br>D2+D1/D2: 11 mice<br>D1: 16 mice<br>D1+D1/D2: 10 mice<br>D1/D2$^{ctl}$: 11 mice | 2-way ANOVA<br>with one factor<br>replication | Stim x Genotype<br>F(15,255)=10.51<br>Stim<br>F(3,255)=18.17<br>Genotype<br>F(5,75)=29.08 | Stim x Genotype<br>P<0.0001<br>Stim<br>P<0.0001<br>Genotype<br>P<0.0001 | Pre vs LaserON 0-5 s \| LaserON 10-15 s \|<br>LaserON 20-25 s :<br>D1/D2: P=0.0049 \| P=0.0021 \| P=0.5491<br>D1: P=0.0258 \| P=0.9926 \| P=0.8267<br>D2: P=0.0016 \| P=0.0017 \| P=0.4943<br>D1+D1/D2: P=0.8831 \| P<0.0001 \| P<0.0001<br>D2+D1/D2: P=0.0018 \| P=0.0064 \| P=0.0276<br>D1/D2$^{ctl}$: P=0.8893 \| P=0.9597 \| P=1 |
| Figure 5c | D1/D2: 11 cells<br>D1: 9 cells<br>D2: 8 cells | 2-way ANOVA<br>with one factor<br>replication | Geno F(2,25)=61.9<br>Drug F(1,25)=37.5<br>Geno x drug F(2,25)=165.2 | Geno P=0.054<br>Drug P=0.032<br>Geno x drug P=2.3e-6 | D1 base vs drug: P=0.008<br>D2 base vs drug: P=0.005<br>D1/D2 base vs drug: P=0.0002<br>Drug effect D1 vs D2: P=9.5e-5<br>Drug effect D1 vs D1/D2: P=0.96<br>Drug effect D2 vs D1/D2: P=2.3e-5 |
| Figure 5d | D1/D2: 11 cells<br>D1: 9 cells<br>D2: 8 cells | 2-way ANOVA<br>with one factor<br>replication | Geno F(2,25)=2.9<br>Drug F(1,25)=19.9<br>Geno x drug F(2,25)=39.8 | Geno P=0.072<br>Drug P=0.0002<br>Geno x drug P=1.7e-8 | D1 base vs drug: P=6.8e-6<br>D2 base vs drug: P=0.02<br>D1/D2 base vs drug: P=0.08<br>Drug effect D1 vs D2: P=3.7e-7 |

| | | | | | |
|---|---|---|---|---|---|
| | | | | | Drug effect D1 vs D1/D2: P=1.8e-5<br>Drug effect D2 vs D1/D2: P=0.002 |
| Figure 5e | D1/D2: 10 cells<br>D1: 9 cells<br>D2: 7 cells | 2-way ANOVA with one factor replication | Geno F(2,23)=3.5<br>Drug F(1,23)=1.2<br>Geno x drug F(2,23)=8.22 | Geno P=0.046<br>Drug P=0.28<br>Geno x drug P=0.002 | D1 base vs drug: P=0.36<br>D2 base vs drug: P=0.08<br>D1/D2 base vs drug: P=0.005<br>Drug effect D1 vs D2: P=0.17<br>Drug effect D1 vs D1/D2: P=0.007<br>Drug effect D2 vs D1/D2: P=0.008 |
| Figure 5f | D1/D2: 12 cells<br>D1: 5 cells<br>D2: 6 cells | 2-way ANOVA with one factor replication | Geno F(2,20)=0.1<br>Drug F(1,20)=13.0<br>Geno x drug F(2,20)=18.4 | Geno P=0.90<br>Drug P=0.002<br>Geno x drug P=2.6e-5 | D1 base vs drug: P=0.0004<br>D2 base vs drug: P=0.026<br>D1/D2 base vs drug: P=0.007<br>Drug effect D1 vs D2: P=0.38<br>Drug effect D1 vs D1/D2: P=1.6e-6<br>Drug effect D2 vs D1/D2: P=0.0045 |
| Figure 5g | D1/D2: 12 cells<br>D1: 5 cells<br>D2: 6 cells | 2-way ANOVA with one factor replication | Geno F(2,20)=0.58<br>Drug F(1,20)=5.8<br>Geno x drug F(2,20)=6.3 | Geno P=0.57<br>Drug P=0.026<br>Geno x drug P=0.007 | D1 base vs drug: P=3.5e-6<br>D2 base vs drug: P=0.22<br>D1/D2 base vs drug: P=0.40<br>Drug effect D1 vs D2: P=0.006<br>Drug effect D1 vs D1/D2: P=0.0003<br>Drug effect D2 vs D1/D2: P=0.14 |
| Extended Data Figure 9c | D1/D2: 12 cells<br>D1: 9 cells<br>D2: 7 cells | 2-way ANOVA with one factor replication | Geno F(2,23)=2.80<br>Drug F(1,23)=0.05<br>Geno x drug F(2,23)=7.66 | Geno P=0.08<br>Drug P=0.82<br>Geno x drug P=0.0028 | D1 base vs drug: P=0.041<br>D2 base vs drug: P=0.021<br>D1/D2 base vs drug: P=0.48<br>Drug effect D1 vs D2: P=0.002<br>Drug effect D1 vs D1/D2: P=0.029<br>Drug effect D2 vs D1/D2: P=0.059 |
| Extended Data Figure 9d | D1/D2: 10 cells<br>D1: 9 cells<br>D2: 8 cells | 2-way ANOVA with one factor replication | Geno F(2,24)=0.38<br>Drug F(1,24)=13.4<br>Geno x drug F(2,23)=23.8 | Geno P=0.69<br>Drug P=0.0012<br>Geno x drug P=2.0e-6 | D1 base vs drug: P=0.0004<br>D2 base vs drug: P=0.032<br>D1/D2 base vs drug: P=0.0002<br>Drug effect D1 vs D2: P=5.7e-5<br>Drug effect D1 vs D1/D2: P=0.08<br>Drug effect D2 vs D1/D2: P=0.0005 |
| Extended Data Figure 9e | D1/D2: 11 cells<br>D1: 5 cells<br>D2: 8 cells | 3-way ANOVA with two factors replication | Geno F(2,21)=3.25<br>Frequency F(16,336)=96.0<br>Geno x freq F(32,336)=3.41<br>Drug F(1,21)=92;6<br>Geno x drug F(2,21)=163.4<br>Freq x drug F(16,336)=22.8<br>Geno x freq x drug F(32,336)=30.7 | Geno: P=0.059<br>Frequency: P<1e-12<br>Geno x freq: P=1e-8<br>Drug: P=3.8e-9<br>Geno x drug: P<1e-12<br>Freq x drug: P<1e-12<br>Geno x freq x drug: P<1e-12 | |
| Extended Data Figure 9f | D1/D2-SPNs<br>D1R antago: 5<br>D2R antago: 5 | 2-way ANOVA with one factor replication | Antago F(1,8)=1.38<br>Drug F(1,8)=8.43<br>Antago x drug F(1,8)=9.62 | Antago: P=0.27<br>Drug: P=0.020<br>Antago x drug: P=0.015 | D1R antago: P=0.87<br>D2R antago: P=0.021<br>Antago effect D1R vs D2R: P=0.016 |
| Extended Data Figure 9g | D1/D2-SPNs<br>D1R antago: 5<br>D2R antago: 5 | 2-way ANOVA with one factor replication | Antago F(1,8)=0.05<br>Drug F(1,8)=41.1<br>Antago x drug F(1,8)=8.94 | Antago: P=0.82<br>Drug: P=0.0002<br>Geno x drug: P=0.017 | D1R antago: P=0.12<br>D2R antago: P=0.0007<br>Antago effect D1R vs D2R: P=0.022 |
| Extended Data Figure 9h | D1/D2-SPNs<br>No antago: 11<br>D1R antago: 5<br>D2R antago: 5 | 2-way ANOVA with one factor replication | Antago F(2,18)=2.04<br>Drug F(1,18)=18.01<br>Antago x drug F(2,18)=7.32 | Antago: P=0.16<br>Drug: P=0.0005<br>Antago x drug: P=0.0047 | No antago base vs drug: P=0.0002<br>D1R antago base vs drug: P=0.37<br>D2R antago base vs drug: P=0.048<br>Drug effect no antago vs D1R antago: P=0.0003<br>Drug effect no antago vs D2R antago: P=0.67<br>Drug effect D1R antago vs D2R antago: P=0.029 |
| Extended Data Figure 9i | D1/D2-SPNs<br>No antago: 11<br>D1R antago: 5<br>D2R antago: 5 | 2-way ANOVA with one factor replication | Antago F(2,18)=0.97<br>Drug F(1,18)=0.81<br>Antago x drug F(2,18)=1.73 | Antago: P=0.40<br>Drug: P=0.38<br>Antago x drug: P=0.21 | No antago base vs drug: P=0.08<br>D1R antago base vs drug: P=0.09<br>D2R antago base vs drug: P=0.57<br>Drug effect no antago vs D1R antago: P=0.021<br>Drug effect no antago vs D2R antago: P=0.51<br>Drug effect D1R antago vs D2R antago: P=0.265 |
| Extended Data Figure 9j | D1/D2-SPNs<br>No antago: 10<br>D1R antago: 5<br>D2R antago: 5 | 2-way ANOVA with one factor replication | Antago F(2,17)=2.34<br>Drug F(1,17)=14.40<br>Antago x drug F(2,17)=2.47 | Antago: P=0.13<br>Drug: P=0.0015<br>Antago x drug: P=0.11 | No antago base vs drug: P=0.005<br>D1R antago base vs drug: P=0.208<br>D2R antago base vs drug: P=0.0298<br>Drug effect no antago vs D1R antago: P=0.018<br>Drug effect no antago vs D2R antago: P=0.29<br>Drug effect D1R antago vs D2R antago: P=0.087 |
| Extended Data Figure 9l | D1/D2-SPNs<br>No antago: 10<br>D1R antago: 5<br>D2R antago: 5 | 2-way ANOVA with one factor replication | Antago F(2,17)=0.45<br>Drug F(1,17)=0.04<br>Antago x drug F(2,17)=3.56 | Antago: P=0.64<br>Drug: P=0.83<br>Antago x drug: P=0.049 | No antago base vs drug: P=0.007<br>D1R antago base vs drug: P=0.41<br>D2R antago base vs drug: P=0.69<br>Drug effect no antago vs D1R antago: P=0.047<br>Drug effect no antago vs D2R antago: P=0.16<br>Drug effect D1R antago vs D2R antago: P=0.85 |
| Extended Data Figure 9m | D1/D2-SPNs<br>No antago: 10<br>D1R antago: 5<br>D2R antago: 5 | 2-way ANOVA with one factor replication | Antago F(2,17)=1.25<br>Drug F(1,17)=0.30<br>Antago x drug F(2,17)=1.15 | Antago: P=0.30<br>Drug: P=0.59<br>Antago x drug: P=0.34 | No antago base vs drug: P=0.39<br>D1R antago base vs drug: P=0.37<br>D2R antago base vs drug: P=0.20<br>Drug effect no antago vs D1R antago: P=0.21<br>Drug effect no antago vs D2R antago: P=0.13<br>Drug effect D1R antago vs D2R antago: P=0.96 |
| Figure 6c | $D1^{WT}$/D2: 12<br>$D1^{cKO}$/D2: 15 | Two tailed unpaired t test | Locomotion: t(25)=2.592<br>Small movements: t(25)=2.073<br>Immobility: t(25)=2.134 | | Locomotion: P=0.0157<br>Small movements: P=0.0487<br>Immobility: P=0.0428 |
| Figure 6d | NaCl :<br>$D1^{WT}$/D2: 18<br>$D1^{cKO}$/D2: 18<br>1 mg/kg :<br>$D1^{WT}$/D2: 18 | 3-way ANOVA with one factor (time) replication | Genotype F(1,122)=38.34<br>Dose F(4,122)=27.14<br>Geno x Dose F(4,122)=2.56<br>Time F(23,2806)=38.83 | Genotype P=5.6e-9<br>Dose P=7.2e-17<br>Geno x Dose P=0.0412<br>Time P=3.6e-30<br>Geno x Time P=3.5e-9 | $D1^{WT}$/D2 vs $D1^{cKO}$/D2 temporal evolution of distance<br>NaCl: P=0.037<br>1 mg/kg: P=0.0173<br>5 mg/kg: P=0.0134 |

| | | | | | |
|---|---|---|---|---|---|
| | D1$^{cKO}$/D2: 18<br>5 mg/kg :<br>D1$^{WT}$/D2: 14<br>D1$^{cKO}$/D2: 14<br>10 mg/kg:<br>D1$^{WT}$/D2: 15<br>D1$^{cKO}$/D2: 15<br>20 mg/kg :<br>D1$^{WT}$/D2: 14<br>D1$^{cKO}$/D2: 13 | | Geno x Time<br>F(23,2806)=11.32<br>Dose x Time<br>F(92,2806)=5.75<br>Geno x Dose x Time<br>F(92,2806)=3.57 | Dose x Time P=3.7e-12<br>Geno x Dose x Time<br>P=2.0e-6 | 10 mg/kg: P<0.0001<br>20 mg/kg: P<0.0001<br>D1$^{WT}$/D2 vs D1$^{cKO}$/D2 total distance<br>NaCl: P=0.0115<br>1 mg/kg: P=0.0144<br>5 mg/kg: P=0.0406<br>10 mg/kg: P<0.0001<br>20 mg/kg: P=0.1713<br>total distance vs NaCl<br>1 mg/kg: D1$^{WT}$/D2 P=0.999 D1$^{cKO}$/D2 P=0.654<br>5 mg/kg: D1$^{WT}$/D2 P=0.957 D1$^{cKO}$/D2 P=0.504<br>10 mg/kg: D1$^{WT}$/D2 P=0.0281 D1$^{cKO}$/D2 P<0.0001<br>20 mg/kg: D1$^{WT}$/D2 P<0.0001 D1$^{cKO}$/D2 P<0.0001 |
| Figure 6e<br>Extended Data Figure 7c | D1$^{WT}$/D2: 17<br>D1$^{cKO}$/D2: 16 | 3-way ANOVA with two factors (time, days) replication | Genotype F(1,31)=9.63<br>Time F(26,806)=37.47<br>Geno x Time<br>F(26,806)=8.36<br>Days F(6,186)=1.32<br>Geno x Days F(6,186)=3.79<br>Days x Time<br>F(156,4836)=4.32<br>Geno x Days x Time<br>F(156,4836)=1.45 | Genotype P=0.0004<br>Time P<0.0001<br>Geno x Time P=0.0002<br>Days P=0.271<br>Geno x Days P=0.0121<br>Days x Time P<0.0001<br>Geno x Days x Time<br>P=0.125 | D1$^{WT}$/D2 vs D1$^{cKO}$/D2 temporal evolution of distance<br>Day 1: P<0.0001<br>Day 2: P<0.0001<br>Day 3: P<0.0001<br>Day 4: P<0.0001<br>Day 5: P<0.0001<br>Day 6: P<0.0001<br>Day 14: P<0.0001<br>effect of days<br>D1$^{WT}$/D2: P<0.0001<br>D1$^{cKO}$/D2 P=0.387<br>total distance vs Day 1<br>Day 2: D1$^{WT}$/D2 P=0.5391 D1$^{cKO}$/D2 P=0.632<br>Day 3: D1$^{WT}$/D2 P=0.0101 D1$^{cKO}$/D2 P=0.993<br>Day 4: D1$^{WT}$/D2 P=0.0043 D1$^{cKO}$/D2 P=0.999<br>Day 5: D1$^{WT}$/D2 P<0.0001 D1$^{cKO}$/D2 P=0.748<br>Day 6: D1$^{WT}$/D2 P<0.0001 D1$^{cKO}$/D2 P=0.879<br>Day 17: D1$^{WT}$/D2 P=0.0005 D1$^{cKO}$/D2 P=0.908 |
| Figure 6f | D1$^{WT}$/D2: 12<br>D1$^{cKO}$/D2: 9 | 2-way ANOVA with one factor replication | Drug dose x Genotype<br>F(1,19)=1.10<br>Drug F(1,19)=31.03<br>Genotype F(1,19)=1.74 | Drug dose x Genotype<br>P=0.308<br>Drug Dose P<0.0001<br>Genotype P=0.203 | D1$^{WT}$/D2 vs D1$^{cKO}$/D2<br>NaCl: P=0.830<br>10 mg/kg: P=0.297 |
| Extended Data Figure 7a | D1$^{WT}$/D2: 12<br>D1$^{cKO}$/D2: 15 | Two tailed unpaired t test | Velocity mean (black):<br>t(25)=2.539<br>Velocity max (green):<br>t(25)=2.179<br>Locomotion bout number:<br>t(25)=0.365<br>Locomotion bout mean duration: t(25)=2.53<br>Small movements bout number: t(25)=1.929<br>Small movements bout mean duration: t(25)=0.979<br>Immobility bout number:<br>t(25)=2.744<br>Immobility bout mean duration: t(25)=0.535 | | Velocity mean: P=0.0177<br>Velocity max: P=0.0390<br>Locomotion bout number: P=0.781<br>Locomotion bout mean duration P=0.0181<br>Small movements bout number: P=0.0651<br>Small movements bout mean duration: P=0.336<br>Immobility bout number: P=0.011<br>Immobility bout mean duration: P=0.598 |
| Extended Data Figure 7b | NaCl :<br>D1$^{WT}$/D2: 8<br>D1$^{cKO}$/D2: 6<br>1 mg/kg :<br>D1$^{WT}$/D2: 8<br>D1$^{cKO}$/D2: 8<br>3 mg/kg :<br>D1$^{WT}$/D2: 7<br>D1$^{cKO}$/D2: 7 | 3-way ANOVA with one factor replication | Genotype F(1,34)=8.03<br>Dose F(2,34)=35.83<br>Geno x Dose F(2,342)=0.39<br>Time F(23,782)=20.02<br>Geno x Time<br>F(23,782)=3.12<br>Dose x Time<br>F(46,782)=9.81<br>Geno x Dose x Time<br>F(46,782)=1.75 | Genotype P=0.0077<br>Dose P=4.3e-9<br>Geno x Dose P=0.681<br>Time P=3.7e-15<br>Geno x Time P=0.0113<br>Dose x Time P=1.9e-12<br>Geno x Dose x Time<br>P=0.0763 | D1$^{WT}$/D2 vs D1$^{cKO}$/D2 temporal evolution of distance<br>NaCl: P=0.009<br>1 mg/kg: P<0.0001<br>3 mg/kg: P=0.947<br>D1$^{WT}$/D2 vs D1$^{cKO}$/D2 total distance over 60 min<br>NaCl: P<0.0001<br>1 mg/kg: P=0.015<br>3 mg/kg: P=0.710 |
